# Influence of phosphate activation chemistry on the selection of the primordial genetic alphabet

**DOI:** 10.64898/2025.12.23.696203

**Authors:** Filip Bošković, Jian Zhang, Alok Apan Swatiputra, Jack W. Szostak

**Author notes:** **Corresponding Author** Jack W. Szostak – Howard Hughes Medical Institute, Department of Chemistry, The University of Chicago, Chicago, Illinois, United States.

## Abstract

RNA copying under mild conditions compatible with protocell integrity requires the input of chemical energy to drive the synthesis of activated nucleotides such as phosphorimidazolides. Recently, two potentially prebiotic classes of phosphate-activating agents have been explored, one based on isonitrile–aldehyde chemistry, the other on imine diimidazole (IDI)-N-cyanoimidazole (NCI) chemistry. Because such highly electrophilic activating agents may lead to undesirable nucleotide modifications, we have examined the reaction of both types of activating agents with the canonical ribonucleotides A, U, C, and G, and the potentially primordial nucleotides 2-thio-C (s^2^C), 2-thio-U (s^2^U), and inosine (I). We find that the isonitrile–aldehyde system shows minimal hydroxyl modification but does modify the nucleobases of U, G, s^2^U, and I. Except for guanosine, these modifications are readily reversible. In contrast, IDI-NCI systems acylate ribonucleotide hydroxyls while modifying nucleobases only transiently; mildly acidic pH suppresses undesired modifications. Both classes of activating agents modify 2-thiopyrimidines on the sulfur, with the isonitrile–aldehyde reaction promoting desulfurization and thus conversion to the canonical pyrimidines. To evaluate compatibility with model protocells, we tested the effects of activation chemistry on fatty acid vesicles and found that protocell integrity was preserved at moderate reagent concentrations. Our findings show that the potentially primordial s^2^U, s^2^C, and I nucleotides are more sensitive to modification than the canonical U, C, and G nucleotides, potentially contributing to the chemical selection of the early genetic alphabet.

## INTRODUCTION

The emergence of RNA as both a genetic polymer and a catalyst was a central step in the transition from prebiotic chemistry to biology^1,2^. For primitive cells to sustain nonenzymatic RNA replication, two requirements must have been met: a source of activated nucleotides and short oligonucleotides capable of undergoing template-directed polymerization, and a physical compartment capable of retaining genetic material^3–5^. Fatty acid vesicles provide a compelling model for early protocells, yet their chemical fragility imposes strict limits on their ambient chemical environment^6–8^.

Nonenzymatic RNA copying relies on activated forms of nucleotides, typically phosphorimidazolides^9–11^. Prebiotic scenarios therefore require *in situ* phosphate activation to regenerate these high-energy intermediates from unactivated monomers^10–17^. However, activation chemistry agents that act directly on complex mixtures of nucleotides and short oligomers and may also lead to deleterious nucleotide modifications^18,19^. This consideration is particularly important for nucleotides such as the 2-thiopyrimidines, which have been proposed as prebiotic precursors of canonical pyrimidines, and for inosine, which is a chemically plausible surrogate for guanosine in early genetic systems^20–23^. As a result, the specific activation chemistry available on the early Earth may have influenced which nucleotides persisted long enough to participate in replication, thereby biasing the composition of the primordial genetic alphabet.

The chemical requirements for RNA replication must also be compatible with protocellular integrity^7,24,25^. Fatty acid vesicles can support growth, division, and RNA encapsulation, but only within a narrow chemical space^25–28^. Activation reagents that disrupt vesicle structure by modifying lipids would lead to loss of genetic material and prevent the emergence of Darwinian evolution. Consequently, the reactions that enabled nucleotide activation and RNA copying must have preserved the integrity of protocell membranes^29^. This dual requirement creates a shared chemical selection pressure on both nucleotide structures and membrane composition.

Herein, we examine how two classes of phosphate-activating agents including isonitrile-based activation^16,30,31^ and N-cyanoimidazole (NCI)–imine diimidazole (IDI)-based activation^32–35^ influence nucleotide modification and protocell integrity. Across both classes, mildly acidic conditions promote selective phosphate activation while limiting undesired nucleotide modification. Further, we find that 2-thiopyrimidines undergo selective and condition-dependent transformation into canonical pyrimidines with isonitrile–aldehyde chemistry. We have also evaluated the stability of fatty acid vesicles in the presence of phosphate activation agents, finding that at moderate concentrations these agents do not affect protocell integrity. However, higher concentrations led to irreversible membrane disruption and loss of encapsulated content. Together, our findings provide constraints on how phosphate activation chemistry and environmental conditions may have contributed to the selection of the early genetic alphabet and the nature of early protocell membranes. Our work helps to define the chemical and physical constraints imposed by prebiotic activation chemistry and illustrates how such constraints could have shaped nucleotide stability, protocell viability, and ultimately the composition of the primordial genetic alphabet.

## RESULTS

### Extent and Reversibility of Nucleobase Modification by Isonitrile-based Phosphate Activation Chemistry

Isonitriles together with aldehydes can support phosphate activation under conditions suitable for nonenzymatic RNA polymerization, but may also introduce nucleobase modifications (**Figure 1**). These arise from nucleophilic attack by the deprotonated N3 of pyrimidines or N1 of purines on the nitrilium intermediate formed from an isonitrile and an aldehyde, yielding imidoyl adducts at the base-pairing interface (**Figure 1a**). To investigate the scope of these nucleobase reactions, we examined a panel of canonical and noncanonical nucleotides, including uridine (U), 2-thiouridine (s^2^U), guanosine (G), inosine (I), and xanthosine (X), characterized by a range of amide pKa values (**Figure 1b**)^36–38^. We expected that lower pKa values would favor deprotonation and lead to enhanced modification^39^. ^1^H NMR analysis revealed that nucleotides with lower pKa values exhibited higher levels of modification with methyl isonitrile (MeNC) and 4-pentenal (**Figure S1– S6**). In the presence of 200 mM activating agents at pH 8, U and I displayed the highest degree of imidoyl modification, followed by G and X (**Figure 1c**). Gel electrophoresis of 10-mer RNA oligomers confirmed these trends, showing that oligonucleotides containing U, I, or G were substantially modified (**Figure 1d**; **Figure S7**). To assess the pH dependence of this reactivity, we examined uridine 5′-monophosphate (UMP) and inosine 5′-monophosphate (IMP) from pH 6–9 and observed a marked reduction in modification at lower pH, consistent with the requirement for deprotonation (**Figure 1e**; **Figure S8** and **Figure S9**).

**Figure 1.**
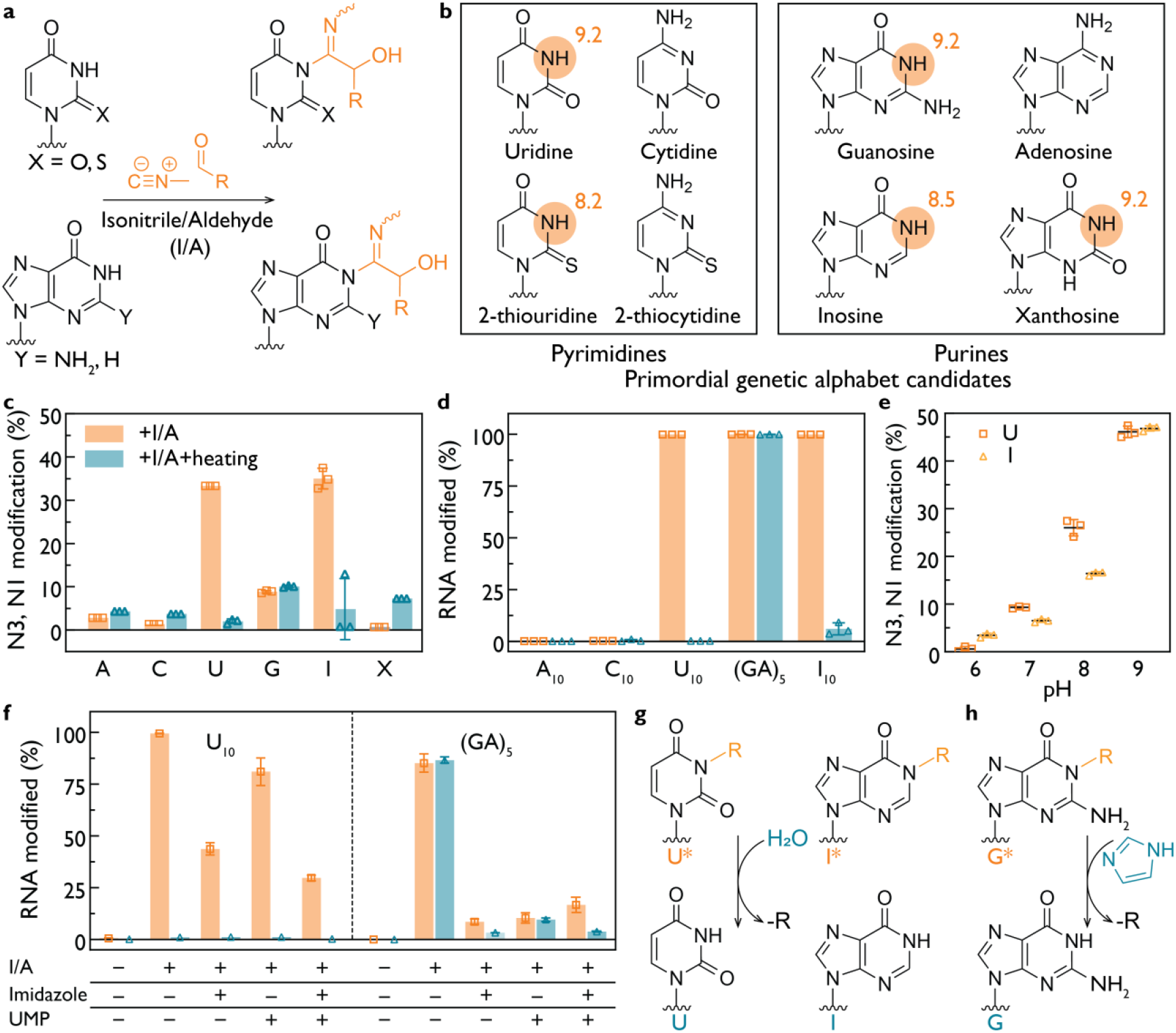
Nucleotide modification by isonitrile–aldehyde chemistry. **(a)** Products of nucleotide modification by isonitrile–aldehyde activation chemistry. The deprotonated N3 of pyrimidines or N1 of purines attacks the nitrilium ion generated from an isonitrile and an aldehyde (I/A), forming an imidoyl adduct on the nucleobase. **(b)** Structures of canonical and noncanonical nucleotides relevant to the primordial genetic alphabet. Reactive positions include N3 of uridine, 2 thiouridine (s^2^U), and N1 of purines, including guanosine (G) and inosine (I). pKa values of the protonated amide groups are indicated. **(c)** Modification of nucleoside for X or nucleotides (A, C, U, G, I) incubated with MeNC and 4-pentenal before and after heating at 95 °C, 15 min, determined by ^1^H NMR. U, I and G exhibit imidoyl modifications whose abundance correlates with the pKa of the protonated site. Heating reverses the modifications of U and I but not of G. **(d)** Electrophoretic mobility shift assays (EMSA) of 3′-Cy3-labeled RNA oligomers (C_10_, U_10_, (GA)_5_, and I_10_ as indicated) treated with isonitrile–aldehyde activation chemistry. Oligomers containing U, I and G are fully modified. Heating (95 °C, 15 min) restores native U and I but not G. **(e)** pH dependent modification of U and I. Higher pH increases the extent of N3 imidoyl modification of U and N1 imidoyl modification of I. **(f)** Suppression of modification in 3′-Cy3-labeled U_10_ and (GA)_5_ by imidazole (200 mM) and UMP (25 mM). Imidazole and UMP together have an additive effect. Imidazole removes the G modification in (GA)_5_. **(g)** Proposed mechanism of reversal of N3 imidoyl U and N1 imidoyl I modifications. **(h)** Proposed mechanism of reversal of G imidoyl modification by imidazole. Reaction conditions for all panels: 25 mM nucleotides or nucleosides or 1 µM RNA oligomer (3′-Cy3-labeled), 200 mM HEPES 8.0 (or at the indicated pH), 200 mM MeNC and 4-pentenal, 24 hours at 18 °C.

While the rate and extent of RNA modification is important, understanding the chemical stability of the resulting modifications is also critical. We therefore investigated the thermal lability of the RNA modifications induced by isonitrile activation chemistry. Heating the modified mononucleotides to 95 °C led to complete hydrolysis of adducts formed on U and I (**Figure 1c–d**; **Figure S10** and **Figure S11**). In contrast, G modifications remained stable in both monomeric and oligonucleotide contexts (**Figure 1c** and **Figure 1d**, respectively; **Figures S12**). Mass spectrometry identified an additional adduct at the N1 position of guanosine, with increased prominence following thermal treatment (**Figure S13**). The fact that N1 is indeed modified was further confirmed by the lack of reactivity of N1-methylguanosine (**Figure S14**) and the presence of the N1 modification in N^2^,N^2^-dimethylguanosine (**Figure S15**), demonstrating that the exocyclic amine is not participating in the reaction.

Next, we examined whether variations on the phosphate activation conditions, including the presence of 200 mM imidazole and/or 25 mM UMP, could limit the extent of the modifications of RNA U_10_ and (GA)_5_ (**Figure 1f**; **Figures S16–S17**). EMSA showed that for U_10_, both imidazole and UMP reduce the extent of modification, with an additive effect when combined. In the case of (GA)_5_, the otherwise thermally stable G modification was decreased in the presence of imidazole. We suggest that these effects may be due to a combination of preferential scavenging of the reactive nitrilium intermediate by imidazole and phosphate^40^, favoring phosphate activation over nucleobase modification (**Figure S18**), and imidazole mediated deprotection of the modified G.

Taken together, these findings demonstrate that nucleobase modification in the presence of isonitrile–aldehyde phosphate activation chemistry depends on both the pKa and structure of the nucleobase, and is less prominent under mildly acidic conditions. U and I adducts are reversible modifications that are removed by hydrolysis (**Figure 1g**), whereas the G adduct is more stable but can be removed by heating in the presence of imidazole (**Figure 1h**). The reversibility of these modifications supports the possibility that isonitrile activation chemistry would have been compatible with RNA replication using the canonical ribonucleotides.

### RNA Modification by Acylimidazole Activation Chemistry

Acylimidazoles have been evaluated for phosphate activation, but their potential to induce nucleotide modifications has not been systematically explored (**Figure 2a**). We focused on the IDI–NCI activation system because of its known use for phosphate activation and for nonenzymatic ligation^32,35^. Using EMSA and NMR spectroscopy, we found that acylimidazoles primarily acylate the 2′,3′-diol termini of RNA, internal 2′-hydroxyls, and the 3′-hydroxyl group of DNA, producing a distribution of esterified species (**Figure 2b**; **Figures S19–23**).

**Figure 2.**
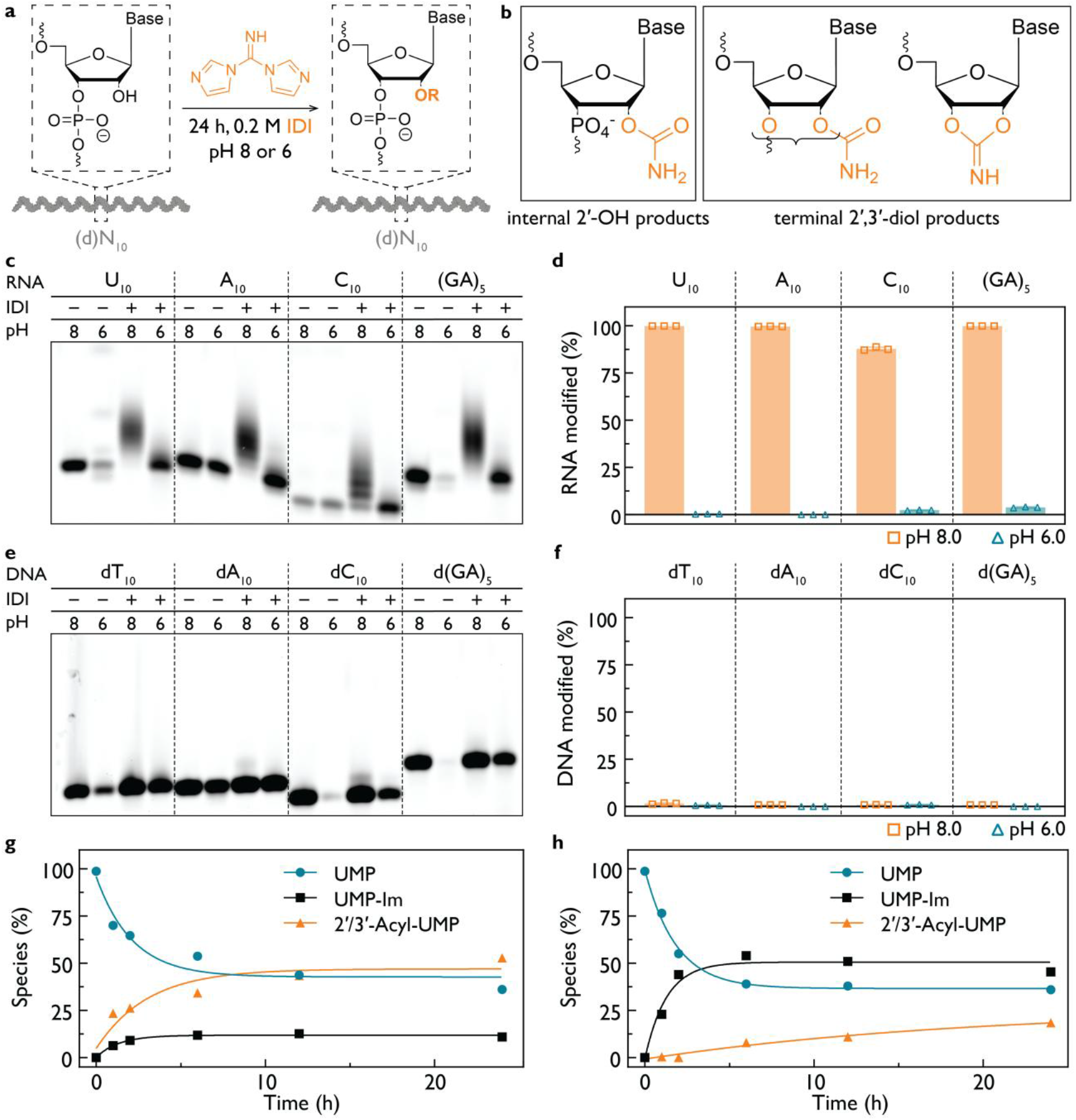
Acylimidazoles acylate oligonucleotide hydroxyl groups in a pH-dependent manner. **(a)** Schematic of the gel-based assay used to monitor chemical modification of RNA and DNA oligonucleotides. **(b)** Possible products of IDI acylation, including internal 2′-OH acylation or terminal acylation of the 2′,3′-diol for RNA, and 3′-OH acylation for DNA. **(c)** EMSA of 3′-Cy3-labeled RNA oligonucleotides treated with IDI at pH 8 and pH 6 and **(d)** gel quantification of modification. RNA is strongly acylated at pH 8, whereas acylation is suppressed at pH 6. **(e)** EMSA analysis of 3′-Cy3-labeled DNA oligonucleotides lacking 2′-OH groups at pH 8 and pH 6 and **(f)** gel quantification. DNA shows no detectable modification even at pH 8, consistent with minimal nucleobase reactivity of IDI. **(g)** Time course of reaction of 5′-UMP with IDI at pH 8.0 as monitored by ^31^P NMR. Curves show fit to a single step kinetic model. Unmodified UMP (turquoise circles), UMP-Im (black rectangles), acyl-UMP modifications (orange triangles). Observed rates: 0.47 h^−1^ for UMP consumption, 0.74 h^−1^ for UMP-Im formation, and 0.33 h^−1^ for acyl-UMP formation. **(h)** As in **g** but at pH 6.0, where UMP-Im formation dominates and acyl-UMP formation is strongly disfavored. Observed rates: 0.55 h^−1^ for UMP consumption, 0.79 h^−1^ for UMP-Im formation, and 0.05 h^−1^ for acyl-UMP formation. Reaction conditions for all panels: 25 mM nucleotides or 1 µM RNA/DNA oligomers, 200 mM HEPES 8.0 or 200 mM HEPES pH 6.0, 24 hours at 18 °C.

To assess the susceptibility of RNA to modification, we examined A_10_, C_10_, U_10_, and (GA)_5_ in the presence of 200 mM IDI at pH 8.0 ± 0.2 or pH 6.0 ± 0.35 for 24 h (**Figure 2c**). RNA modification was extensive at pH 8 but strongly reduced at pH 6.0 (**Figure 2d**; **Figures S23–S25**), consistent with hydroxyl pKa driving nucleophilic attack. We next asked whether these acylation modifications could be reversed or mitigated after formation. Heating modified RNA oligonucleotides at 95 °C for 30 min did not lead to a significant reduction in modification levels, indicating that the acylated species are thermally stable under these conditions (**Figures S25–S26**). Because duplex formation can reduce the accessibility of internal 2′-hydroxyl groups^19^, we examined reactions performed in the presence of complementary RNA strands and observed a partial reduction in acylation upon duplex formation (**Figure S27**). In contrast, addition of excess imidazole to reactions containing NCI did not measurably alter the extent of RNA modification (**Figure S28**), suggesting that once formed, hydroxyl acylation is not readily reversed by imidazole exchange.

To identify modification sites, we synthesized DNA analogs lacking internal 2′-hydroxyls, 5′-labeled with ATTO550 (**Figure 2e**). These oligonucleotides showed no detectable reaction with IDI, indicating that IDI selectively targets hydroxyl groups rather than nucleobases (**Figure 2f**; dN_10_-Cy3 data are shown in **Figures S29–S30**).

To determine whether hydroxyl acylation is a general property of acylimidazoles, we also evaluated NCI and thiocarbonyldiimidazole (TCDI) using NMR and EMSA (**Figures S30–S36**). All three reagents acylated available hydroxyl groups in both ribo- and deoxyribonucleotides. To isolate nucleobase-specific chemistry, we examined 5′-dimethylphosphate dideoxyuridine (mmddU), which lacks all free hydroxyls, and observed only low level transient N3 modification of uridine during the reaction course (**Figures S31–S36**). TCDI exhibited substantially reduced modification (**Figures S30**), likely due to rapid hydrolysis in water (**Figures S37–S38**). IDI, in contrast, showed high aqueous stability and minimal hydrolysis, consistent with its function as a persistent acylation reagent (**Figures S39–S40**).

We then quantified the competing formation of phosphorimidazolide (UMP-Im) and acylated/modified products using ^31^P and ^1^H NMR. For UMP at pH 8.0, acylation of the 2′,3′-diol dominated, with the observed rates of 0.47 h^−1^ (UMP consumption), 0.74 h^−1^ (UMP-Im formation), and 0.33 h^−1^ (acylated UMP formation) over 24 h (**Figure 2g**; **Figures S41–S42**). At pH 6.0, phosphate activation exceeded 50% conversion and hydroxyl acylation was strongly suppressed, with the observed rates of 0.55 h^−1^ (UMP), 0.79 h^−1^ (UMP-Im), and 0.05 h^−1^ (acylated UMP with PPi-UMP), respectively (**Figure 2h**). These data reflect strong pH control over chemoselectivity, favoring phosphate activation under mildly acidic conditions. AMP, CMP, and GMP rates displayed similar behavior (**Figure S43**). In reactions initiated but not actively maintained at pH 6.0, gradual release of imidazole increased the pH to ∼7.5, enabling late-stage hydroxyl acylation even in DNA oligonucleotides bearing only a terminal hydroxyl (**Figure S30**). This pH drift highlights the importance of buffered mildly acidic conditions for selective phosphate activation.

Given these observations, we next examined the kinetics and potential mechanism of phosphate activation with IDI and NCI with ^31^P NMR (**Figure S44**). We monitored phosphorimidazolide formation from UMP at controlled pH 6.0 over the course of the reaction to determine whether IDI and NCI operate through an imidazole-dependent or intramolecular pathway^32,41,42^. At pH 6.0, both reagents generated phosphorimidazolide at comparable rates (**Figure S44b**), indicating that free imidazole is unlikely to dominate the reaction under mildly acidic conditions. These observations are consistent with proposals that imidazole-activated phosphoryl transfer can proceed, at least in part, through an intramolecular collapse of the acylimidazole intermediate rather than exclusively through intermolecular imidazole exchange (**Figure S44c**)^41,43,44^. This interpretation is further supported by prior work showing that N-cyano-2-aminoimidazole at pH 5.5-6 yields the corresponding phosphorimidazolide without requiring external imidazole^45^. Together, these data suggest that an intramolecular pathway for phosphorimidazolide formation is chemically plausible, under mildly acidic conditions.

### Compatibility of 2-Thiopyrimidines with In Situ Phosphate Activation

2-thiocytidine (s^2^C) and 2-thiouridine (s^2^U) are attractive candidates for a primordial genetic alphabet because they form isoenergetic base pairs with inosine and adenosine, respectively, supporting more uniform nonenzymatic RNA copying^21,23^. To evaluate their compatibility with *in situ* phosphate-activation chemistry, we first examined their reactivity under isonitrile**–**aldehyde conditions (**Figure 3**).

**Figure 3.**
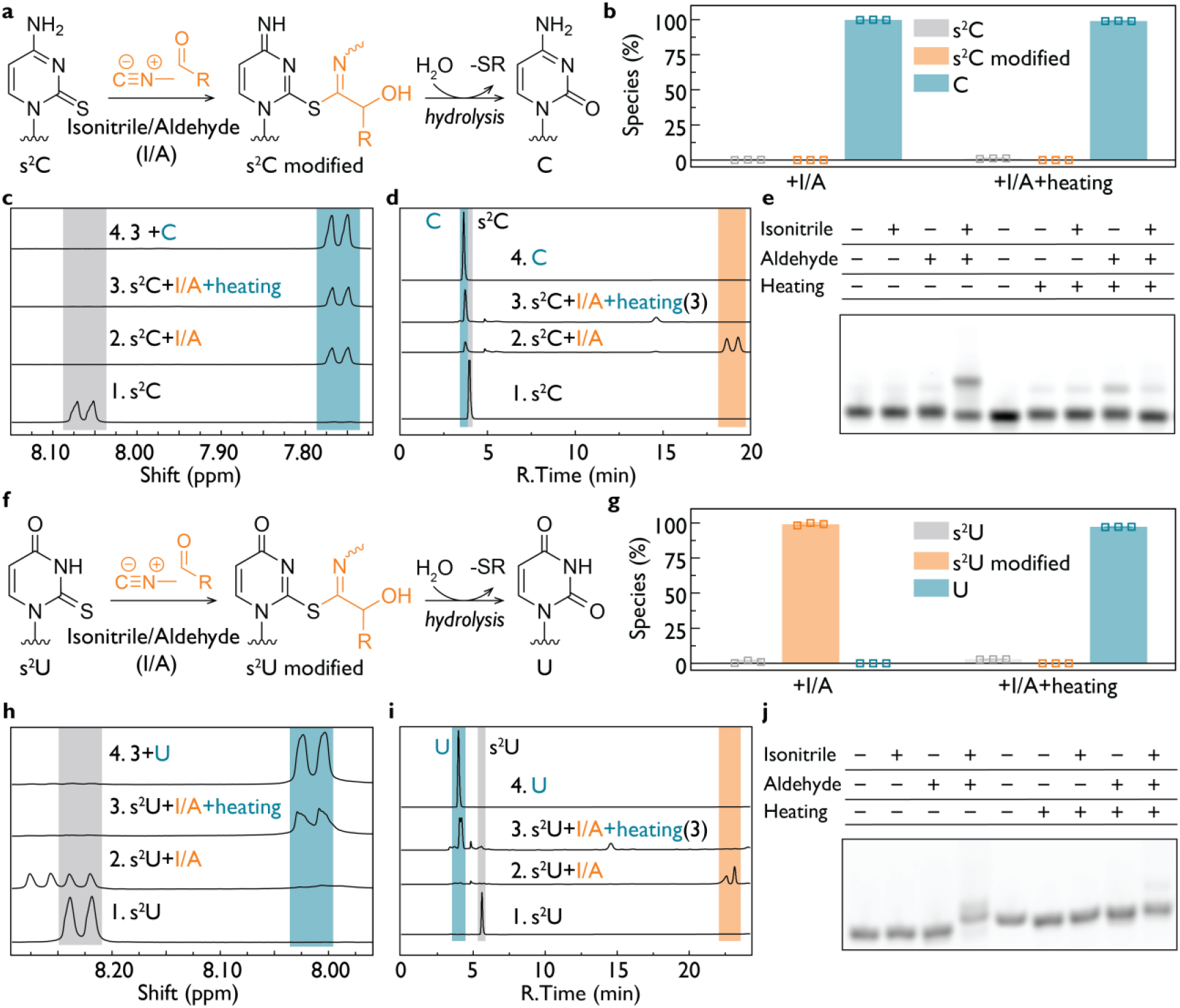
Conversion of 2-thiopyrimidines to canonical pyrimidines in the presence of isonitrile– aldehyde activation chemistry. **(a)** Reaction scheme showing s^2^C treated with isonitrile–aldehyde (I/A) activation chemistry. Nucleophilic attack on the nitrilium intermediate yields an S-imidoyl adduct that hydrolyzes to canonical C. **(b)** Reaction products of s^2^C in the presence of activation chemistry with and without heating, followed by ^1^H NMR, showing that s^2^C converts almost completely to C even without heating. **(c)** Analysis of (1) s^2^C, (2) s^2^C + activation chemistry, (3) s^2^C + activation chemistry + heat. (4) Spiking of sample 3 with authentic C confirms conversion of s^2^C to C. **(d)** Analytical HPLC confirms partial conversion of s^2^C to C upon treatment with I/A, with modified s^2^C species detected. Following heating, the dominant peak corresponds to canonical C. **(e)** EMSA analysis of s^2^C-containing RNA (C_4_s^2^CC_5_) showing that modification requires both isonitrile and aldehyde and that the modified adduct is hydrolyzed to C as verified by qTOF MS (Figure S46). **(f)** Reaction scheme for s^2^U treated with activation chemistry, forming an S-imidoyl adduct that hydrolyzes to U. **(g)** Incubation of s^2^U with isonitrile and aldehyde forms modified adduct that converts to U only upon heating, as followed by ^1^H NMR. **(h)** ^1^H NMR analysis of (1) s^2^U, (2) s^2^U + activation chemistry, (3) s^2^U + activation chemistry + heat. (4) Spiking of sample 3 with authentic U confirms that heating leads to conversion of s^2^U to U. **(i)** Analytical HPLC confirming that s^2^U fully converts to U only after heating. **(j)** EMSA analysis of s^2^U-containing RNA (C_4_s^2^UC_5_) demonstrating that modification requires both isonitrile and aldehyde and that heating removes the modification. Reaction conditions for all panels: 25 mM nucleotides or nucleosides or 1 µM RNA oligomer (3′-Cy3-labeled), 200 mM HEPES 8.0 (or at the indicated pH), 200 mM MeNC and 4-pentenal, 24 h (or 12 h for analytical HPLC) at 18 °C. Heating, where indicated, was performed at 95 °C for 15 min.

Treatment of s^2^C with MeNC and an aldehyde resulted in rapid formation of an S-imidoyl adduct *via* nucleophilic attack of sulfur on the nitrilium intermediate (**Figure 3a**). ^1^H NMR analysis showed that s^2^C converted almost quantitatively with isonitrile–aldehyde addition to canonical cytidine even in the absence of heating (**Figure 3b–c**; **Figure S45**). Analytical HPLC confirmed formation of a product coeluting with authentic C after exposure to activation chemistry (**Figure 3d**). EMSA analysis of an s^2^C-containing RNA oligomer demonstrated that modification required both isonitrile and aldehyde components and that the modified adduct hydrolyzed to C, as confirmed by qTOF MS (**Figure 3e**; **Figure S46**). As further evidence that the reaction proceeds through S-modification of s^2^C, mass spectrometry revealed a distinct desulfurization intermediate, 2-hydroxy-N-methyl-5-hexenthioamide (**Figure S47**), formed in reactions with 4-pentenal and MeNC, alongside the expected 2-hydroxy-N-methyl-5-hexenamide byproduct^16^. In experiments performed at pH 6.0 and 7.0, s^2^C displayed the same desulfurization behavior, converting to C without requiring heating (**Figures S48–S49**). Together, these observations show that s^2^C undergoes efficient, spontaneous desulfurization under isonitrile**–**aldehyde phosphate activation conditions, yielding canonical cytidine.

s^2^U followed a similar chemical pathway but with distinct kinetics. Activation chemistry yielded an initial S-imidoyl adduct (**Figure 3f**), which accumulated as the major product at 18 °C. In contrast to s^2^C, complete conversion to native uridine required heating at 95 °C for 15 min, as demonstrated by ^1^H NMR and analytical HPLC (**Figure 3g-i**; **Figures S50**). EMSA analysis of an s^2^U-containing RNA oligomer confirmed that modification occurred only when both isonitrile and aldehyde reagents were present (qTOF MS data in **Figure S46**) and that heating removed the adduct and restored U (**Figure 3j**). In the case of s^2^U, we observed the same desulfurization thioamide intermediate (**Figure S47**), indicating that S-imidoyl modification occurs for s^2^U as well as for s^2^C. The same qualitative behavior occurred at pH 6.0 and 7.0, s^2^U consistently formed the S-imidoyl adduct under activation conditions and required heating for complete conversion to U (**Figures S51–S52**). These data indicate that s^2^U undergoes the same desulfurization pathway as s^2^C but requires heating for complete conversion to U.

In summary, isonitrile**–**aldehyde activation promotes selective and condition-dependent transformation of noncanonical 2-thiopyrimidines into their canonical counterparts *via* an S-imidoyl intermediate (**Figure S53**). The spontaneous conversion of s^2^C to C, and the thermally assisted conversion of s^2^U to U, reveal a robust pathway by which prebiotically plausible 2-thiopyrimidines could have been transformed during cycles of activation and heating. Such reactivity provides a plausible mechanism by which primordial genetic polymers incorporating thiopyrimidines could have transitioned toward the canonical pyrimidines used in modern biology.

### Protocell Integrity in the Presence of Phosphate Activation Chemistry

Fatty acid vesicles have been widely used as model protocells because they form membranes under mild conditions and support encapsulation of functional RNA molecules^4,27^. To assess the compatibility of phosphate activation chemistry with protocell stability, we prepared giant unilamellar vesicles (GUVs) composed of oleic acid^46^, with encapsulated Cy5-labeled 12-nt RNA (blue) and visualized with Rhodamine B-stained membranes (red) (**Figure 4a**). In the absence of activating reagents, vesicles maintained normal morphology and retained their RNA contents (**Figure 4b**; **Figure S54**).

**Figure 4.**
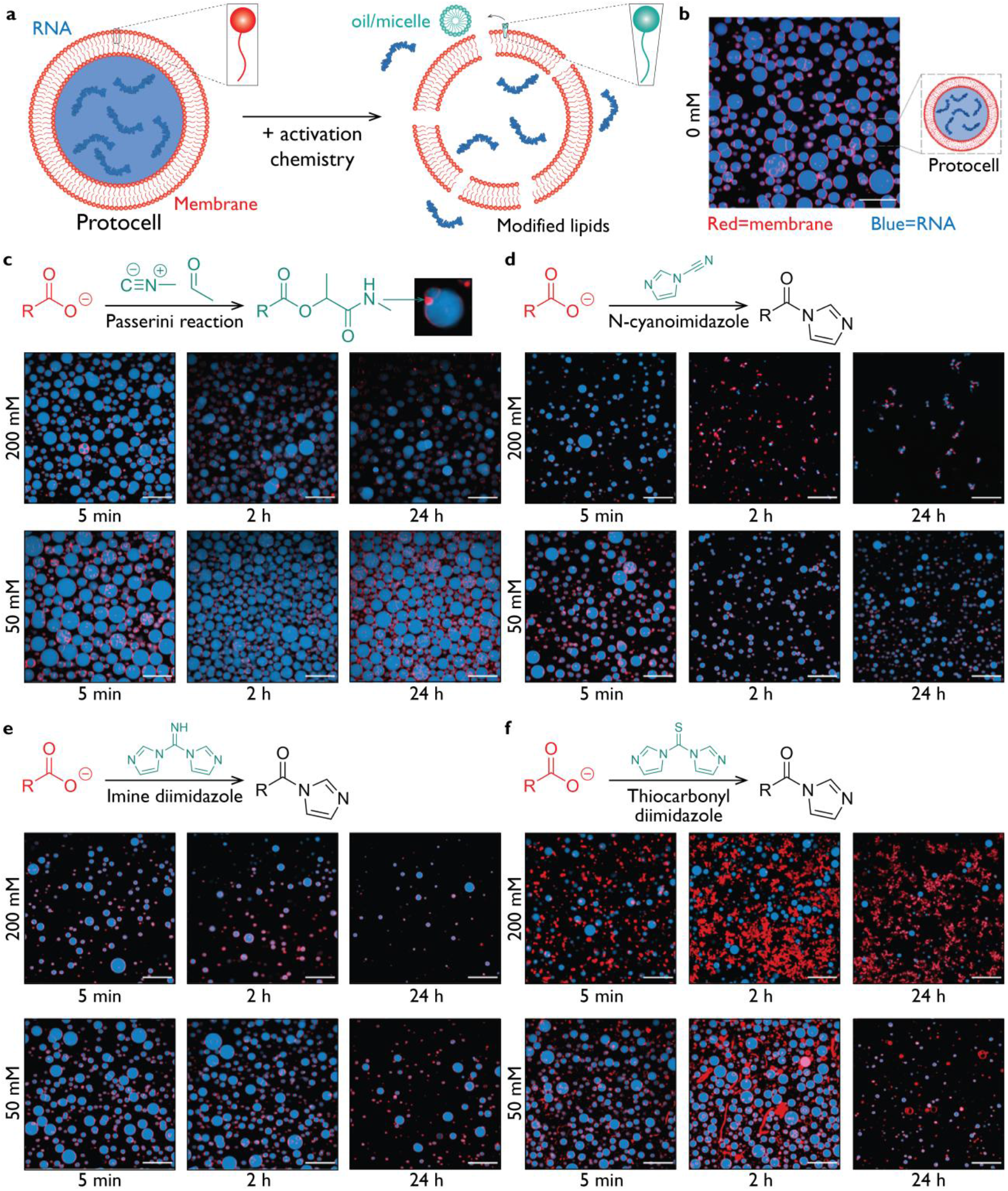
Fatty acid vesicles tolerate moderate levels of phosphate activation chemistry. **(a)** Oleic acid/oleate GUVs encapsulating Cy5-labeled 12 nt RNA (blue), membranes stained by Rhodamine B (red). Modification of oleic acid vesicles by activation chemistry reagents yields more hydrophobic lipids that can destabilize membranes. **(b)** In the absence of activation chemistry, fatty acid protocells maintain normal morphology and structural integrity. **(c)** MeNC, acetaldehyde and oleic acid generate insoluble α-acyloxy amide lipids that form oil droplets (Figure S53) embedded within vesicles and at membrane–membrane interfaces. **(d)** NCI is tolerated at 50 mM but disrupts vesicles at 200 mM. **(e)** IDI similarly preserves membrane structure at moderate concentrations but compromises vesicles at higher levels. **(f)** TCDI perturbs vesicle morphology even at 50 mM, producing elongated and distorted structures indicative of enhanced lipid reactivity. Reaction conditions: 5 mM oleic acid, 100 mM HEPES pH 8.0, 200 mM of activation chemistry reagent(s).

The Passerini reaction of acetaldehyde, MeNC, and the fatty acid carboxylate produces α-acyloxy amides (**Figure 4c**; **Figure S55**)^47^. To confirm the identity of reaction products formed *in situ*, we performed chloroform extractions of vesicle–activation reagent mixtures followed by ^1^H NMR in CDCl_3_ (Lipid Synthesis in the Supplementary Materials; **Figures S56–S60**). These analyses showed that the dominant lipid-derived product generated in the presence of MeNC and acetaldehyde was the expected Passerini α-acyloxy amide, which matched the synthetic standard (**Figure S61**). At 50 mM MeNC and acetaldehyde, oleic acid vesicles remained intact over several days. At 200 mM, however, vesicles exhibited oil-like droplet formation at the membrane interfaces (**Figure 4c**, inset; **Figures S62–S68**). These droplets are consistent with accumulation of the hydrophobic Passerini product, which perturbs membrane packing and induces local phase separation.

We next examined three acylimidazole activating agents, NCI, IDI, and TCDI. All three reagents were compatible with vesicles at 50 mM, with membranes preserving morphology and retaining their RNA cargo (**Figure 4d–f**, upper panels; **Figures S69–S86**). Chloroform extractions followed by CDCl_3_ NMR revealed that reactions between oleic acid and acylimidazole reagents predominantly yielded N-oleoylimidazolide (Lipid Synthesis in the Supplementary Materials; **Figures S87–S93**), with no detectable oleic anhydride under the conditions tested (**Figures S91– S93**). This is consistent with selective activation of the fatty acid carboxylate by IDI and NCI. Because oleic anhydride is known to cause severe membrane destabilization and droplet formation, its absence explains why moderate concentrations of these reagents do not disrupt vesicle structure. At 200 mM, both NCI and IDI caused substantial vesicle loss and membrane defects (**Figure 4d– e**; **Figures S69–S80**). Such effects could arise from pH fluctuations or from formation of acylimidazole intermediates, or formation of fatty acid anhydrides, any of which might compromise membrane packing and stability^46,48,49^. TCDI altered vesicle morphology even at 50 mM, producing elongated and distorted structures (**Figure 4f**; **Figures S81–S86**), indicative of enhanced reactivity with membrane lipids.

Together, these results establish that the effects of phosphate activation chemistry on protocell membranes are strongly concentration dependent. Moderate levels of both activation reagents are compatible with fatty acid vesicle integrity, whereas high concentrations lead to accumulation of lipid-derived products that disrupt bilayer packing, modify pH, and reduce vesicle stability. The results highlight an important constraint on prebiotic phosphate activation chemistry, which must preserve the compartmentalization required for protocellular function.

## DISCUSSION

Phosphate activation chemistry is central to all models of nonenzymatic RNA replication because it supplies the chemical energy required for forming the reactive phosphorimidazolides that drive primer extension and ligation^10,13,16,34^. However, activation agents do not act solely on phosphate; they also react with ribose hydroxyls and nucleobases^34,50^. Our results show that these competing pathways might have imposed chemical selection pressures that would have shaped both the primordial genetic alphabet (**Table 1**) and the physical environment in which replication occurred.

**Table 1.**
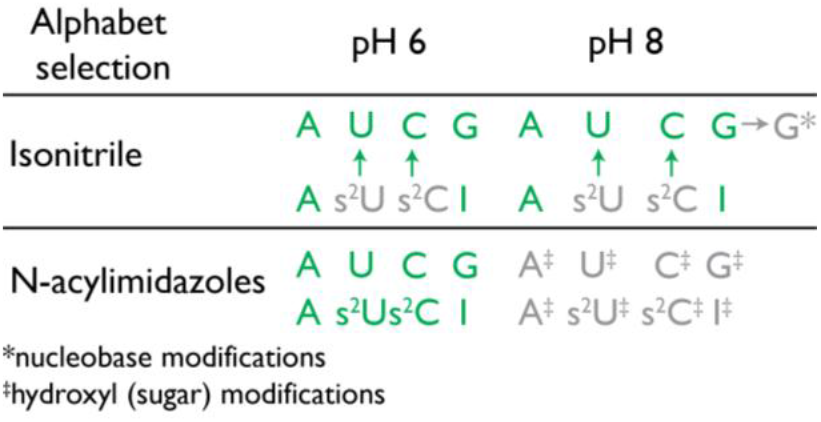
Compatibility of prebiotic phosphate activation chemistries with candidate primordial genetic alphabets. Summary of how isonitrile-based and acylimidazole-based activation agents interact with canonical and noncanonical nucleotides at pH 6 and pH 8. Isonitrile–aldehyde activation chemistry promotes conversion of 2-thiopyrimidines to canonical pyrimidines at both pH values. Acylimidazoles are broadly compatible at pH 6 but at pH 8 induces extensive 2′/3′-OH acylation.

A key outcome of our work is the demonstration that G is uniquely vulnerable to irreversible N1 imidoyl modification under isonitrile-type activation chemistry. This constraint suggests several possibilities for RNA copying chemistry. One is that G may have been sparsely used in primordial RNA, consistent with proposals that early alphabets omitted strongly pairing nucleobases^21,51,52^. Alternatively, if activation reagent concentrations were low, N1-modified G is minimal. Because N1-modified purine nucleotides cannot base pair^53,54^, they cannot participate in primer extension, and would not act as inhibitors, this chemistry could simply remove a fraction of G from the reactive pool without catastrophic consequences. A third possibility is that early Earth environments contained bulkier or milder isonitriles^39^, structurally analogous to modern sterically hindered aldehydes^55^, that would activate phosphate efficiently while reducing nucleobase reactivity. Each scenario highlights that the chemical environment would have constrained which purines could persist long enough to participate in replication.

Chemical modifications may also have influenced the effective nucleotide pool available for replication. Because substrates with lower pKa values^38^ are more readily modified, oxidatively damaged nucleotides, such as 5-hydroxyuridine, 5-hydroxymethyluridine, and 8-oxoguanosine, may have been preferentially scavenged or removed. This raises the possibility that activation chemistry could have been used as a chemical quality-control mechanism, enriching protocells in undamaged nucleotides.

The pH dependence of activation is highly consequential. At pH 6, nucleobase and hydroxyl modifications disappear almost entirely for both activation systems, pinpointing a regime in which selective phosphate activation can occur with minimal side chemistry. However, this observation raises two problems. First, primer extension is slow at pH 6^10^, implying that replication would require either environmental pH fluctuations or alternative catalysts capable of accelerating copying under mildly acidic conditions. Iron(II), for example, has been shown to enhance primer extension rates at lower pH^50^, and such metal-dependent mechanisms may help reconcile chemical activation with efficient copying. Second, fatty acid vesicles are unstable at pH 6^8,46,56^ unless stabilized by low pKa amphiphiles such as alkyl phosphates^24,57^. Thus low-pH-compatible membrane systems may have been essential for protocells exposed to activation chemistry.

Alternatively, further research may reveal new activation chemistries that function at higher pH without causing deleterious modifications.

An intriguing aspect of our findings is that 2-thiopyrimidines undergo slow, condition-dependent conversion to canonical C and U in the presence of isonitrile–aldehyde activation. This transformation provides a chemically plausible pathway by which a primordial alphabet containing s^2^C, s^2^U, and I could gradually transition to the canonical C, U, and G over time^23,34^. Moreover, the desulfurization chemistry proceeds even under mildly acidic conditions where most side reactions are suppressed. Such slow “chemical aging” could offer functional advantages. For example, 2-thiopyrimidines support more uniform nonenzymatic copying through isoenergetic base pairing, while canonical pyrimidines provide structural stability and functionality for folded RNAs^58^. However, the feasibility of this scenario will depend on how desulfurization rates compare to protocell replication cycles.

Although fatty acid vesicles tolerated moderate concentrations of both classes of activation reagents, higher concentrations led to membrane disruption. Our observations suggest that protocell survival might have required a narrow concentration range of activation reagents that was high enough to drive nucleotide activation yet low enough to preserve compartment integrity. For isonitrile-based activation agents, the Passerini-type byproduct generated hydrophobic oil droplets associated with vesicle surfaces. By partitioning fatty acids into oil-like domains, this process altered vesicle morphology and could reduce internal volume, thereby increasing the local concentration of encapsulated solutes. For acylimidazoles, the formation of N-oleoylimidazolide and oleic anhydride altered membrane stability but might enable the synthesis of fatty acyl esters^48,57^. Such a pathway could allow protocells to grow by absorbing fatty acids, yet maintain a more stable membranes by constantly converting a fraction of the fatty acid to fatty acyl esters.

Overall, our results show that activation chemistry, nucleotide structure, pH, and membrane stability are tightly linked variables that together constrain the emergence of a functional genetic system. Each candidate alphabet implies a compatible activation chemistry and environmental regime, and conversely each activation chemistry highlights which nucleotides could realistically persist. This interdependence suggests that the selection of the primordial genetic alphabet might not have been governed solely by base-pairing thermodynamics, but also by the chemical compatibility of its component nucleotides with the pathways that supplied activated substrates and sustained protocellular compartments.

## ASSOCIATED CONTENT

### Supporting Materials

Materials and Methods with extended experimental procedures, NMR and MS analysis, PAGE gels, Supporting Figures S1–S93, and Tables S1–S2 (PDF).

## AUTHOR INFORMATION

## COMPETING INTERESTS

F.B., J.Z., and J.W.S are inventors on a provisional U.S.A. patent application (63/818,859) submitted by the University of Chicago.

## ACKNOWLEDGEMENTS

F.B., J.Z., J.W.S., acknowledge funding from Howard Hughes Medical Institute (HHMI). J.W.S. is an Investigator of the Howard Hughes Medical Institute. F.B. acknowledges funding from the European Molecular Biology Organization through a Long-Term Fellowship (ALTF 106-2023). We are grateful to Dr. Daniel Duzdevich, Anmol Mishra, and Dr. Ben W. F. Colville for helpful discussions, and to Dr. Josh Kurutz for expert advice on NMR spectroscopy.

