## Supplementary Materials for "Influence of phosphate activation chemistry on the selection of the primordial genetic alphabet"

**This PDF file includes:**
Materials and Methods
Figs. S1 to S93
Tables S1 to S2

### MATERIALS AND METHODS

#### *Materials*

Methyl isonitrile (ATEH99BCA209) and 4-pentenal (CDS000721) were purchased from Millipore Sigma. The activation reagents IDI (A110025), TCDI (A107641), and NCI (A279193) were obtained from Ambeed. Oleic acid (O1383), 1 M potassium hydroxide (24-4720), NMR tubes (Z565229), and deuterium oxide (1133660100) were purchased from Millipore Sigma. Buffer solutions, including 1 M HEPES pH 8.0 (J63578.AP) and 1 M MES pH 6.0 (J60763.AP), were obtained from ThermoFisher or prepared fresh for other pH values. Nucleotides (UMP, AMP, CMP, GMP, IMP) and nucleosides (2-thiouridine, 2-thiocytidine, xanthosine, inosine, and dideoxyuridine) were sourced from Ambeed or Millipore Sigma.

#### *Oligonucleotide Solid-State Synthesis*

RNA and DNA oligonucleotides were synthesized on a solid-phase synthesizer using standard phosphoramidites and purified by preparative denaturing PAGE or reverse-phase HPLC as appropriate[1]. DNA oligonucleotides with 3′ terminal dideoxynucleotide were prepared using the corresponding reverse phosphoramidites. After deprotection and cleavage, all oligonucleotides were desalted (Sep-Pak C18), lyophilized, and resuspended in nuclease-free water. Fluorescently labeled 3′ dideoxy DNA strands were generated by conjugation of ATTO550 to a 5′ terminal amine using NHS chemistry. Concentrations were determined by UV absorbance, and samples were stored as 10 µM stocks at −20 °C for routine use.

#### *Nucleotide and Oligonucleotide Reactions with Isonitrile–Aldehyde Activation*

Nucleotides and nucleosides were reacted under isonitrile–aldehyde activation conditions by dissolving each substrate (25 mM) in 200 mM HEPES pH 8.0 containing 100 mM methyl isonitrile and 100 mM 4-pentenal. Each reaction mixture contained 10% (v/v) $D_2O$ for NMR locking, was prepared in a final volume of 550–600 µL, transferred to 5 mm NMR tubes, and incubated for 24 h at 20–22 °C prior to NMR analysis.

RNA oligonucleotides ($U_{10}$, $A_{10}$, $C_{10}$, $I_{10}$, and $(GA)_5$) bearing a free 5′-OH and a 3′-Cy3 label were synthesized as described above and incubated at 1–3 µM for 24 h in 200 mM HEPES pH 8.0 containing 200 mM methyl isonitrile and 200 mM 4-pentenal at room temperature. Reactions were quenched by 1:10 dilution into 98% (v/v) formamide containing 5 mM EDTA, and 1 µL aliquots were resolved on 20% (v/v) denaturing polyacrylamide gels in 1× TBE. Gels were pre-run at 10 W for 10 min and then at 30 W for 1.5–2.5 h before imaging on a Typhoon™ scanner in the Cy3 channel.

#### *Nucleotide and Oligonucleotide Reactions with Acylimidazole Activation*

Nucleotides and nucleosides (25 mM) were incubated with 200 mM HEPES pH 8.0 or 200 mM MES pH 6.0 containing 200 mM imidazole-based activating reagent (imine diimidazole (IDI), N-cyanoimidazole (NCI), or thiocarbonyl diimidazole (TCDI)). Reaction mixtures contained 10% (v/v) $D_2O$ and were incubated for 24 h at 20–22 °C. For experiments requiring the maintenance of acidic conditions at pH 6.0, 200 mM HCl was added at the start of the reaction to counteract the pH increase caused by imidazole release during IDI hydrolysis.

RNA oligonucleotides ($U_{10}$, $A_{10}$, $C_{10}$, and $(GA)_5$) containing a free 5′-OH and a 3′-Cy3 label were prepared as described above. DNA oligonucleotides ($dT_{10}$, $dA_{10}$, $dC_{10}$, and $d(GA)_5$) were synthesized using reverse phosphoramidites, equipped with a 5′-ATTO550 label introduced by NHS-ester coupling to a C6-amine, and terminated with a 3′ dideoxycytidine. For gel-based assays, each oligonucleotide (1 µM) was incubated in 200 mM HEPES pH 8.0 or 200 mM MES pH 6.0 containing 200 mM IDI, NCI, or TCDI for 24 h at room temperature. Reactions were quenched by 1:10 dilution into 98% formamide with 5 mM EDTA, and 1 µL aliquots were resolved on 20% (v/v) denaturing polyacrylamide gels in 1× TBE. Gels were pre-run at 10 W for 10 min and electrophoresed at 25 W for 2–2.5 h before Typhoon™ imaging in the Cy3 (RNA) or ATTO550 (DNA) channels.

***NMR Data Acquisition***

$^{1}H$ and $^{31}P$ NMR spectra were collected on either a 400 MHz Bruker Avance III or a 600 MHz Bruker Avance III HD spectrometer equipped with broadband-observe or TCI cryoprobes. All spectra were acquired at 298 K. For $^{1}H$ NMR, 128–256 scans were recorded with automatic receiver gain and excitation-sculpting water suppression. Processed spectra were zero-filled to 64k points and apodized with 0.3–0.5 Hz exponential line broadening. Chemical shifts were referenced to residual HEPES resonances.

For $^{31}P$ NMR, proton-decoupled spectra were collected using standard Bruker pulse programs, typically with 256 scans. Chemical shifts were referenced to external 85% $H_3PO_4$. All spectra were processed using MestReNova (v14.2.0).

Quantification of nucleotide modification was performed by integrating diagnostic resonances corresponding to unmodified substrate and product species. Percent conversion was calculated using:

$$\%\ \text{modified} = \frac{I_{\text{adduct}}}{I_{\text{adduct}} + I_{\text{unmodified}}} \times 100$$

Independent replicates agreed within ±3%.

***PAGE Band Quantification***

Fluorescent gel bands were quantified using ImageQuant™ TL (Typhoon software) with background subtraction. Percent modification was calculated as:

$$\%\ \text{modified} = \frac{I_{\text{shifted}}}{I_{\text{shifted}} + I_{\text{unshifted}}} \times 100$$

Values were consistent across at least three independent experiments.

***Mass Spectrometry***

For analysis of nucleobase-modified species, guanosine 5′-monophosphate (GMP, 20 mM) was incubated with 200 mM methyl isonitrile and 200 mM acetaldehyde, and the pH was adjusted to 8.5 prior to reaction. Adenosine 5′-monophosphate (AMP, 20 mM) was reacted with 200 mM IDI at pH 8.0. Immediately before measurement, each reaction mixture was diluted to 100 µM in methanol. Mass spectra were acquired by direct infusion using a LTQ-XL linear ion trap mass spectrometer (Thermo Fisher Scientific) operated in negative-ion electrospray mode. Standard ESI source parameters supplied by the manufacturer were used, and mass assignments were made based on the expected m/z values and isotopic distributions.

High-resolution mass spectrometry of oligonucleotides was performed using an Agilent 1200 HPLC system coupled to an Agilent 6520 quadrupole time-of-flight LC/MS operated in negative electrospray ionization mode. Oligonucleotides desalted using C18 Zip Tip were separated by ion-pair reversed-phase HPLC on a 100 mm × 1 mm XBridge C18 column (3.5 µm, Waters) maintained at 50 °C. Chromatographic separation was achieved using a linear gradient of 2.5-15% methanol in 200 mM 1,1,1,3,3,3-hexafluoro-2-propanol (HFIP) containing 1 mM triethylamine (TEA), pH 7.0.

The Q-TOF instrument was operated in negative-ion mode over an m/z range of 239–3200 with a scan rate of 1 spectrum/s. Mass spectra were calibrated using manufacturer-supplied reference ions, and data were analyzed with Agilent MassHunter Qualitative Analysis software. Intact oligonucleotides and modified species were identified based on accurate mass and expected isotopic distributions.

#### *Analytical Reverse-Phase HPLC of Nucleotides and Nucleosides*

Analytical HPLC analyses of nucleotide and nucleoside reaction mixtures were performed on a Shimadzu LC system equipped with a photodiode array detector. Separations were carried out on an Atlantis T3 C18 column (4.6 × 150 mm, 3 µm, Waters) using a mobile phase consisting of 50 mM triethylammonium acetate (TEAA) pH 7.0 (solvent A) and acetonitrile (solvent B). The column was maintained at ambient temperature, and elution was monitored at 260 nm.

For uridine 5′-monophosphate (UMP), samples were analyzed using a linear gradient from 3% to 10% acetonitrile over 12 min, followed by an additional 5 min gradient from 10% to 35% acetonitrile. For 2-thiocytidine ($s^2C$) and 2-thiouridine ($s^2U$), separations were performed using a gradient from 3% to 10% acetonitrile over 12 min, followed by 10 min from 10% to 35% acetonitrile. Chromatograms were processed using Shimadzu LabSolutions software, and product identities were assigned by comparison with authentic standards of U, C, $s^2C$, and $s^2U$.

#### *Fatty Acid Protocell Preparation*

Oleic acid vesicles were prepared by inducing the micelle to vesicle transition through a controlled pH drop, following previously described procedures[2]. To generate oleate micelles, 425 µL of Milli-Q water was mixed with 65 µL of 1 M potassium hydroxide, after which 15.6 µL of neat oleic acid was added. The mixture was vortexed vigorously until a clear solution was obtained, indicating complete micelle formation.

Vesicles were assembled by preparing a solution containing 100 mM HEPES pH 8.0, 100 mM sucrose, and 5 µM Cy5-labeled 12 nt RNA. This solution was made by combining 20 µL of 1 M HEPES pH 8.0, 20 µL of 1 M sucrose, 10 µL of 100 µM Cy5-RNA, 130 µL of Milli-Q water, and 20 µL of the oleate micelle solution in a 1.5 mL microcentrifuge tube. The mixture was incubated on a horizontal shaker at 30 rpm for 24 to 48 h to promote vesicle formation.

Oleic acid vesicles were purified by sucrose–glucose gradient washing. A 50 µL aliquot of the vesicle suspension was diluted into 450 µL of washing buffer consisting of 100 mM HEPES pH 8.0 and 100 mM glucose, followed by centrifugation at 2000 × g for 3 min. The supernatant (approximately 450 µL) was discarded. This wash was repeated twice to remove unencapsulated Cy5-RNA.

***Confocal Microscopy Imaging of Oleic acid vesicles Treated with Phosphate Activation Agents***

Purified vesicles (1 µL) were mixed with 0.2 µL of 5 µM Rhodamine B to stain the membrane and 3.8 µL of washing buffer containing 100 mM HEPES pH 8.0 and 100 mM glucose. Confocal images were collected on a Nikon A1R HD25 confocal laser scanning microscope equipped with a LU-N4/N4S four-laser unit. The focal plane was first identified using Rhodamine B fluorescence excited with a 561 nm laser.

Imaging was performed using minimal laser power (<1%) for both Rhodamine B and Cy5 channels, with Cy5 fluorescence excited at 640 nm. After baseline imaging, 1 µL of the phosphate activation reagent, prediluted in anhydrous DMSO to avoid premature hydrolysis, was added directly to the sample to yield a final concentration of 50 mM or 200 mM of methyl isonitrile–acetaldehyde, IDI, NCI, or TCDI. Control samples received an equivalent volume of DMSO.

***Fatty Acid Reaction Product Extraction and Their NMR Analysis***

Oleic acid vesicles (5 mM) were incubated with each activation reagent (methyl isonitrile and acetaldehyde, NCI, IDI, or TCDI) at 50 mM or 200 mM for 24 h at 18 °C. After incubation, samples were lyophilized to remove the aqueous phase. The resulting dry material was extracted with $CDCl_3$ to isolate lipophilic species for NMR analysis. Extracts were examined by $^{1}H$ and $^{13}C$ NMR and the lipid standards were authenticated by $^{1}H$–$^{13}C$ and $^{15}N$–$^{1}H$ HSQC and HMBC experiments in deuterated $CDCl_3$.

Oleic acid, oleic anhydride (Millipore Sigma), N-oleoylimidazolide, and the Passerini-derived α-acyloxy amide were used as reference standards. These standards were analyzed in parallel with the vesicle extracts to identify oleic acid, N-oleoylimidazolide, oleic anhydride, and the Passerini-derived α-acyloxy amide formed under the different activation conditions.

***Synthesis of Lipid Standards***

N-oleoylimidazolide (IUPAC (Z)-1-(1H-imidazol-1-yl)octadec-9-en-1-one) was synthesized by dissolving imidazole (11.72 mmol) in 20 mL anhydrous THF under nitrogen with stirring, followed by dropwise addition of oleoyl chloride (5.86 mmol). The reaction was stirred overnight at 18 °C. The product was collected by filtration, washed three times with THF, and dried by rotary evaporation. The resulting N-oleoylimidazolide was confirmed to be 95.32% pure by $^{1}H$ NMR, $^{13}C$ NMR, and HSQC and HMBC spectra in deuterated $CDCl_3$.

The Passerini-derived α-acyloxy amide standard (IUPAC 1-(methylamino)-1-oxopropan-2-yl oleate) was prepared by mixing oleic acid (50 mM), methyl isonitrile (400 mM), and acetaldehyde (400 mM) in $CHCl_3$ for 2 days at 18 °C, followed by solvent removal by evaporation. The identity of the resulting α-acyloxy amide product was confirmed by $^{1}H$, $^{13}C$, HSQC, and HMBC NMR in $CDCl_3$ with the purity of 99.51%.

**Synthesis of α-acyloxy amide Passerini product of oleic acid**

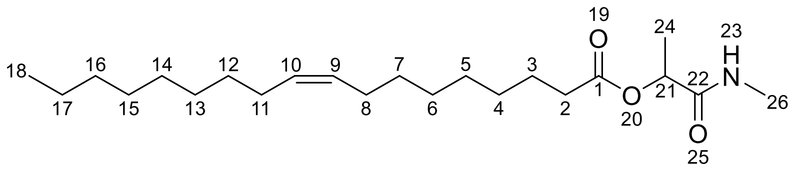


1-(methylamino)-1-oxopropan-2-yl oleate

$^{1}H$ NMR (400 MHz, $CDCl_3$) δ 6.14 (s, 1H), 5.44 – 5.28 (m, 2H), 5.23 (q, *J* = 6.8 Hz, 1H), 2.85 (d, *J* = 4.9 Hz, 3H), 2.38 (t, *J* = 7.5 Hz, 2H), 2.01 (tt, *J* = 6.3, 3.4 Hz, 4H), 1.66 (dq, *J* = 14.8, 6.3 Hz, 2H), 1.46 (d, *J* = 6.8 Hz, 3H), 1.30 (dd, *J* = 17.1, 5.1 Hz, 20H), 0.97 – 0.80 (m, 3H).

$^{13}C$ NMR (400 MHz, $CDCl_3$) δ 172.23, 171.06, 130.09, 129.67, 70.45, 34.35, 31.92, 29.78, 29.69, 29.53, 29.34, 29.16, 29.10, 29.07, 27.24, 27.16, 26.00, 24.87, 22.70, 22.69, 17.95, 14.12. **Yield: 99.51 %**

Two-dimensional NMR spectra, including $^{1}H$–$^{1}H$ COSY, $^{13}C$–$^{1}H$ HSQC, $^{13}C$–$^{1}H$ HMBC, $^{15}N$–$^{1}H$ HSQC, and $^{15}N$–$^{1}H$ HMBC, are shown in **Figures S56–S60**.

**Synthesis of N-oleoylimidazolide**

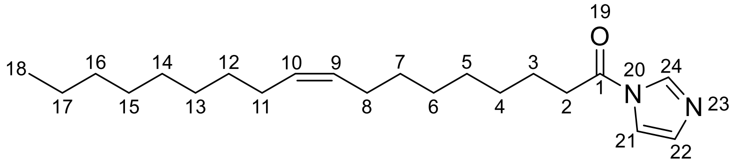


(Z)-1-(1H-imidazol-1-yl)octadec-9-en-1-one

$^{1}H$ NMR (400 MHz, $CDCl_3$) δ 8.16 (t, *J* = 1.1 Hz, 1H), 7.48 (t, *J* = 1.5 Hz, 1H), 7.10 (dd, *J* = 1.7, 0.8 Hz, 1H), 5.50 – 5.24 (m, 2H), 2.85 (t, *J* = 7.4 Hz, 2H), 2.03 (d, *J* = 6.7 Hz, 4H), 1.80 (p, *J* = 7.4 Hz, 2H), 1.49 – 1.18 (m, 20H), 1.00 – 0.80 (m, 3H).

$^{13}C$ NMR (400 MHz, $CDCl_3$) δ 169.51, 136.16, 131.02, 130.13, 129.63, 116.01, 35.27, 31.92, 29.78, 29.65, 29.54, 29.34, 29.18, 29.03, 28.98, 27.24, 27.15, 24.11, 22.70, 22.69 14.12. **Yield: 95.32 %**

Two-dimensional NMR spectra, including $^{1}H$–$^{1}H$ COSY, $^{13}C$–$^{1}H$ HSQC, $^{13}C$–$^{1}H$ HMBC, and $^{15}N$–$^{1}H$ HMBC, are shown in **Figures S87–S90**.

**Figure S1.**

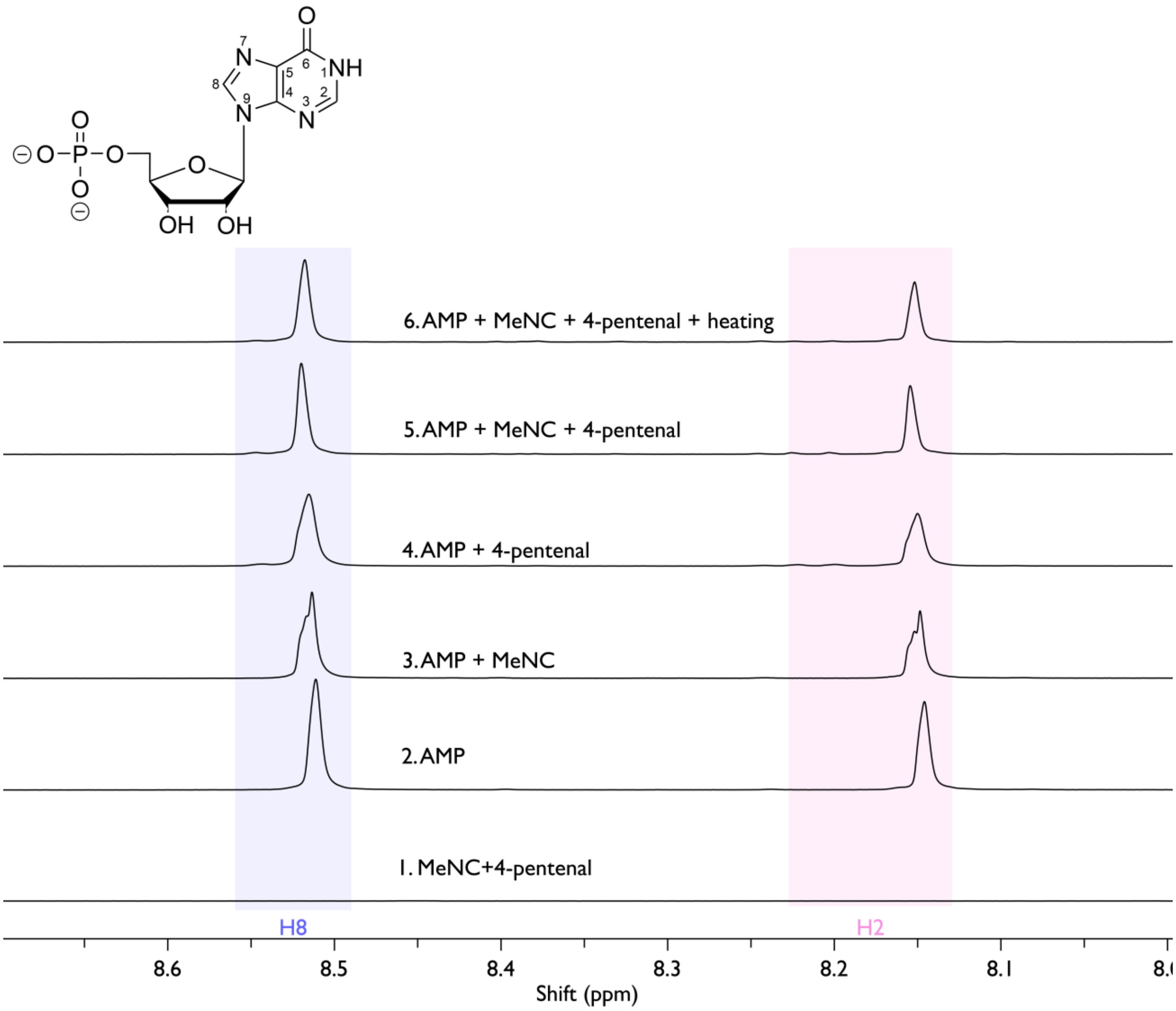


**Figure S1.** [1]H NMR analysis of adenosine 5′-monophosphate (AMP; 25 mM) incubated with methyl isocyanide (MeNC, 100 mM) and 4-pentenal (100 mM) in 200 mM HEPES, pH 8.0, in 10 % $D_2O$ and 90 % $H_2O$ for 12 hours at 18 °C. Spectra correspond to : (1) 100 mM MeNC plus 100 mM 4-pentenal; (2) AMP; (3) AMP with 100 mM MeNC; (4) AMP with 100 mM 4-pentenal; (5) AMP with 100 mM MeNC and 100 mM 4-pentenal; (6) AMP with 100 mM MeNC and 100 mM 4-pentenal after heating at 95 °C for 15 minutes. Key resonances are highlighted, and the chemical-shift changes associated with N1 modification are marked with black rectangles.

**Figure S2.**

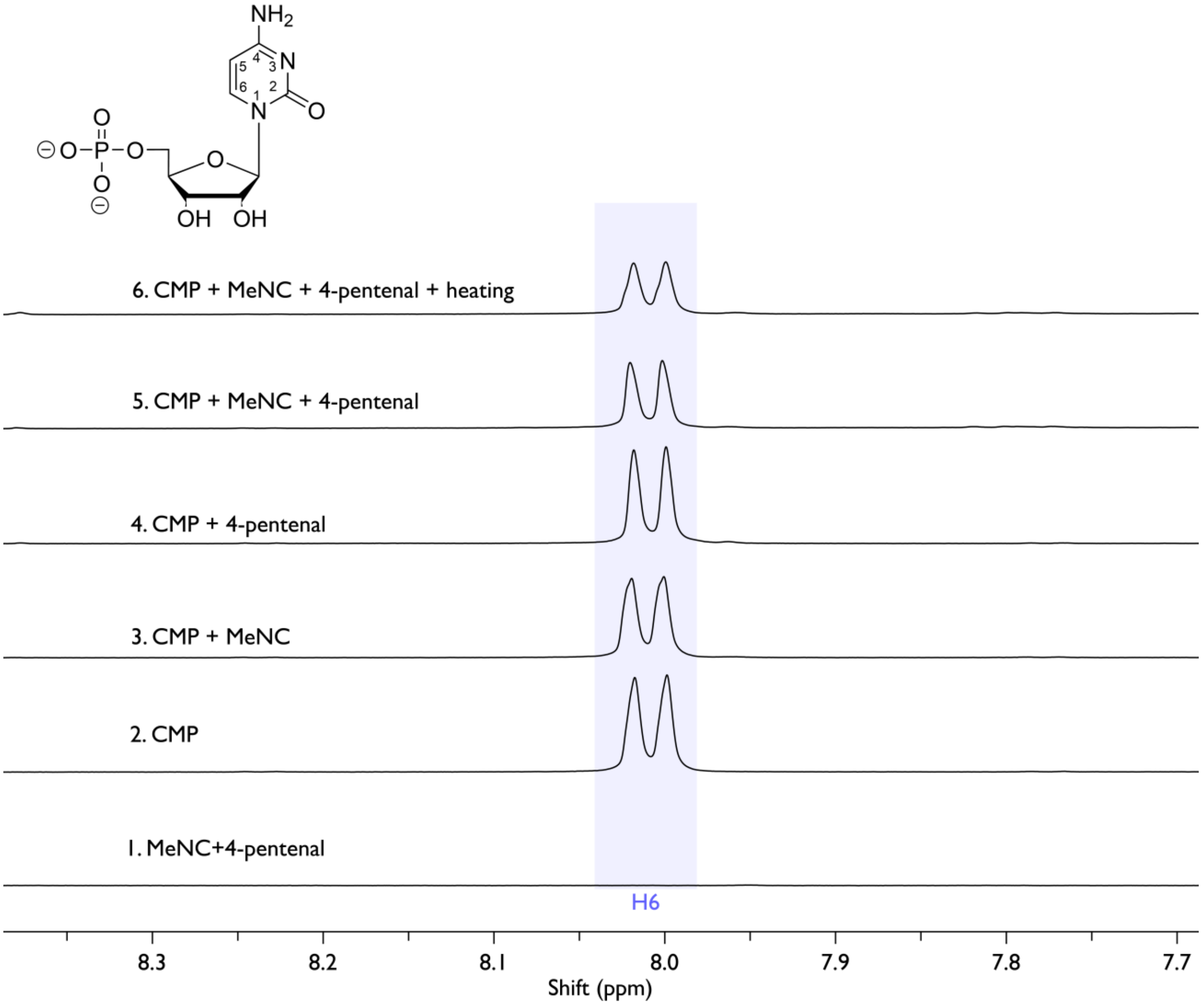


**Figure S2.** $^1$H NMR analysis of cytidine 5′-monophosphate (CMP; 25 mM) incubated with methyl isocyanide (MeNC, 100 mM) and 4-pentenal (100 mM) in 200 mM HEPES, pH 8.0, in 10 % $D_2O$ and 90 % $H_2O$ for 12 hours at 18 °C. Spectra correspond to : (1) 100 mM MeNC plus 100 mM 4-pentenal; (2) CMP; (3) CMP with 100 mM MeNC; (4) CMP with 100 mM 4-pentenal; (5) CMP with 100 mM MeNC and 100 mM 4-pentenal; (6) CMP with 100 mM MeNC and 100 mM 4-pentenal after heating at 95 °C for 15 minutes. Key resonances are highlighted, and the chemical-shift changes associated with N1 modification are marked with black rectangles.

**Figure S3.**

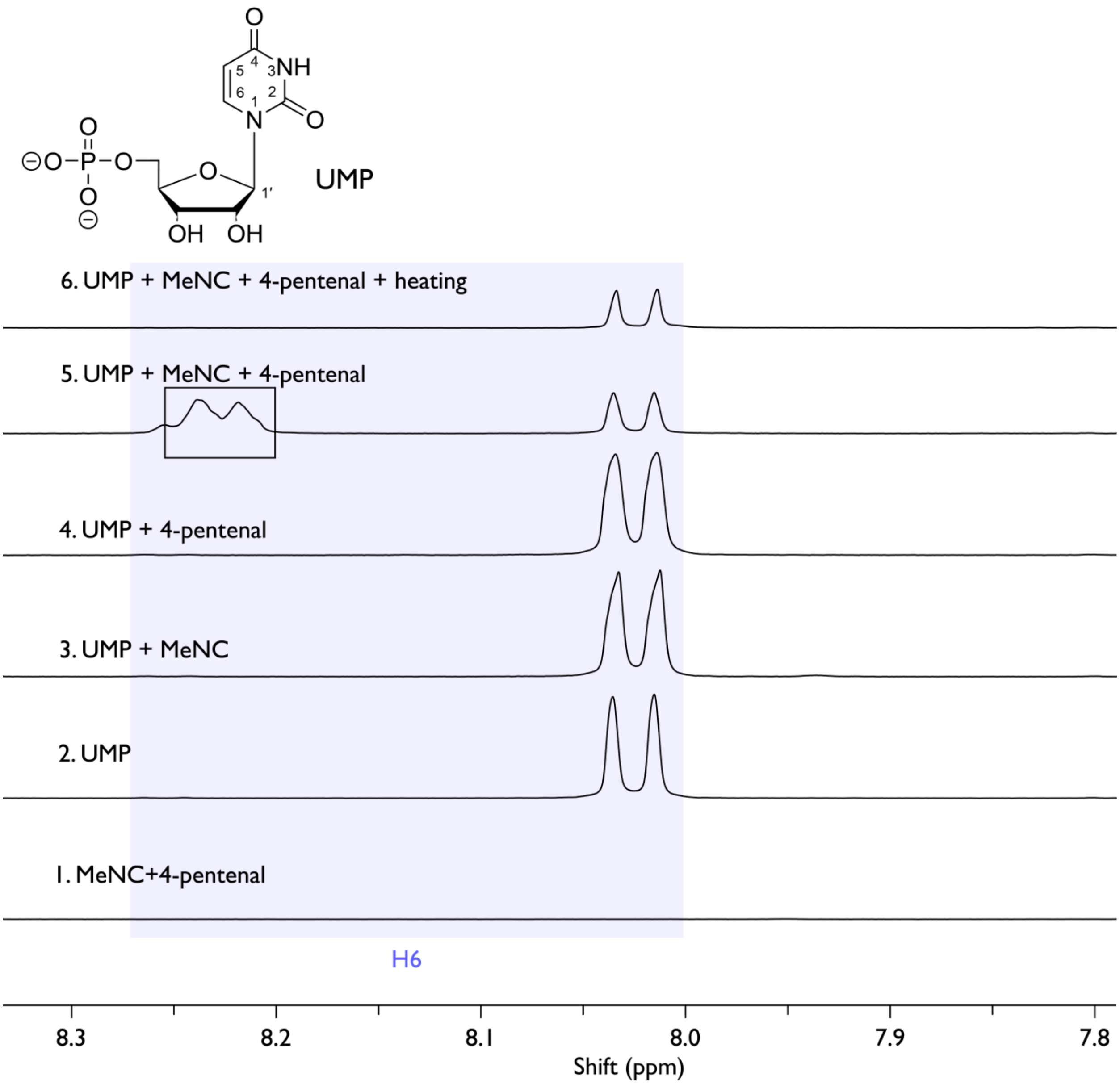


**Figure S3.** $^1$H NMR analysis of uridine 5′-monophosphate (UMP; 25 mM) incubated with methyl isocyanide (MeNC, 100 mM) and 4-pentenal (100 mM) in 200 mM HEPES, pH 8.0, in 10 % $D_2O$ and 90 % $H_2O$ for 12 hours at 18 °C. Spectra correspond to : (1) 100 mM MeNC plus 100 mM 4-pentenal; (2) UMP; (3) UMP with 100 mM MeNC; (4) UMP with 100 mM 4-pentenal; (5) UMP with 100 mM MeNC and 100 mM 4-pentenal; (6) UMP with 100 mM MeNC and 100 mM 4-pentenal after heating at 95 °C for 15 minutes. Key resonances are highlighted, and the chemical-shift changes associated with N1 modification are marked with black rectangles.

**Figure S4.**

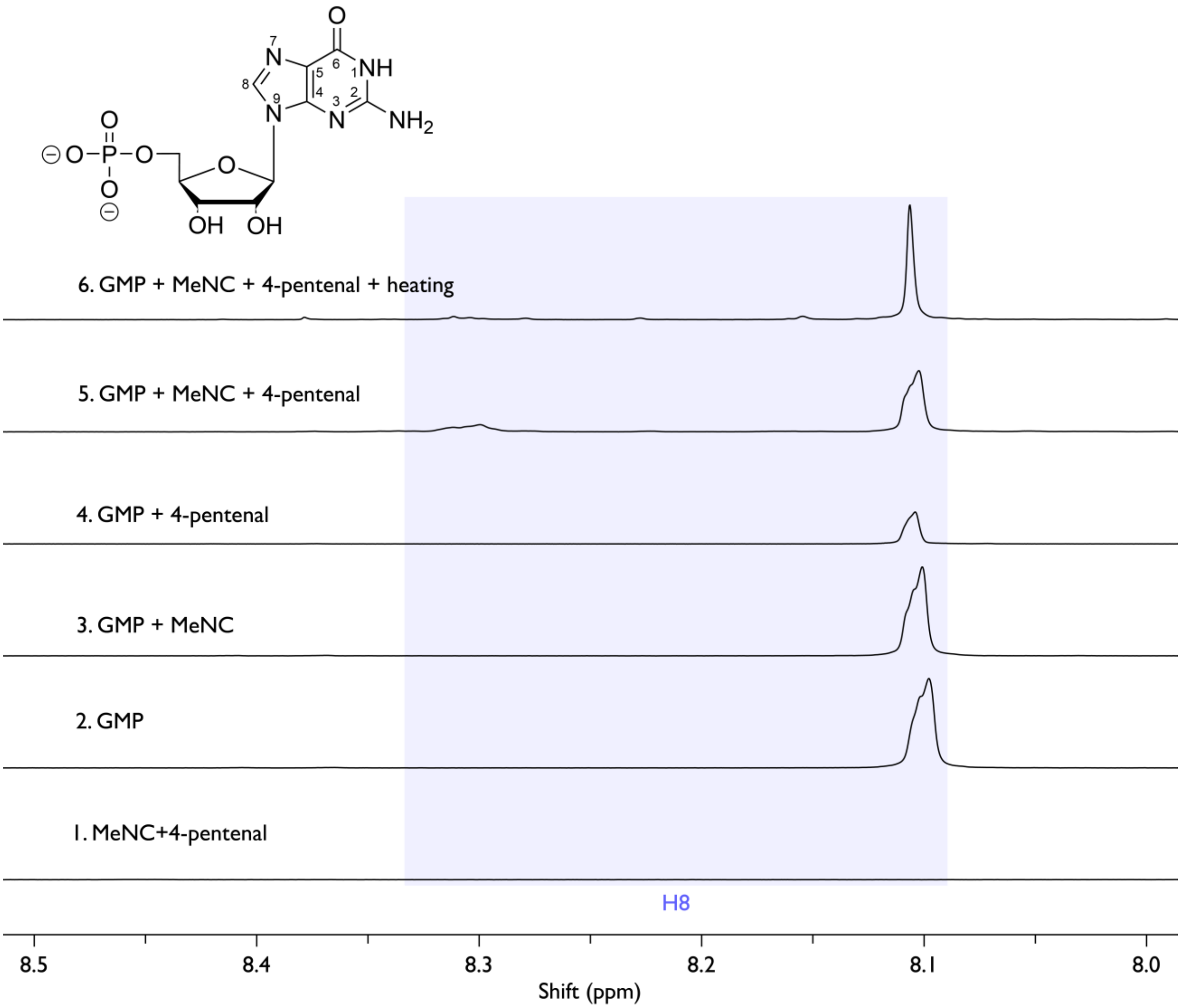


**Figure S4.** $^1$H NMR analysis of guanosine 5′-monophosphate (GMP; 25 mM) incubated with methyl isocyanide (MeNC, 100 mM) and 4-pentenal (100 mM) in 200 mM HEPES, pH 8.0, in 10 % $D_2O$ and 90 % $H_2O$ for 12 hours at 18 °C. Spectra correspond to : (1) 100 mM MeNC plus 100 mM 4-pentenal; (2) GMP; (3) GMP with 100 mM MeNC; (4) GMP with 100 mM 4-pentenal; (5) GMP with 100 mM MeNC and 100 mM 4-pentenal; (6) GMP with 100 mM MeNC and 100 mM 4-pentenal after heating at 95 °C for 15 minutes. Key resonances are highlighted, and the chemical-shift changes associated with N1 modification are marked with black rectangles.

**Figure S5.**

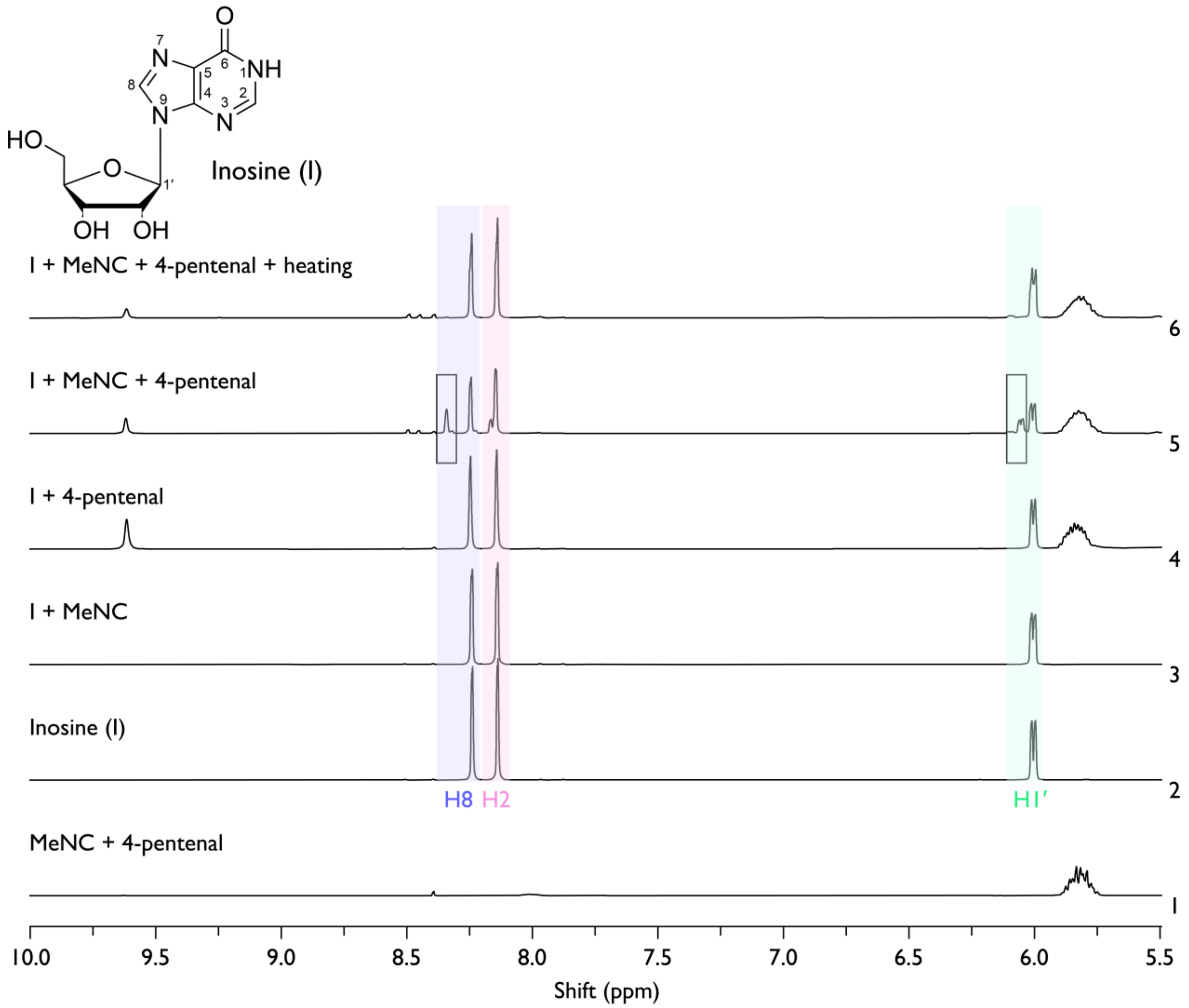


**Figure S5.** $^1$H NMR analysis of inosine (25 mM) incubated with methyl isocyanide (MeNC, 100 mM) and 4-pentenal (100 mM) in 200 mM HEPES, pH 8.0, in 10 % $D_2O$ and 90 % $H_2O$ for 12 hours at 18 °C. Spectra correspond to : (1) 100 mM MeNC plus 100 mM 4-pentenal; (2) inosine; (3) inosine with 100 mM MeNC; (4) inosine with 100 mM 4-pentenal; (5) inosine with 100 mM MeNC and 100 mM 4-pentenal; (6) inosine with 100 mM MeNC and 100 mM 4-pentenal after heating at 95 °C for 15 minutes. Key resonances are highlighted, and the chemical-shift changes associated with N1 modification are marked with black rectangles.

**Figure S6.**

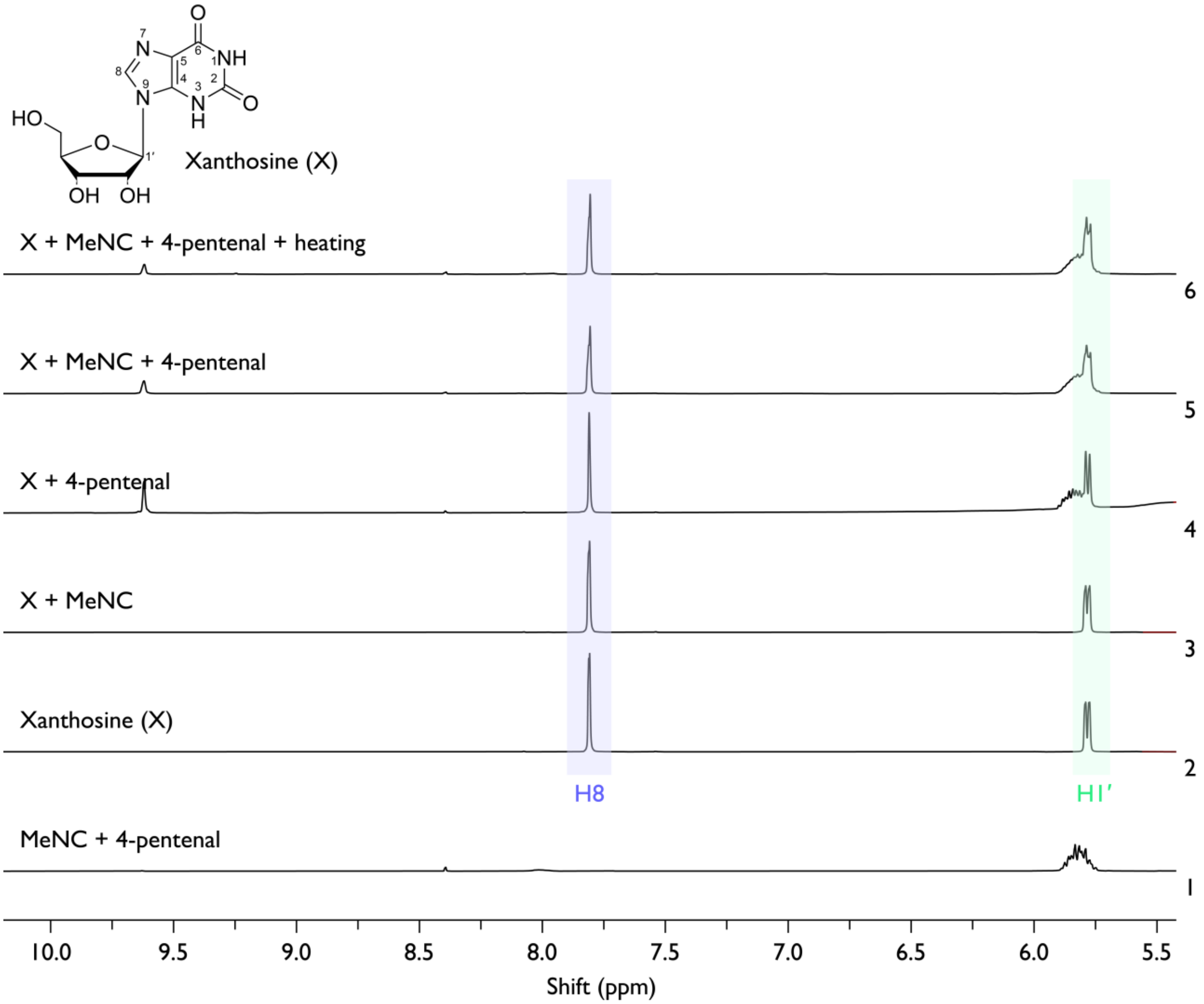


**Figure S6.** $^1$H NMR analysis of xanthosine (25 mM) incubated with MeNC (100 mM) and 4-pentenal (100 mM) in 200 mM HEPES, pH 8.0, containing 10 % $D_2O$ and 90 % $H_2O$ for 12 hours at 18 °C. Spectra correspond to: (1) 100 mM MeNC plus 100 mM 4-pentenal; (2) xanthosine; (3) xanthosine with 100 mM MeNC; (4) xanthosine with 100 mM 4-pentenal; (5) xanthosine with 100 mM MeNC and 100 mM 4-pentenal; (6) xanthosine with 100 mM MeNC and 100 mM 4-pentenal after heating at 95 °C for 15 minutes.

**Figure S7.**

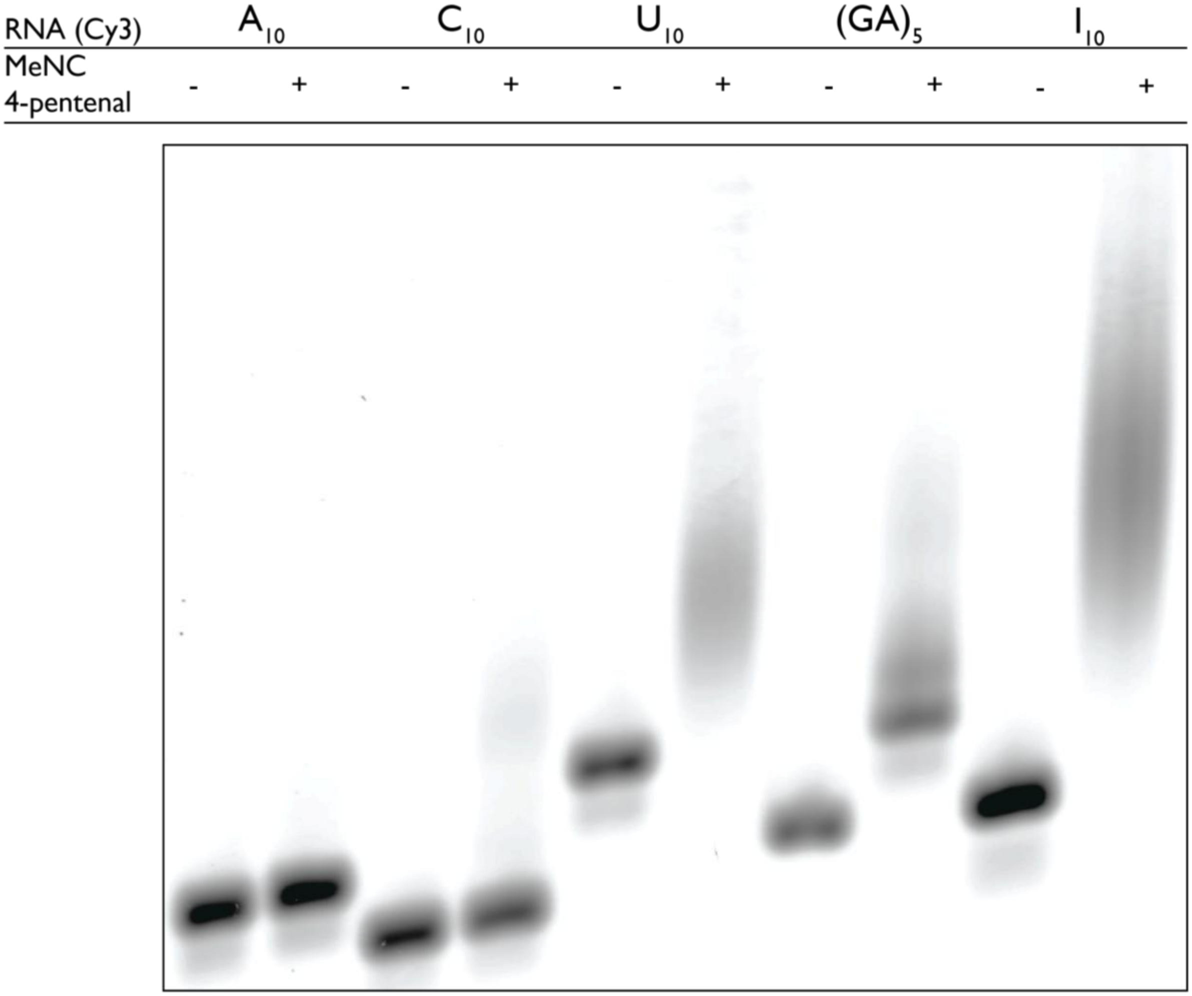


**Figure S7.** Electrophoretic mobility shift assay (EMSA) performed on 20 % (v/v) PAGE in 1× Tris–borate–EDTA (TBE). Reactions contained 3 μM of the indicated 3′-Cy3–labeled RNA tenmer, 200 mM MeNC (+), 200 mM 4-pentenal (+), and 200 mM HEPES, pH 8.0, incubated for 24 hours at 18 °C.

**Figure S8.**

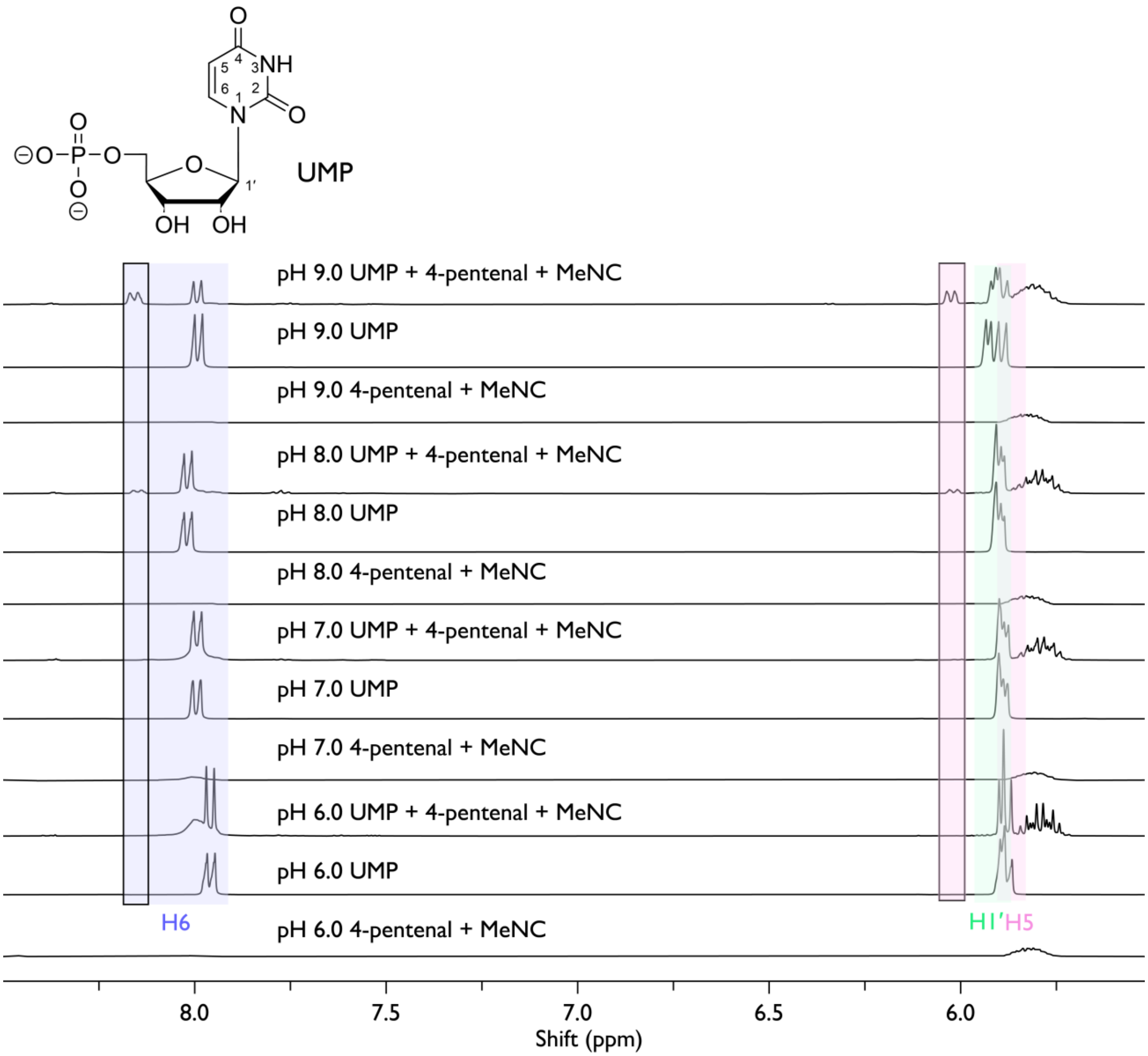


**Figure S8.** $^1$H NMR analysis of UMP (25 mM) incubated with MeNC (100 mM) and 4-pentenal (100 mM) in 200 mM HEPES at pH 6–9 (as indicated), containing 10 % $D_2O$ and 90 % $H_2O$, for 16 hours at 18 °C. Key resonances are highlighted, and the chemical-shift changes associated with N3 modification are marked with black rectangles.

**Figure S9.**

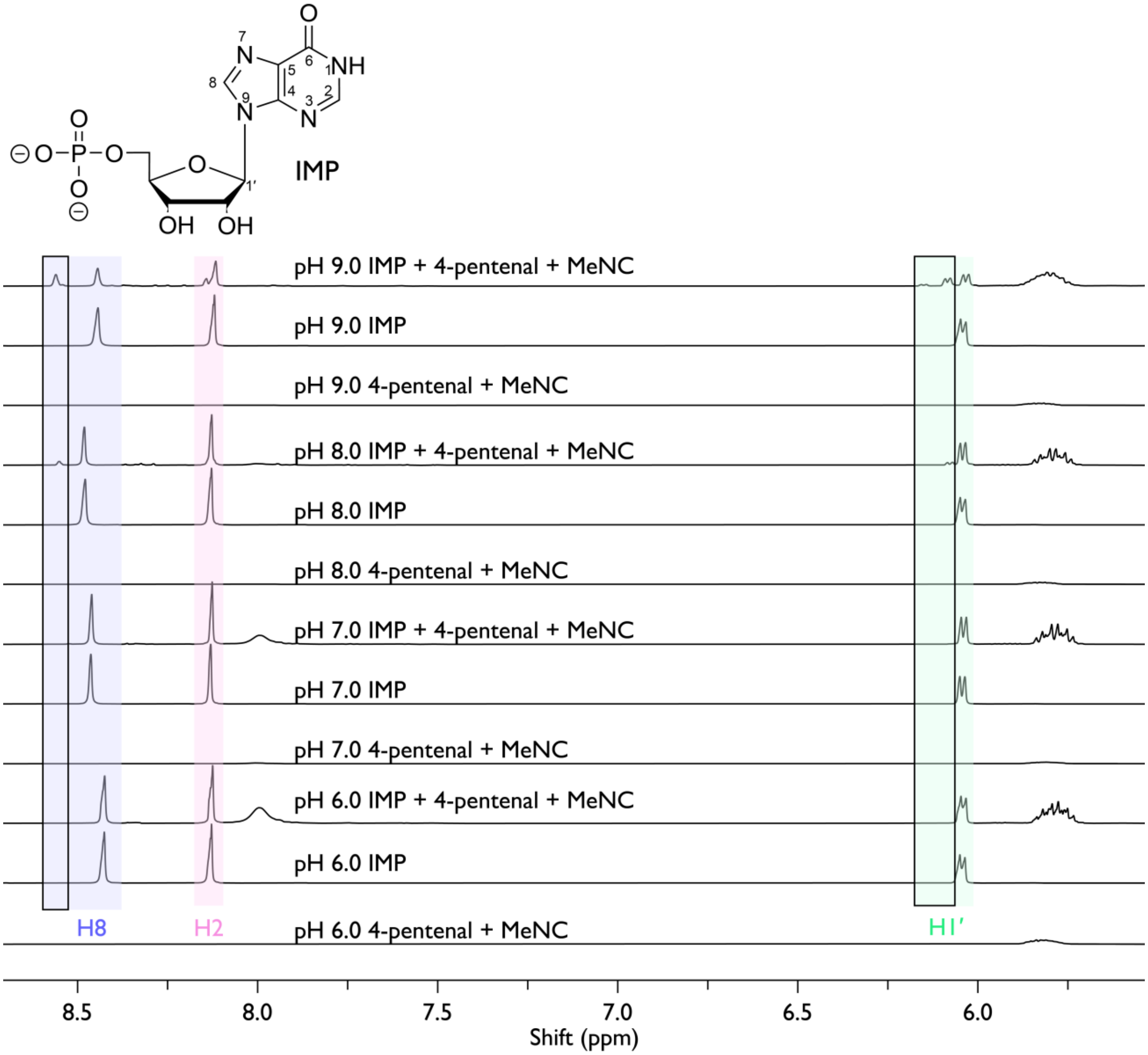


**Figure S9.** [1]H NMR analysis of inosine 5′-monophosphate (IMP, 25 mM) incubated with MeNC (100 mM) and 4-pentenal (100 mM) in 200 mM HEPES at pH 6–9 (as indicated), containing 10 % $D_2O$ and 90 % $H_2O$, for 16 hours at 18 °C. Key resonances are highlighted, and the chemical-shift changes associated with N3 modification are marked with black rectangles.

**Figure S10.**

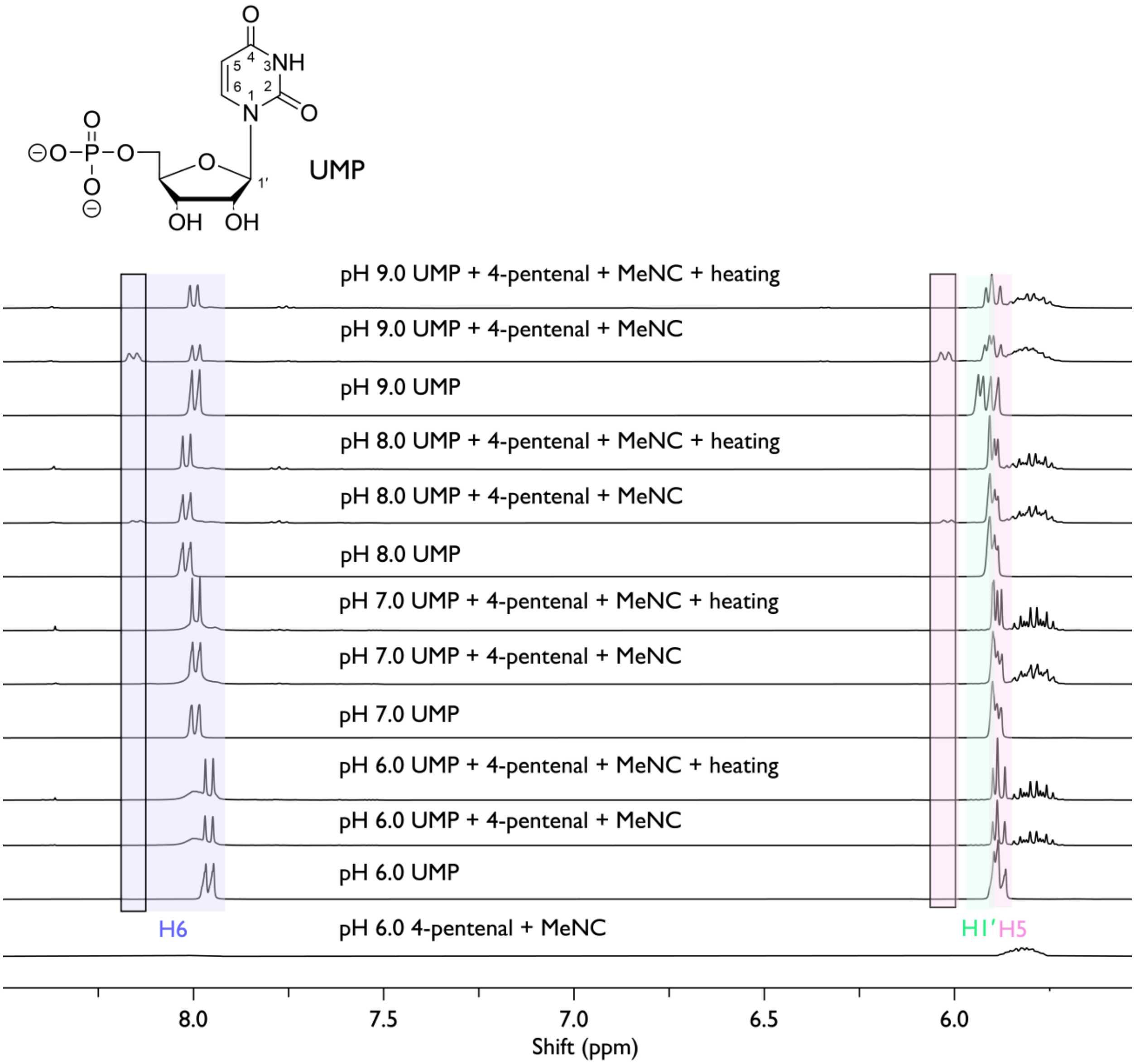


**Figure S10.** $^1$H NMR analysis of UMP (25 mM) incubated with MeNC (100 mM) and 4-pentenal (100 mM) in 200 mM HEPES at pH 6–9 (as indicated), containing 10 % $D_2O$ and 90 % $H_2O$, for 16 hours at 18 °C. Key resonances are highlighted, and the chemical-shift changes associated with N3 modification are marked with black rectangles. Heating was at 95 °C for 15 minutes.

**Figure S11.**

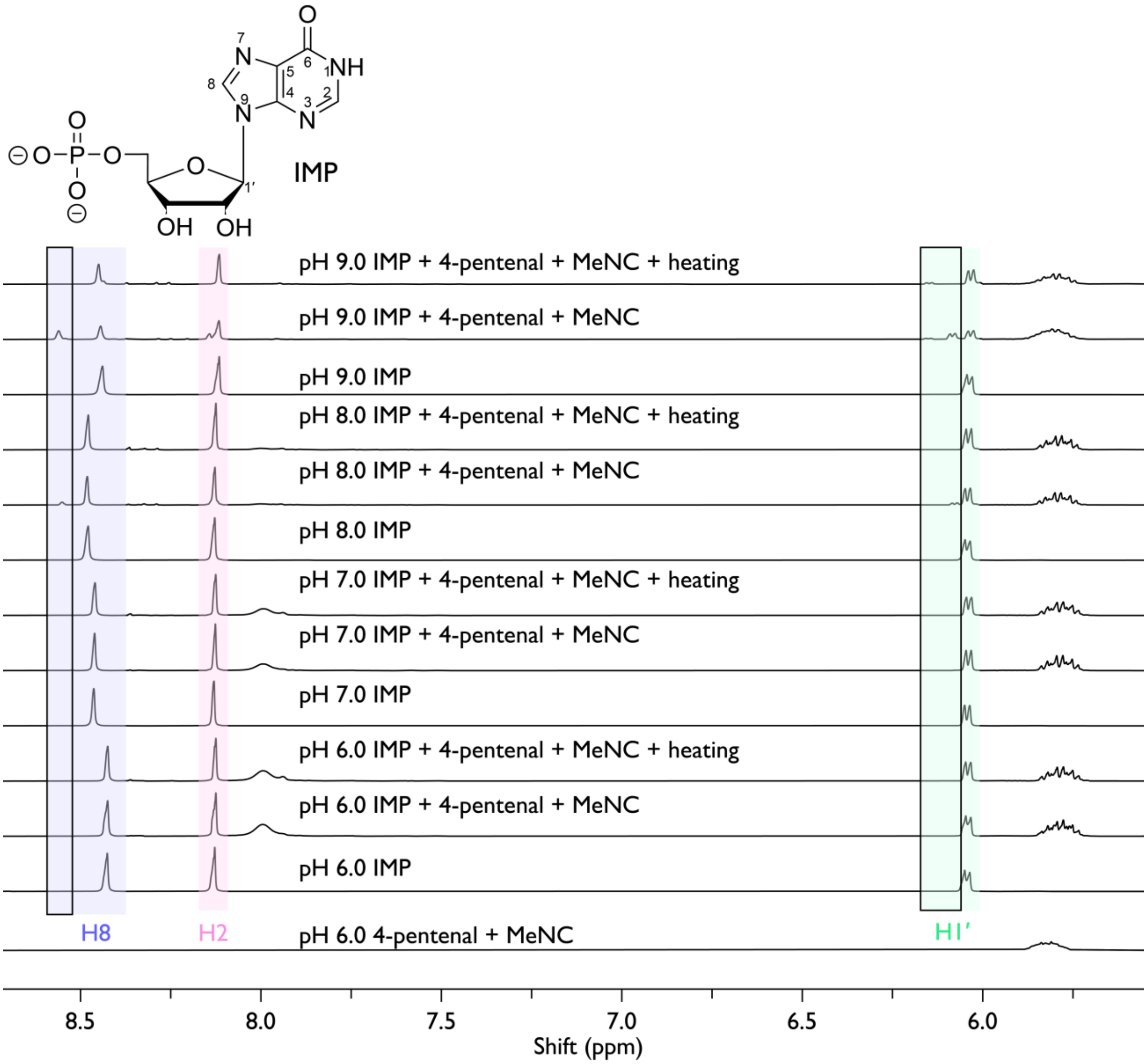


**Figure S11.** $^1$H NMR analysis of IMP (25 mM) incubated with MeNC (100 mM) and 4-pentenal (100 mM) in 200 mM HEPES at pH 6–9 (as indicated), containing 10 % $D_2O$ and 90 % $H_2O$, for 16 hours at 18 °C. Key resonances are highlighted, and the chemical-shift changes associated with N3 modification are marked with black rectangles. Heating was at 95 °C for 15 minutes.

**Figure S12.**

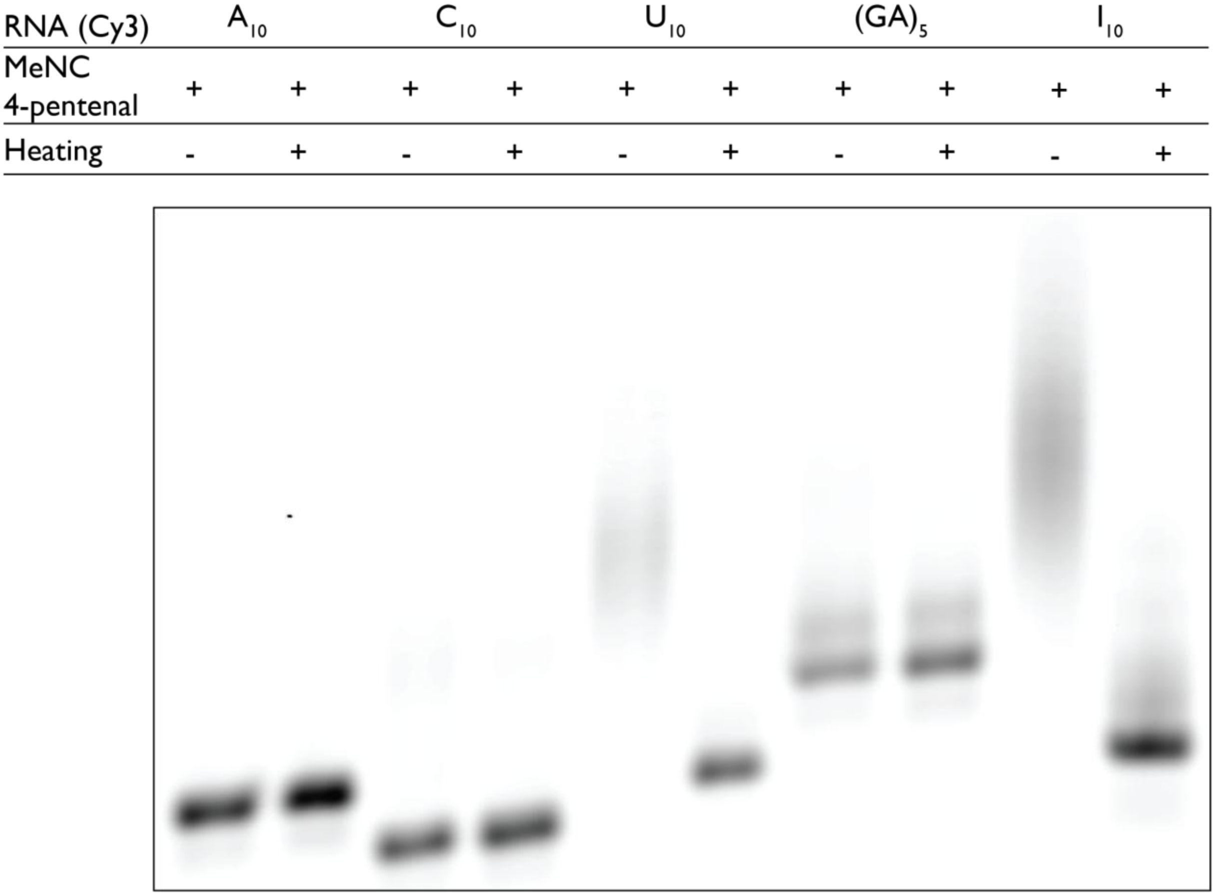


**Figure S12.** EMSA performed on 20 % PAGE in 1× TBE. Reactions contained 3 μM of the indicated 3′-Cy3–labeled RNA tenmer, 200 mM MeNC, 200 mM 4-pentenal, and 200 mM HEPES, pH 8.0, incubated for 24 hours at 18 °C. Heating (+) indicates treatment at 95 °C for 15 minutes.

**Figure S13.**

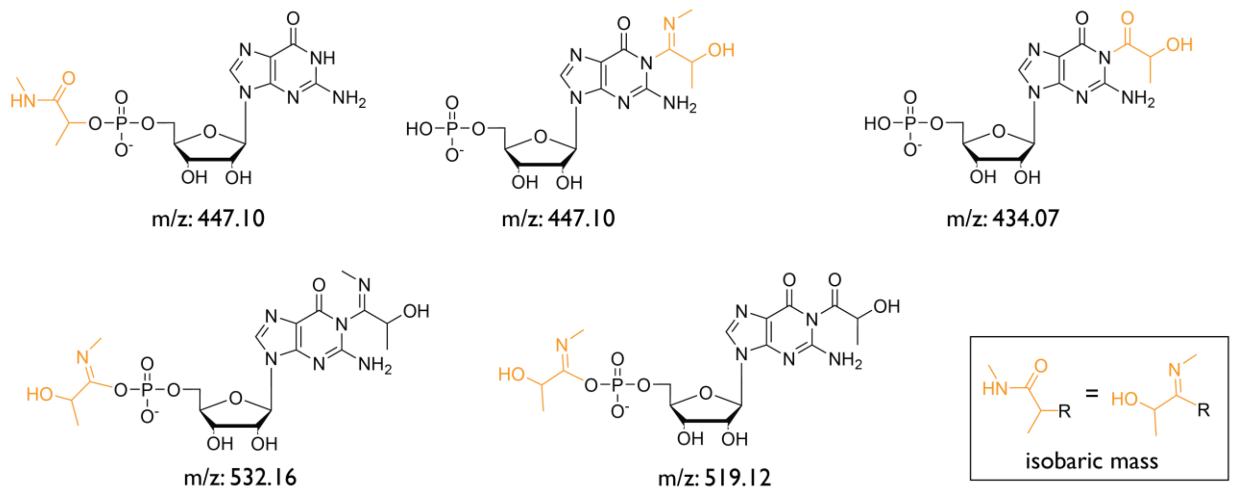


**Figure S13.** *Mass spectrometry analysis of GMP adducts formed in the presence of MeNC and acetaldehyde.* Reactions were carried out with 20 mM guanosine 5′-monophosphate (GMP), 200 mM methyl isonitrile, and 200 mM acetaldehyde in aqueous solution at pH 8.5. Prior to analysis, the reaction mixture was diluted to 50 µM GMP in methanol and analyzed by direct injection in negative ion mode. In addition to unmodified GMP, the major peaks corresponded to an N1-imidoyl adduct of GMP (m/z 447.10) and a phosphate-linked Passerini-type byproduct (also m/z 447.10), which are isobaric. The peak at m/z 447.10 is primarily attributed to the N1-imidoyl-GMP species. Within the first two hours, the dominant products were imidoyl-GMP (m/z 447.10) and its phosphoesterified form (m/z 532.16). After 24 hours, especially following heating at 95 °C for 15 minutes, a new class of adducts emerged that include an α-hydroxyacyl-GMP adduct (m/z 434.07) and its phosphate-linked form (m/z 519.12).

**Figure S14.**

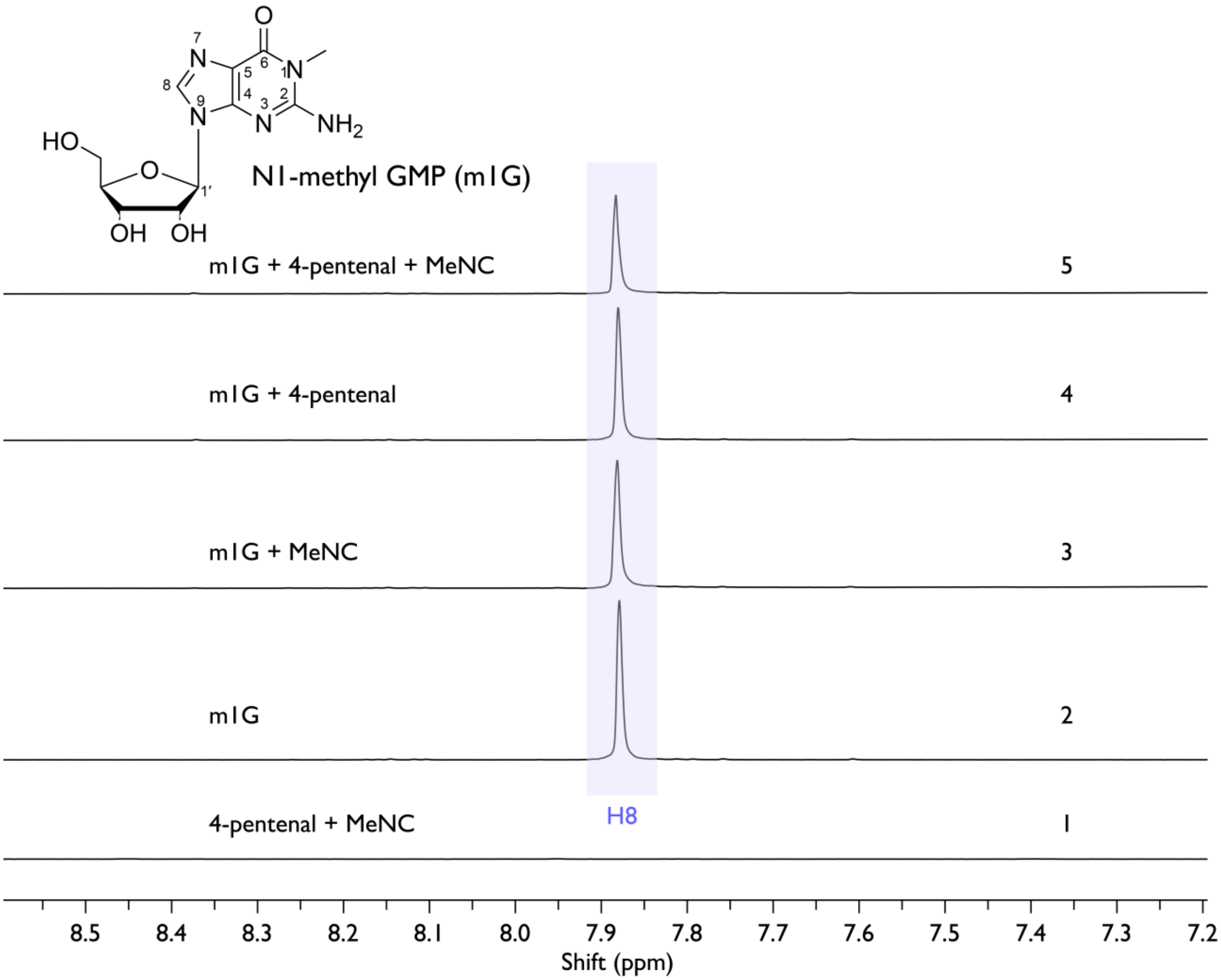


**Figure S14.** $^{1}$H NMR spectra showing lack of reaction of N1-methylguanosine (m1G) 25 mM, in the presence of 100 mM MeNC, 100 mM 4-pentenal, 200 mM HEPES pH 8.0 in 10 % (v/v) $D_2O$, 90 % (v/v) $H_2O$, incubated 16 h at room temperature. (1) 4-pentenal+MeNC, (2) m1G, (3) m1G+MeNC, (4) m1G+4-pentenal, (5) m1G+MeNC+4-pentenal.

**Figure S15.**

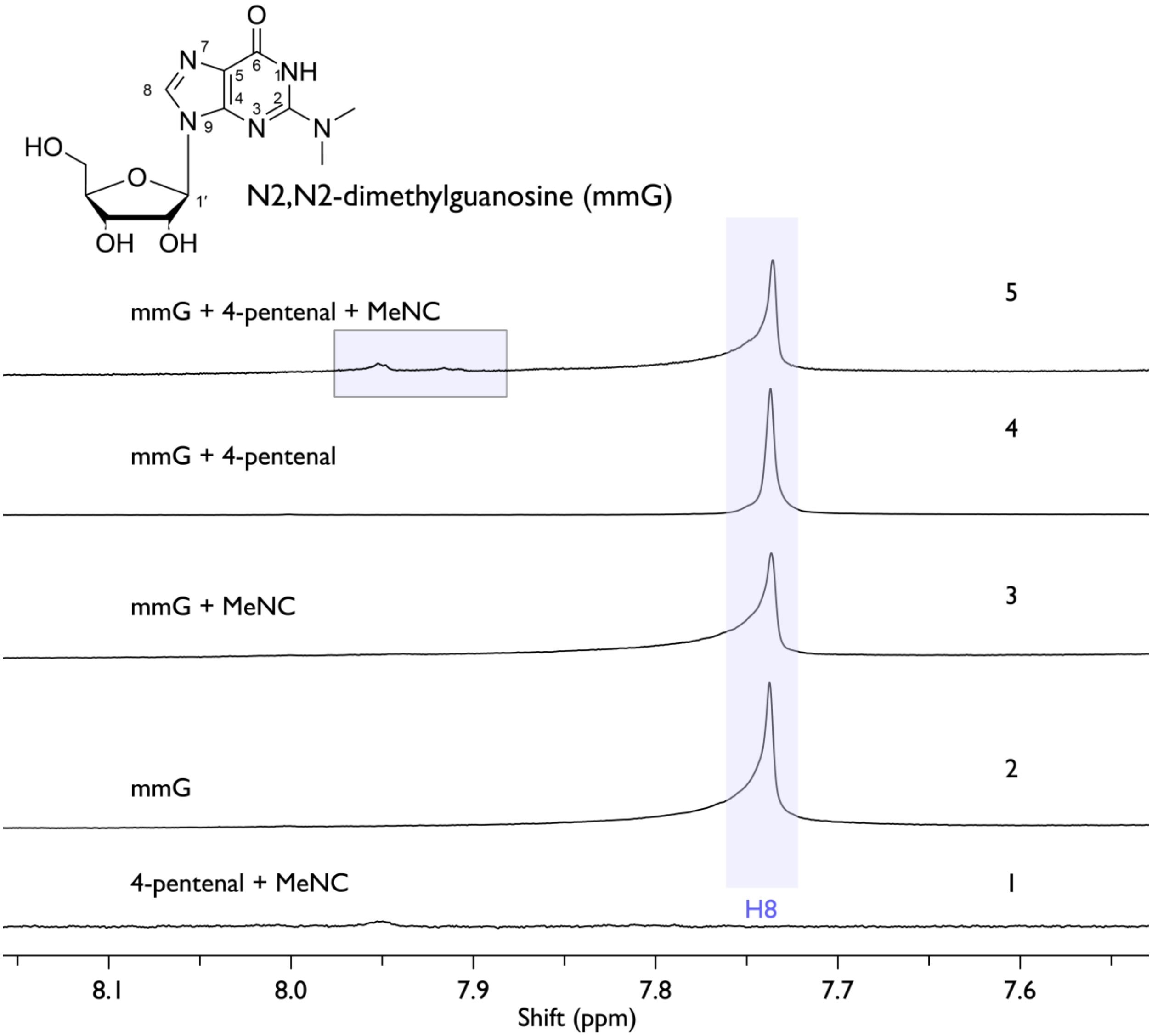


**Figure S15.** $^{1}$H NMR showing N2,N2-dimethylguanosine (mmG) 25 mM, 100 mM MeNC, 100 mM 4-pentenal, 200 mM HEPES pH 8.0 in 10 % (v/v) $D_2O$, 90 % (v/v) $H_2O$, incubated 16 h at room temperature. (1) mmG, (2) mmG+MeNC, (3) mmG+4-pentenal, (4) mmG+MeNC+4-pentenal.

**Figure S16.**

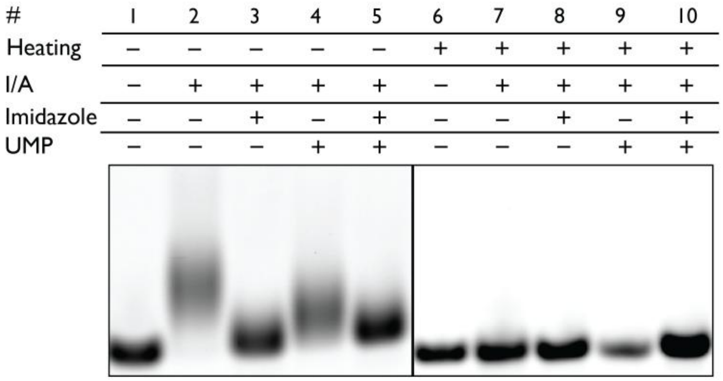


**Figure S16.** EMSA performed on 20% PAGE in 1× TBE. Reactions contained 2 μM $U_{10}$ 3′-Cy3–labeled RNA and 200 mM HEPES, pH 8.0, incubated for 24 h at 18 °C. Lane assignments are as follows: (1) $U_{10}$ alone; (2) $U_{10}$ + 200 mM MeNC + 200 mM 4-pentenal; (3) $U_{10}$ + 200 mM MeNC + 200 mM 4-pentenal + 200 mM imidazole; (4) $U_{10}$ + 200 mM MeNC + 200 mM 4-pentenal + 25 mM UMP; (5) $U_{10}$ + 200 mM MeNC + 200 mM 4-pentenal + 200 mM imidazole + 25 mM UMP. Lanes (6–10) correspond to conditions (1–5), respectively, following heating at 95 °C for 15 min.

**Figure S17.**

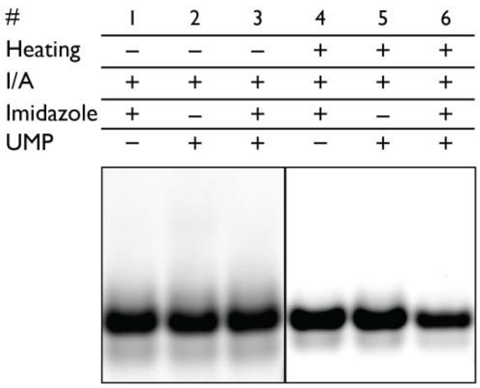


**Figure S17.** EMSA performed on 20% PAGE in 1× TBE. Reactions contained 2 μM $(GA)_5$ 3′-Cy3–labeled RNA and 200 mM HEPES, pH 8.0, incubated for 24 h at 18 °C. Lane assignments are as follows: (1) $(GA)_5$ + 200 mM MeNC + 200 mM 4-pentenal + 200 mM imidazole; (2) $(GA)_5$ + 200 mM MeNC + 200 mM 4-pentenal + 25 mM UMP; (3) $(GA)_5$ + 200 mM MeNC + 200 mM 4-pentenal + 200 mM imidazole + 25 mM UMP. Lanes (4–6) correspond to conditions (1–3), respectively, following heating at 95 °C for 15 min.

**Figure S18.**

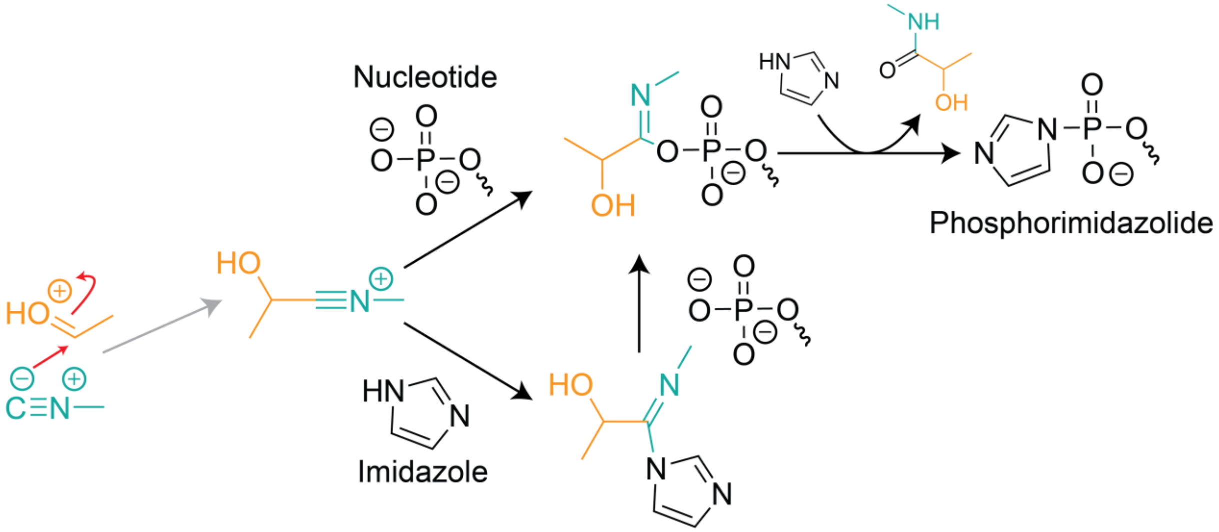


**Figure S18.** Proposed mechanism[3] for scavenging of the reactive nitrilium ion generated under isonitrile–aldehyde activation conditions by imidazole or phosphate. Nucleophilic interception of the nitrilium ion by imidazole yields an intermediate that facilitates formation of the phosphorimidazolide activated nucleotide, while interception by the phosphate of a nucleotide directly promotes phosphate activation. This pathway is suggested to reduce competing nucleobase modification by diverting reactive intermediates toward productive phosphate activation.

**Figure S19.**

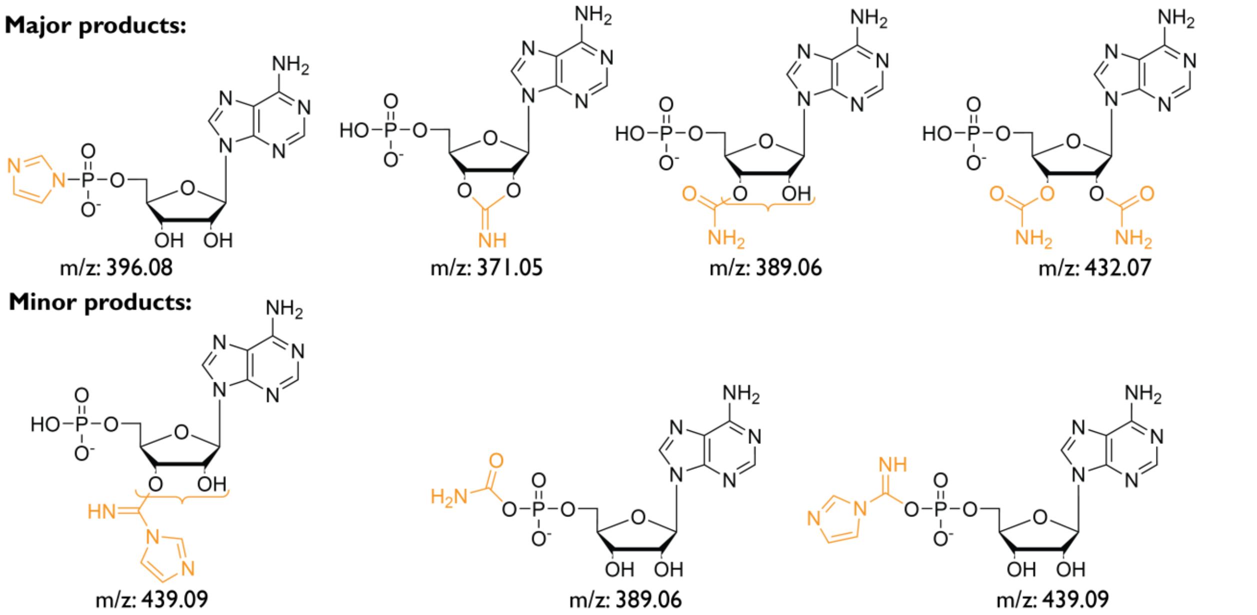


**Figure S19.** *Mass spectrometry of AMP adducts formed under acylimidazole type chemistry.* Reactions were carried out with 20 mM adenosine 5′-monophosphate (GMP), 200 mM IDI at pH 8.0. Prior to analysis, the reaction mixture was diluted to 50 μM AMP in methanol and analyzed by direct injection in negative ion mode. In addition to unmodified AMP, the major detected peaks corresponded to phosphorimidazolide (m/z 396.08), 2′,3′-cyclic imidoyl ester (m/z 371.05), 2′ or 3′-carbamoyl ester (m/z 389.06) and 2′,3′-dicarbamoyl ester (m/z 432.07). We also detected O-imidazolyl formimidate that can be either on 2′ or 3′ or phosphate (m/z 439.09), and carbamoyl phosphate (m/z 389.06).

**Figure S20.**

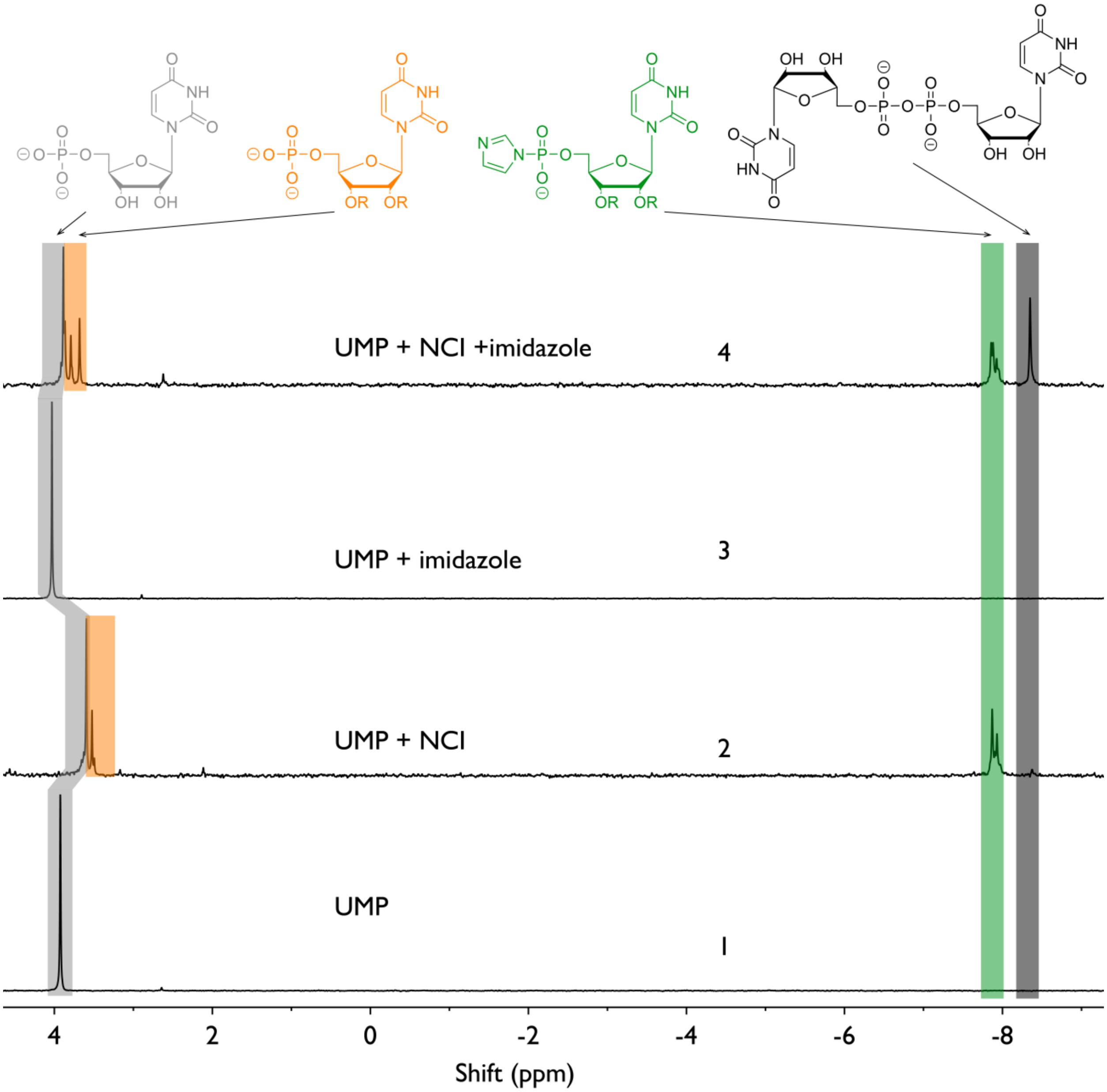


**Figure S20.** $^{31}$P NMR spectra of uridine 5′-monophosphate (UMP) reaction mixtures after 24 h in 10% (v/v) $D_2O$ and 90% (v/v) $H_2O$. Conditions: (1) 25 mM UMP; (4) 25 mM UMP + 200 mM NCI; (5) 25 mM UMP + 200 mM imidazole at pH 8.0; (7) 25 mM UMP + 200 mM NCI + 200 mM imidazole at pH 8.0. Peak assignments: grey, unmodified UMP; orange, 2′/3′-acylated UMP (Acyl-UMP); green, phosphorimidazolide intermediate (UMP-Im); black, UMP-pyrophosphate.

**Figure S21.**

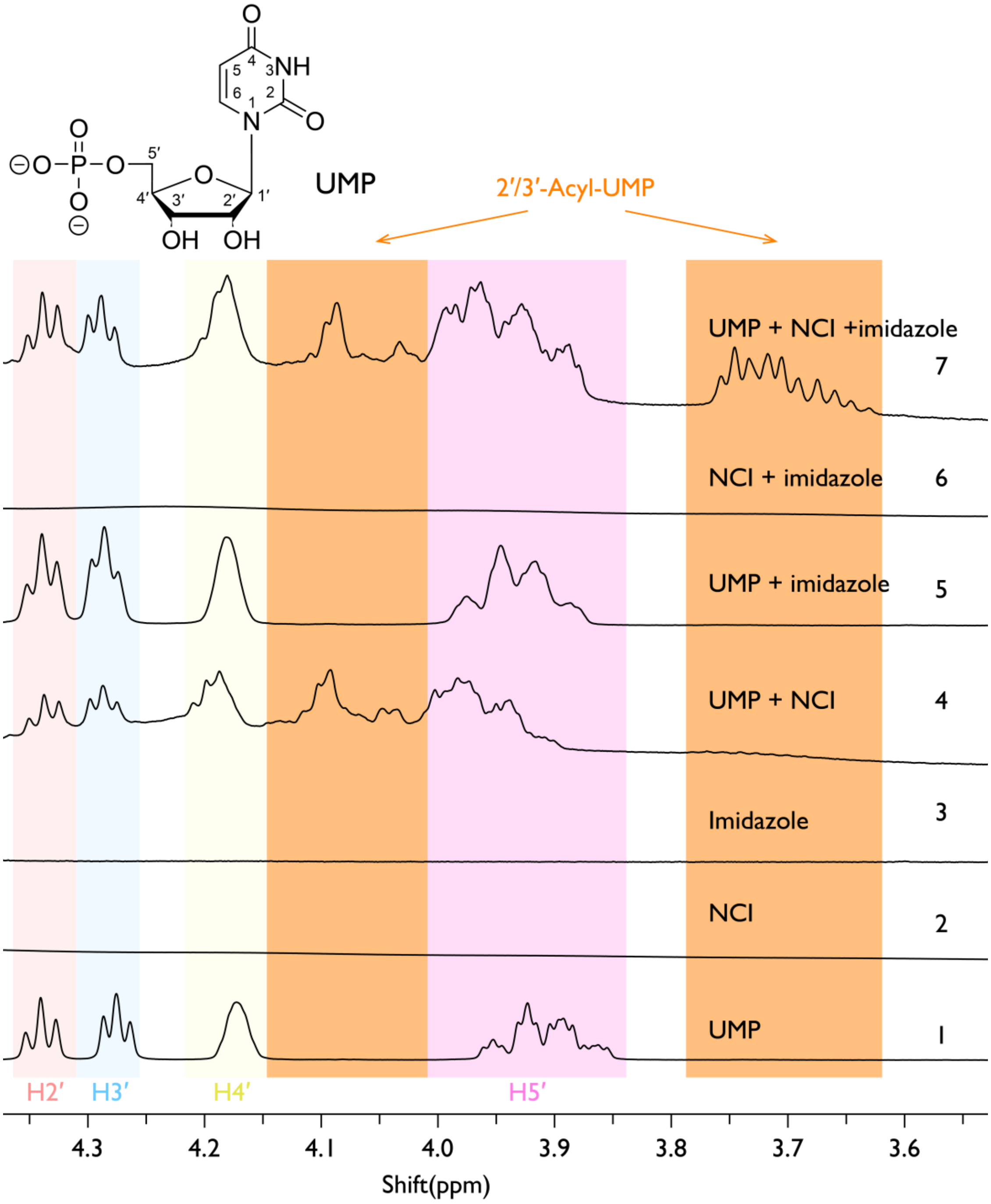


**Figure S21.** $^1$H NMR of UMP with NCI after 24 h in 10 % (v/v) $D_2O$, 90 % (v/v) $H_2O$. (1) 25 mM UMP; (2) 200 mM NCI; (3) 200 mM imidazole; (4) 25 mM UMP+200 mM NCI; (5) 25 mM UMP+200 mM imidazole; (6) 200 mM NCI + 200 mM imidazole; (7) 25 mM UMP+200 mM NCI+200 mM imidazole.

**Figure S22.**

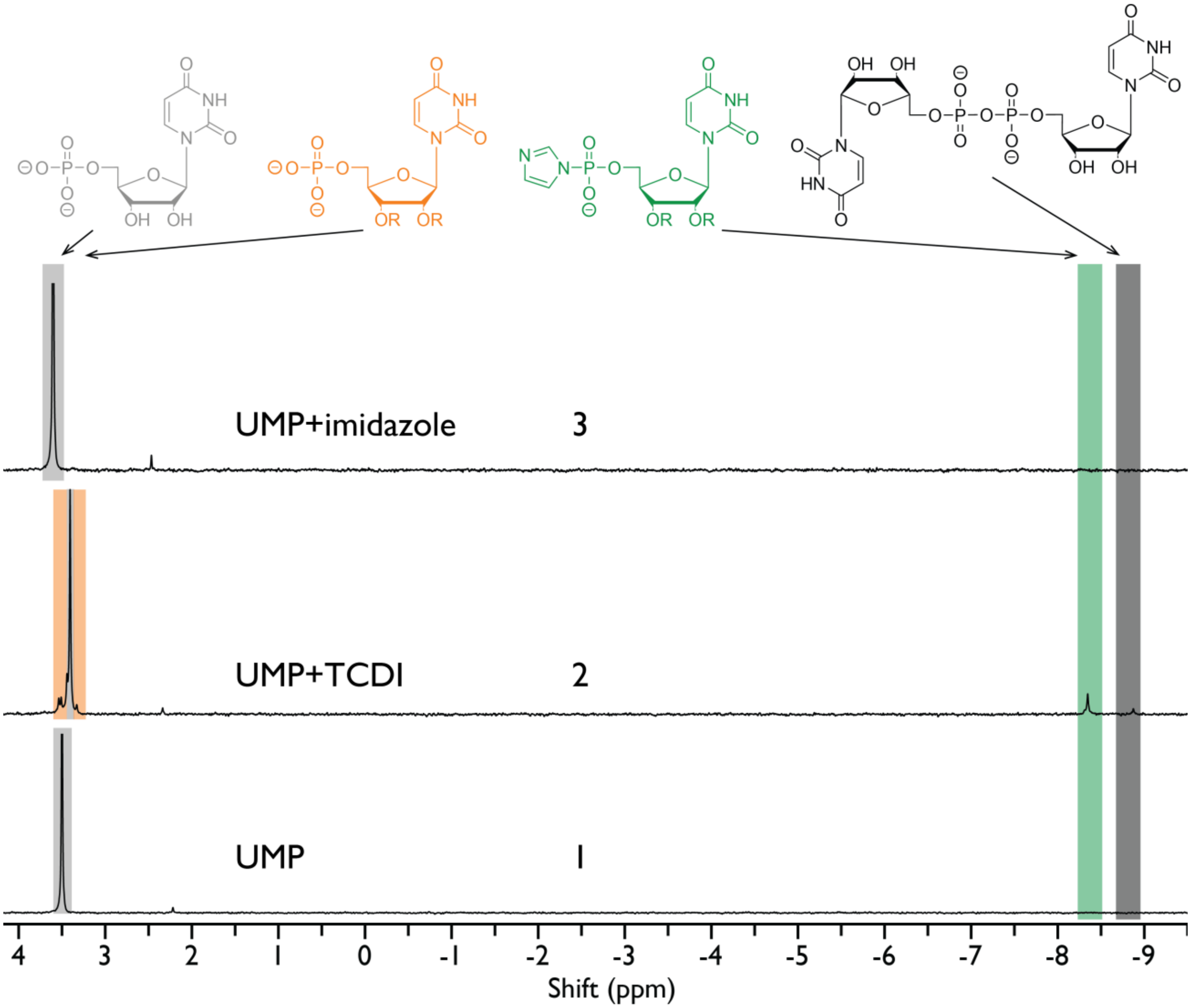


**Figure S22.** $^{31}P$ NMR of UMP with TCDI after 24 h in 10 % (v/v) $D_2O$, 90 % (v/v) $H_2O$. (1) 25 mM UMP; (2) 25 mM UMP+200 mM TCDI; (3) 25 mM UMP+200 mM imidazole.

**Figure S23.**

| # | 1 | 2 | 3 | 4 | 5 | 6 | 7 | 8 | 9 | 10 | 11 | 12 | 13 | 14 | 15 | 16 |
|---|---|---|---|---|---|---|---|---|---|---|---|---|---|---|---|---|
| $N_{10}$ | U | A | U | A | U | A | U | A | U | A | U | A | U | A | U | A |
| NCI | + | + | + | + | – | – | – | – | – | – | – | – | – | – | – | – |
| IDI | – | – | – | – | + | + | + | + | – | – | – | – | – | – | – | – |
| TCDI | – | – | – | – | – | – | – | – | + | + | + | + | – | – | – | – |
| pH | 8 | 8 | 6 | 6 | 8 | 8 | 6 | 6 | 8 | 8 | 6 | 6 | 8 | 8 | 6 | 6 |

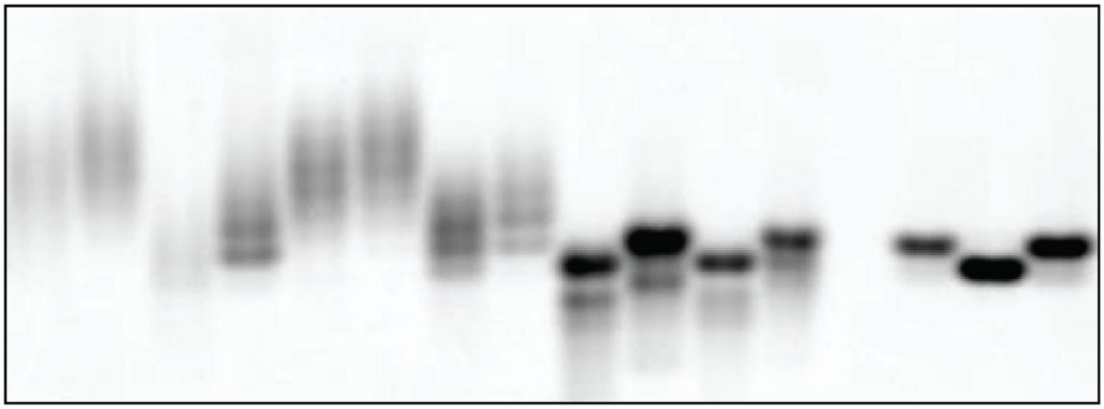

**Figure S23.** 20% (v/v) PAGE analysis of $U_{10}$-Cy3 and $A_{10}$-Cy3 oligonucleotides under acylimidazole activation conditions. Lanes 1, 3, 5, 7, 9, 11, 13, and 15 contain $U_{10}$-Cy3; lanes 2, 4, 6, 8, 10, 12, 14, and 16 contain $A_{10}$-Cy3. Samples in lanes 1, 2, 5, 6, 9, 10, 13, and 14 were incubated in 200 mM HEPES pH 8.0, and samples in lanes 3, 4, 7, 8, 11, 12, 15, and 16 were incubated in 200 mM HEPES pH 6.0. Activation reagents were: lanes 1–4, 200 mM NCI; lanes 5–8, 200 mM IDI; lanes 9–12, 200 mM TCDI; lanes 13–16, no activation reagent controls.

**Figure S24.**

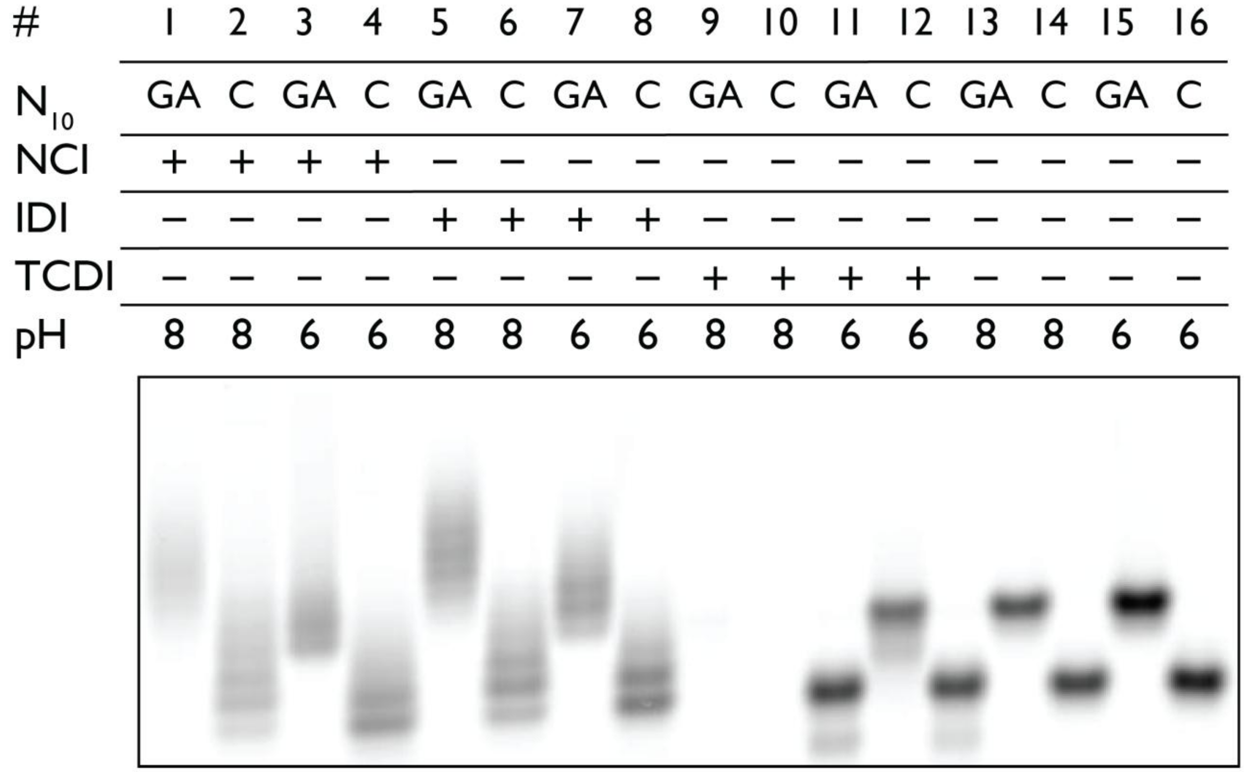


**Figure S24.** 20% (v/v) PAGE analysis of $(GA)_5$-Cy3 and $C_{10}$-Cy3 oligonucleotides under activation conditions. Lanes 1, 3, 5, 7, 9, 11, 13, and 15 contain $(GA)_5$-Cy3; lanes 2, 4, 6, 8, 10, 12, 14, and 16 contain $C_{10}$-Cy3. Samples in lanes 1, 2, 5, 6, 9, 10, 13, and 14 were incubated in 200 mM HEPES pH 8.0, and samples in lanes 3, 4, 7, 8, 11, 12, 15, and 16 were incubated in 200 mM HEPES pH 6.0. Activation reagents were applied as follows: lanes 1–4, NCI; lanes 5–8, IDI; lanes 9–12, TCDI; lanes 13–16, no activation reagent controls.

**Figure S25.**

| | $I_{10}$ | | | | | | | | | | | | | | | |
|---|---|---|---|---|---|---|---|---|---|---|---|---|---|---|---|---|
| # | 1 | 2 | 3 | 4 | 5 | 6 | 7 | 8 | 9 | 10 | 11 | 12 | 13 | 14 | 15 | 16 |
| Heat | − | − | − | − | − | − | − | − | + | + | + | + | + | + | + | + |
| NCI | + | + | − | − | − | − | − | − | + | + | − | − | − | − | − | − |
| IDI | − | − | + | + | − | − | − | − | − | − | + | + | − | − | − | − |
| TCDI | − | − | − | − | + | + | − | − | − | − | − | − | + | + | − | − |
| pH | 8 | 6 | 8 | 6 | 8 | 6 | 8 | 6 | 8 | 6 | 8 | 6 | 8 | 6 | 8 | 6 |

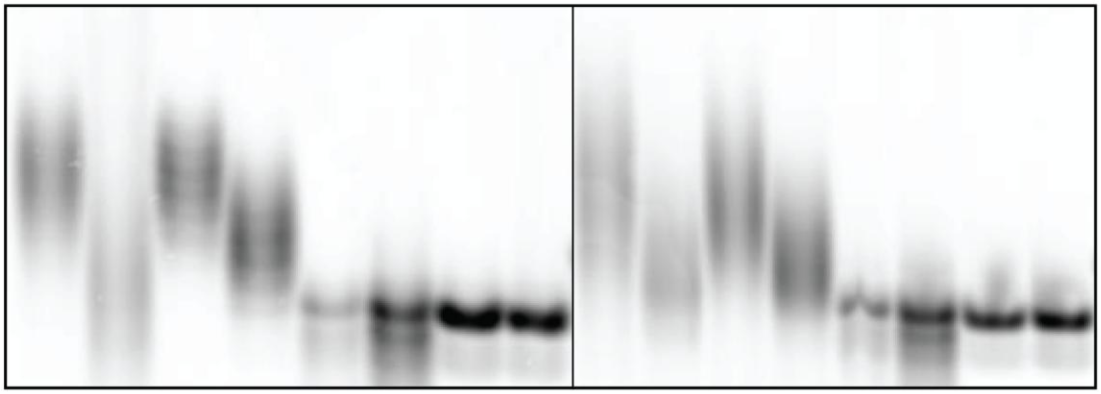

**Figure S25.** 20% (v/v) PAGE analysis of $dI_{10}$-Cy3 oligonucleotides incubated with activating agents, before (left) and after (right) heating at 95 °C for 30 min. Lane identities are identical between the two gels. Lanes 1, 3, 5, and 7 were incubated in 200 mM HEPES pH 8.0; lanes 2, 4, 6, and 8 were incubated in 200 mM HEPES pH 6.0. Activation reagents were applied as follows: lanes 1–2, NCI; lanes 3–4, IDI; lanes 5–6, TCDI; lanes 7–8, no activation reagent controls.

**Figure S26.**

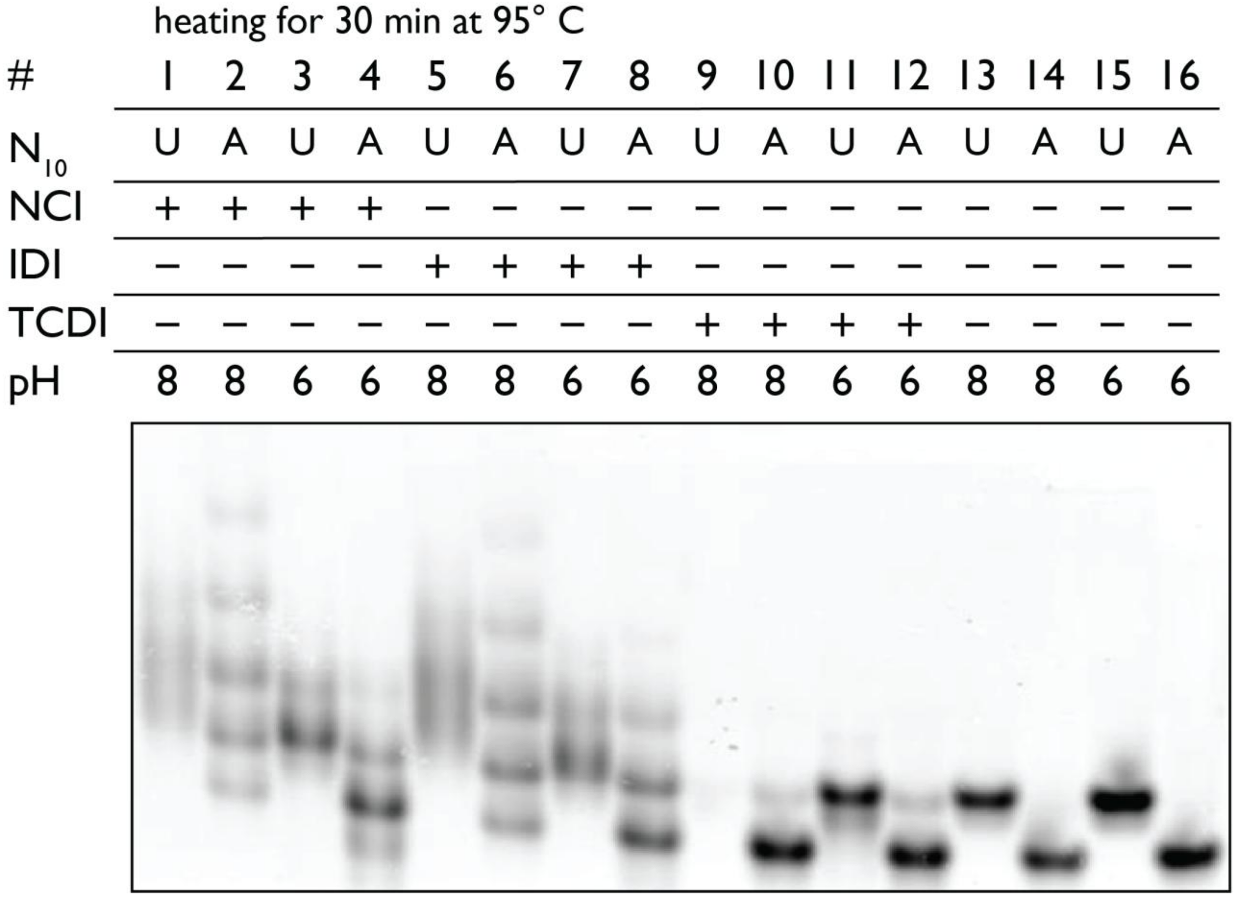


**Figure S26.** 20% (v/v) PAGE analysis of $U_{10}$-Cy3 and $A_{10}$-Cy3 oligonucleotides after activation reactions followed by heating at 95 °C for 30 min. Lanes 1, 3, 5, 7, 9, 11, 13, and 15 contain $U_{10}$-Cy3; lanes 2, 4, 6, 8, 10, 12, 14, and 16 contain $A_{10}$-Cy3. Samples in lanes 1, 2, 5, 6, 9, 10, 13, and 14 were incubated in 200 mM HEPES pH 8.0, and samples in lanes 3, 4, 7, 8, 11, 12, 15, and 16 were incubated in 200 mM HEPES pH 6.0. Activation reagents were applied as follows: lanes 1–4, NCI; lanes 5–8, IDI; lanes 9–12, TCDI; lanes 13–16, no activation reagent controls.

**Figure S27.**

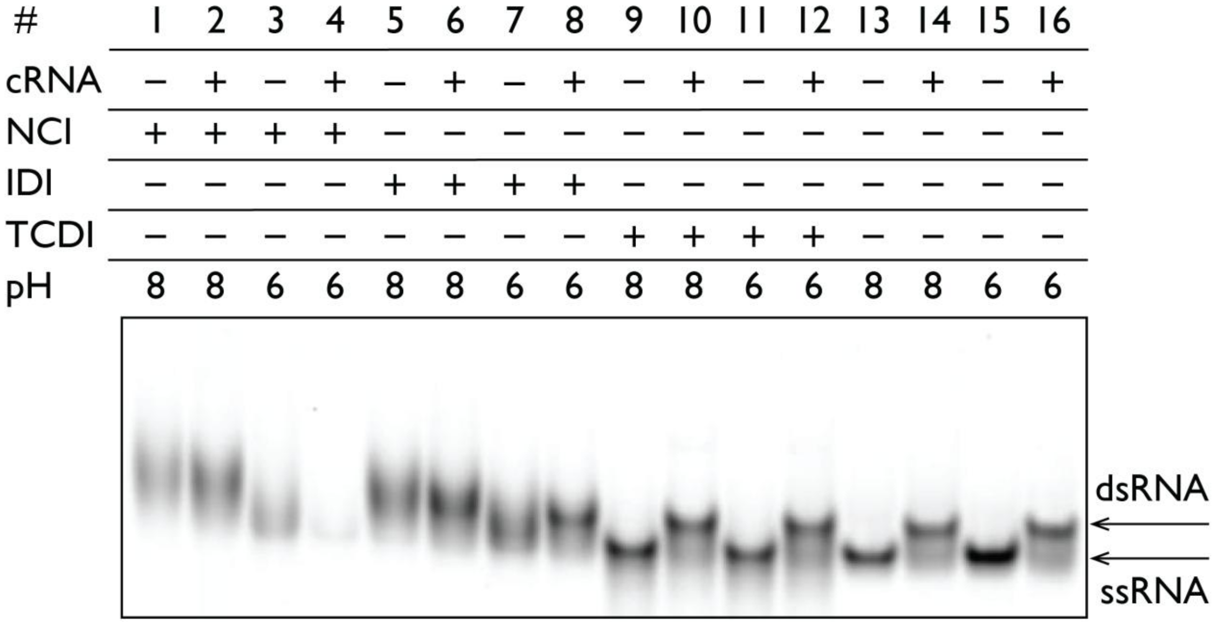


**Figure S27.** 20% (v/v) PAGE analysis of oligonucleotide G-001 incubated with either single-stranded (ss) or duplex (ds) $N_{12}$-Cy3 for 24 h at room temperature. Lanes 1, 3, 5, 7, 9, 11, 13, and 15 contain ssRNA; lanes 2, 4, 6, 8, 10, 12, 14, and 16 contain dsRNA. Samples in lanes 1, 2, 5, 6, 9, 10, 13, and 14 were incubated in 200 mM HEPES pH 8.0; samples in lanes 3, 4, 7, 8, 11, 12, 15, and 16 were incubated in 200 mM HEPES pH 6.0. Activation reagents were applied as follows: lanes 1–4, NCI; lanes 5–8, IDI; lanes 9–12, TCDI; lanes 13–16, no activation reagent controls.

**Figure S28.**

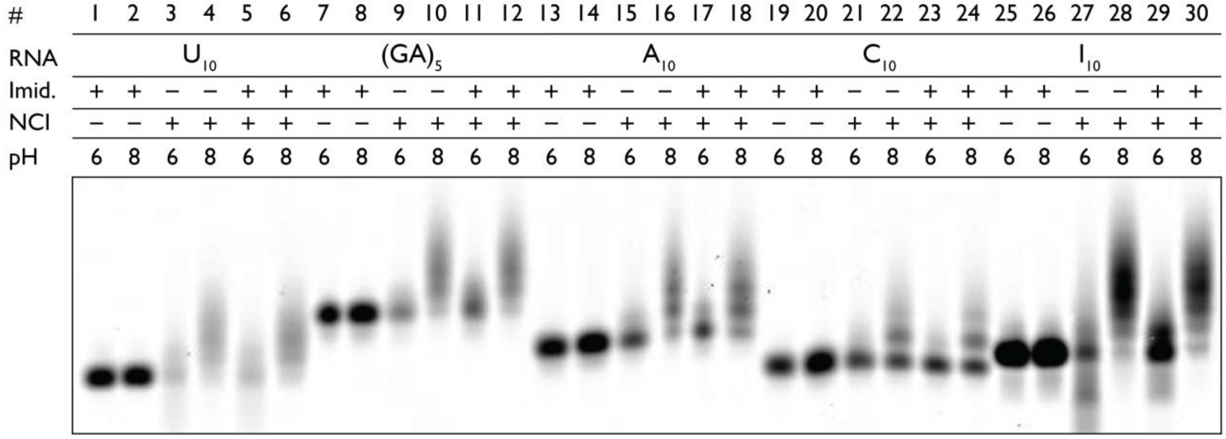


**Figure S28.** 20% (v/v) PAGE analysis of $N_{10}$-Cy3 oligonucleotides incubated with activation reagents for 24 h at room temperature. Lanes 1–6 contain $U_{10}$; lanes 7–12 contain $(GA)_5$; lanes 13–18 contain $A_{10}$; lanes 19–24 contain $C_{10}$; lanes 25–30 contain $dI_{10}$. Samples in lanes 1, 3, 5, 7, 9, 11, 13, 15, 17, 19, 21, 23, 25, 27, and 29 were incubated in 200 mM MES pH 6.0; samples in lanes 2, 4, 6, 8, 10, 12, 14, 16, 18, 20, 22, 24, 26, 28, and 30 were incubated in 200 mM HEPES pH 8.0. Activation reagents were applied as follows: lanes 3–6, 9–12, 15–18, 21–24, and 27–30 contained 50 mM NCI; lanes 1, 2, 5, 6, 7, 8, 11, 12, 13, 14, 17, 18, 19, 20, 23, 24, 25, 26, 29, and 30 contained 50 mM imidazole.

**Figure S29.**

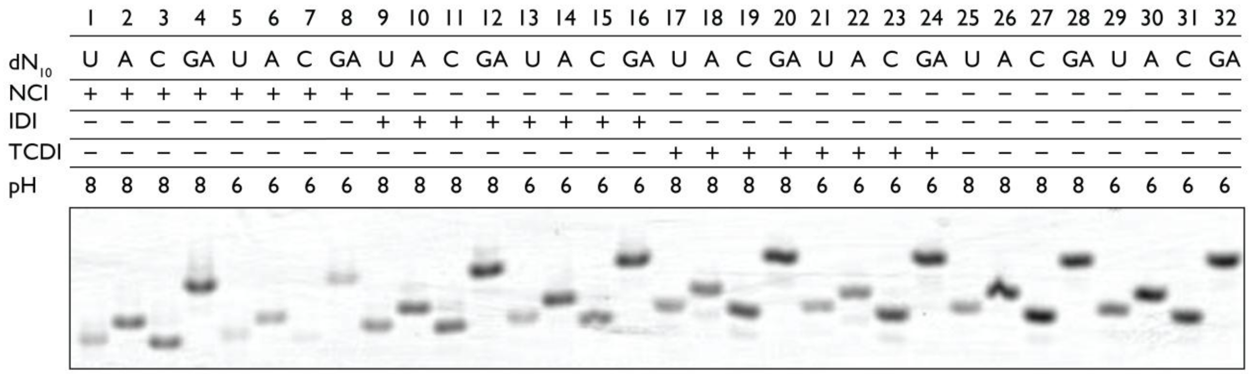


**Figure S29.** 20% (v/v) PAGE analysis of DNA oligonucleotides ($dN_{10}$-Cy3) incubated with activation reagents for 24 h at room temperature. Lanes 1–8 contain samples treated with NCI; lanes 9–16 with IDI; lanes 17–24 with TCDI; lanes 25–32 are controls. Samples in lanes 5–8, 13–16, 21–24, and 29–32 were incubated in 200 mM HEPES pH 6.0. Lane assignments for each DNA sequence are as follows: lanes 1, 5, 9, 13, 17, 21, 25, and 29 contain $dU_{10}$; lanes 2, 6, 10, 14, 18, 22, 26, and 30 contain $dA_{10}$; lanes 3, 7, 11, 15, 19, 23, 27, and 31 contain $dC_{10}$; lanes 4, 8, 12, 16, 20, 24, 28, and 32 contain $dG_{10}$.

**Figure S30.**

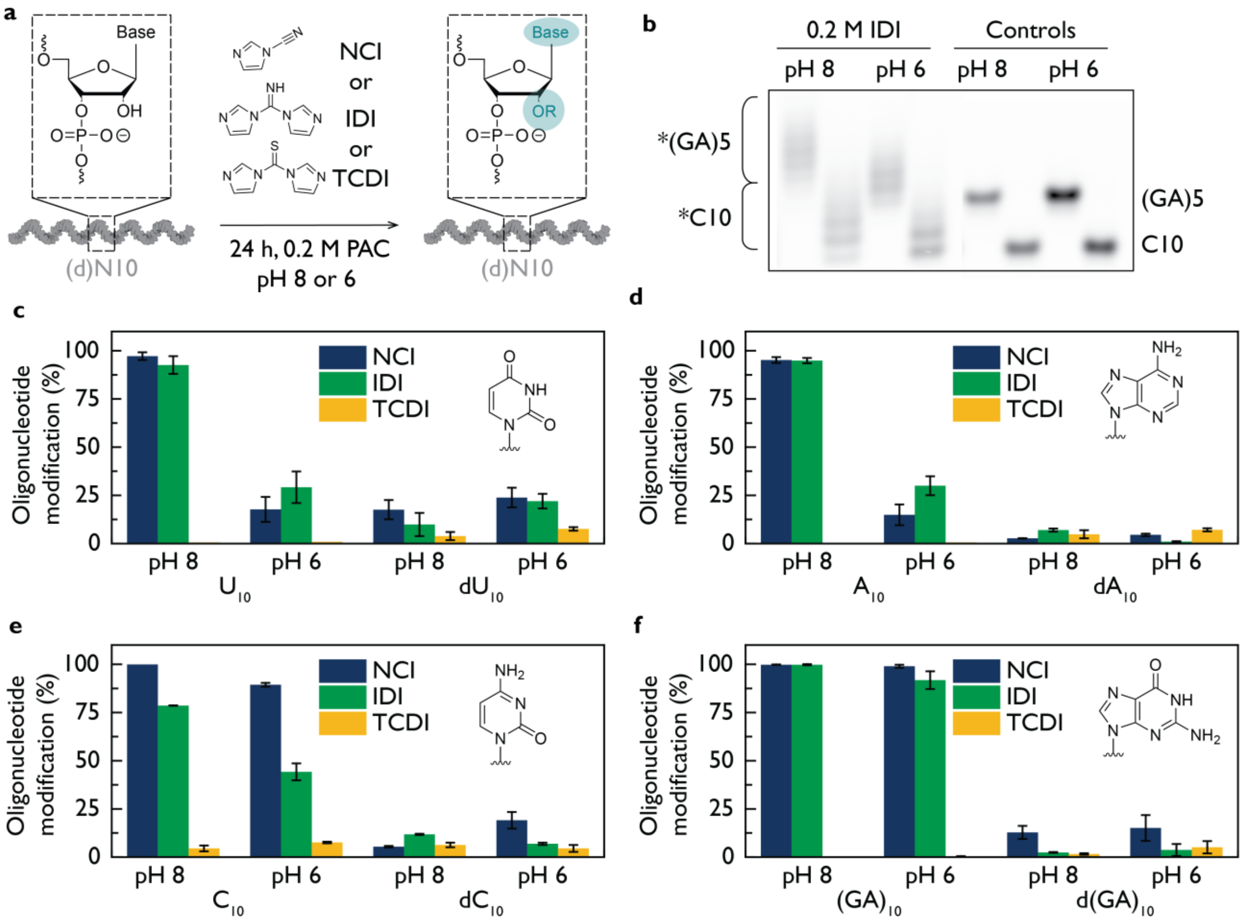


**Figure S30.** Active imidazoles modify oligonucleotides through nucleobase-specific reactions and 2′-OH acylation. **(a)** Design of RNA and DNA 10-mer oligonucleotides bearing a 3′-Cy3 label for PAGE analysis. Reactions contained 1 μM oligonucleotide, 0.2 M activation reagent, and 0.2 M HEPES at pH 8.0 or pH 6.0. The activation reagents tested were NCI, IDI, and TCDI. **(b)** Representative 20% (v/v) PAGE gel of $(GA)_5$ and $C_{10}$ RNA oligonucleotides treated with IDI, showing more extensive modification at pH 8.0 than at pH 6.0. **(c–f)** Summary of modification patterns across RNA and DNA sequences. At pH 8.0, both NCI and IDI modify all nucleobases, with substantially reduced modification at pH 6.0 or when deoxyoligonucleotides are used. TCDI produces only limited S-acylation of both RNA and DNA. *The pH 6.0 reactions were prepared at the indicated pH, but imidazole released during activation increased the final pH to approximately 7.5 ± 0.15 by the end of the incubation.

**Figure S31.**

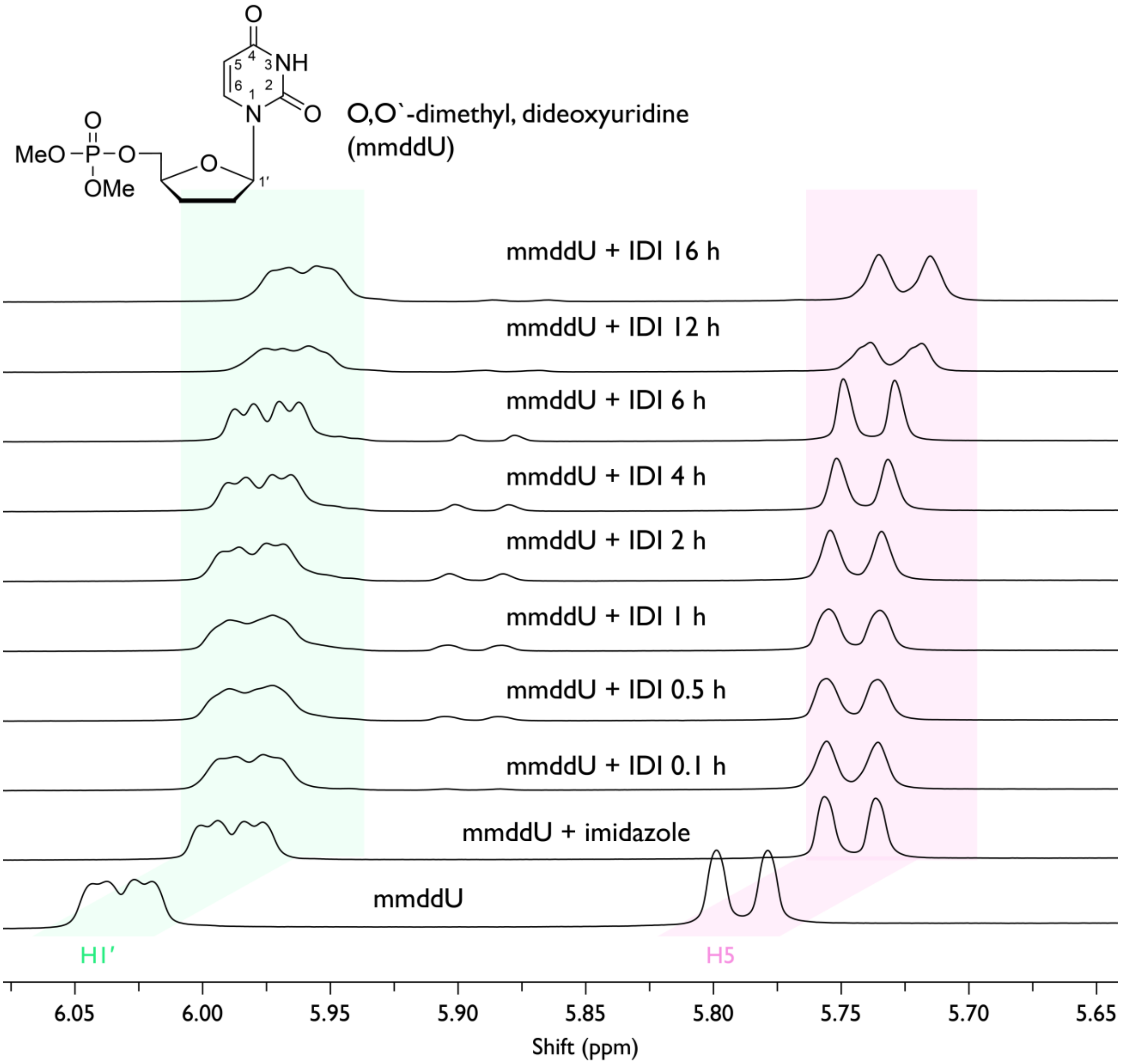


**Figure S31.** $^1$H NMR spectra of 25 mM mmddU incubated with 200 mM IDI in 10 % (v/v) $D_2O$ / 90 % (v/v) $H_2O$ over the course of the reaction. Progressive shifts in the highlighted proton resonances arise from the increase in pH caused by imidazole release due to IDI hydrolysis.

**Figure S32.**

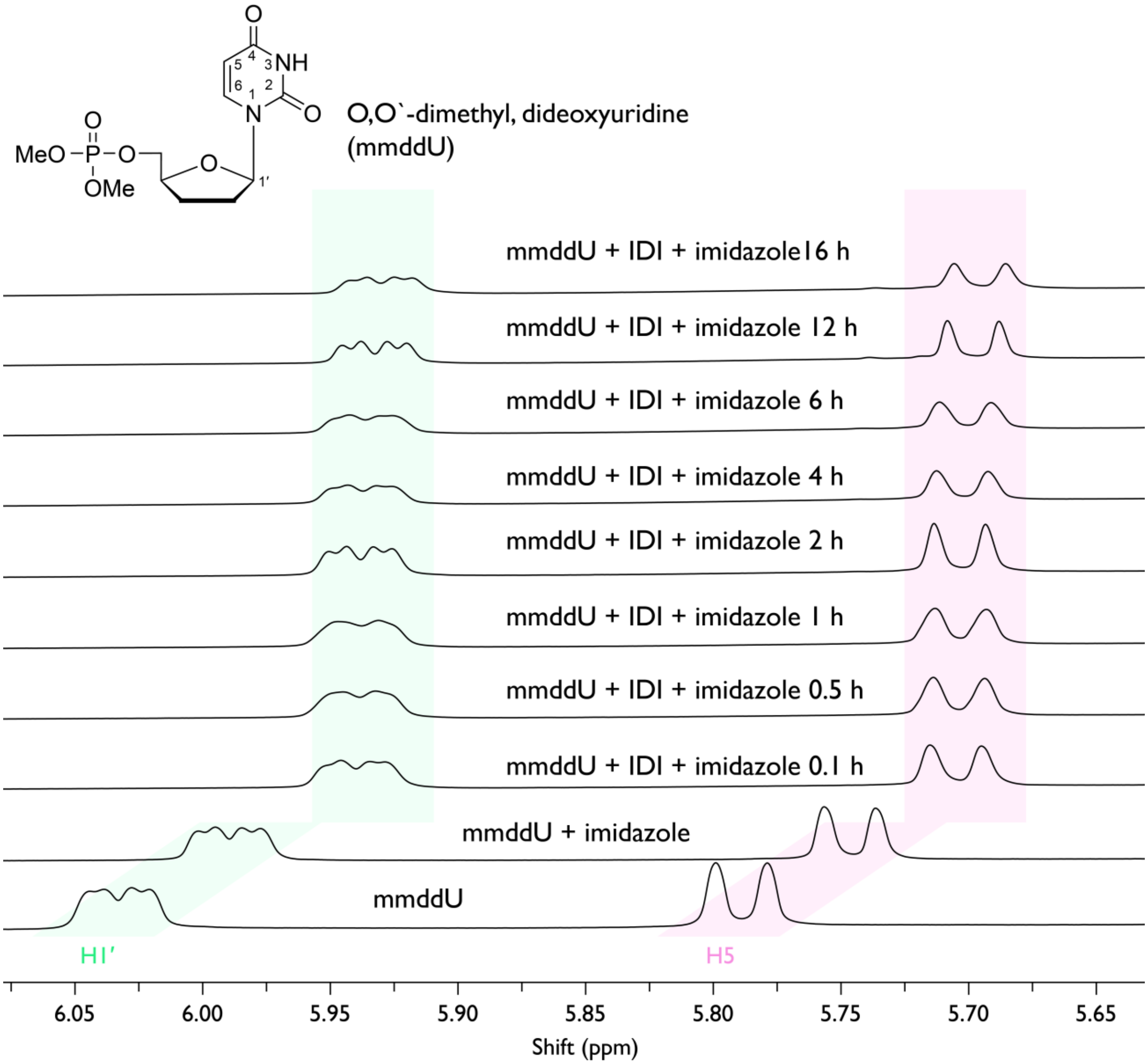


**Figure S32.** $^{1}H$ NMR spectra of mmddU (25 mM) incubated with 200 mM IDI and 200 mM imidazole in 10 % (v/v) $D_2O$ and 90 % (v/v) $H_2O$ over the course of the reaction. Progressive shifts in the highlighted proton resonances arise from the increase in pH caused by imidazole release due to IDI hydrolysis.

**Figure S33.**

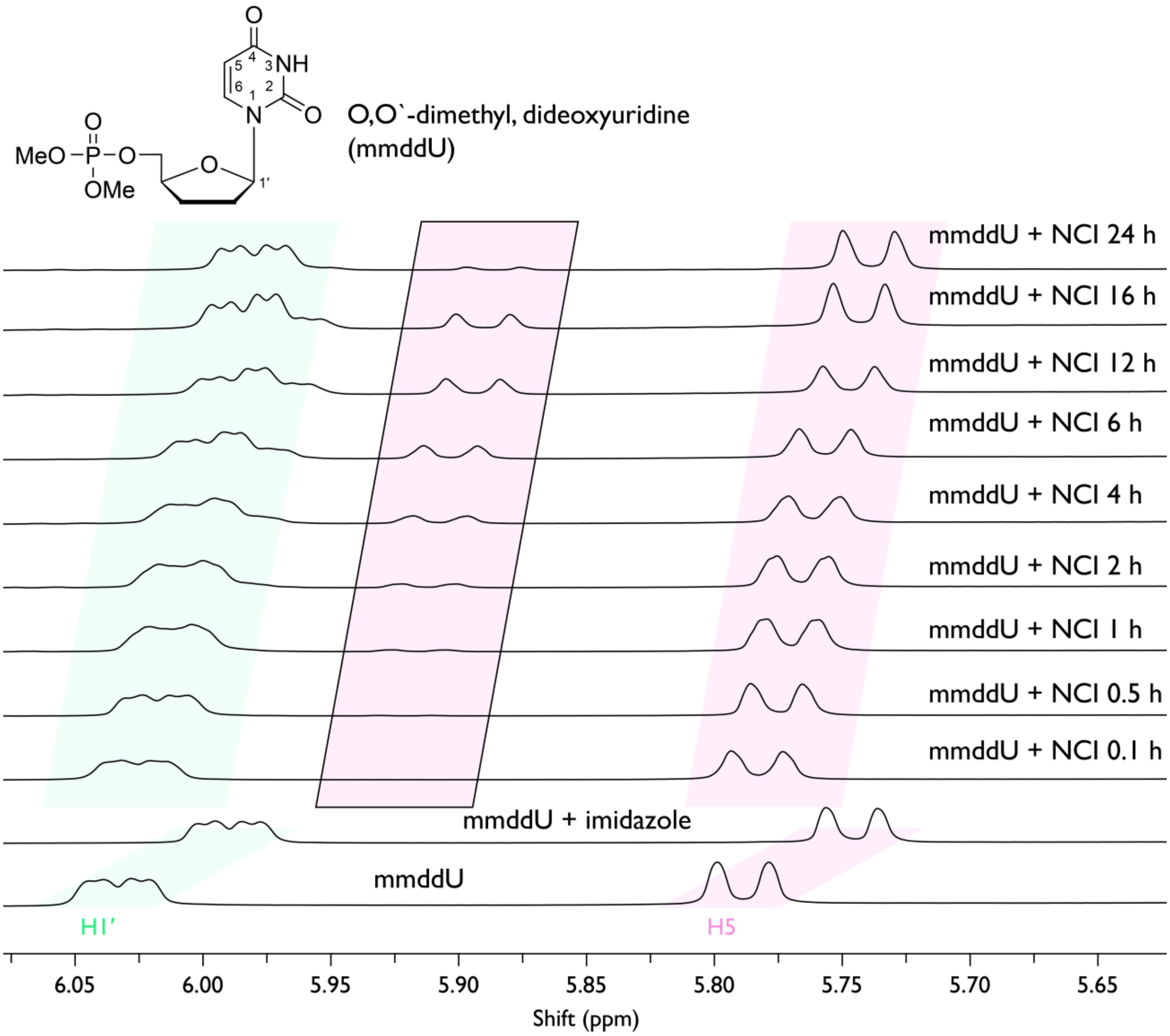


**Figure S33.** $^{1}H$ NMR spectra of mmddU (25 mM) incubated with 200 mM NCI in 10 % (v/v) $D_2O$ and 90 % (v/v) $H_2O$ over the course of the reaction. Progressive shifts in the highlighted proton resonances arise from the increase in pH caused by imidazole release due to IDI hydrolysis.

**Figure S34.**

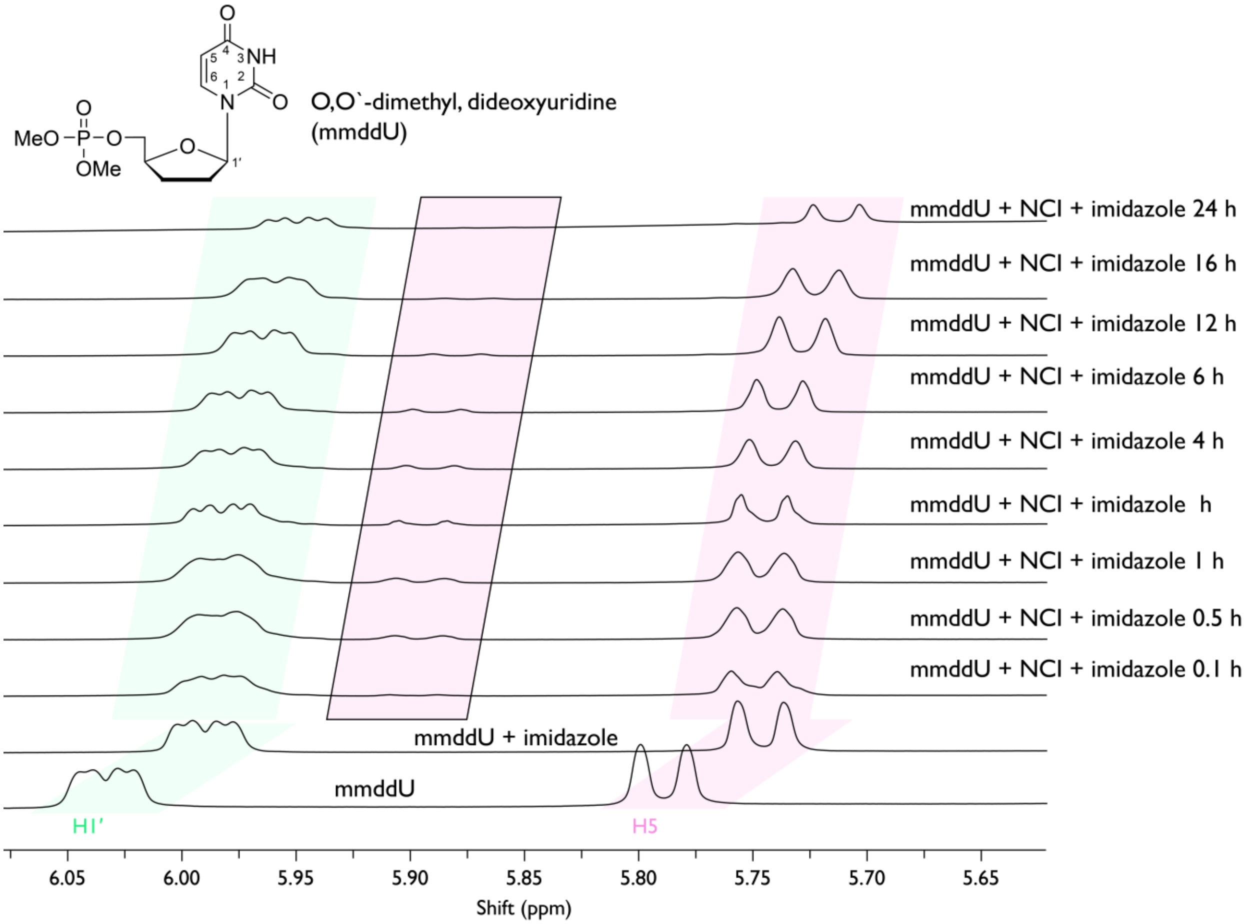


**Figure S34.** $^1H$ NMR spectra of mmddU (25 mM) incubated with 200 mM NCI and 200 mM imidazole in 10 % (v/v) $D_2O$ and 90 % (v/v) $H_2O$ over the course of the reaction. Progressive shifts in the highlighted proton resonances arise from the increase in pH caused by imidazole release due to IDI hydrolysis.

**Figure S35.**

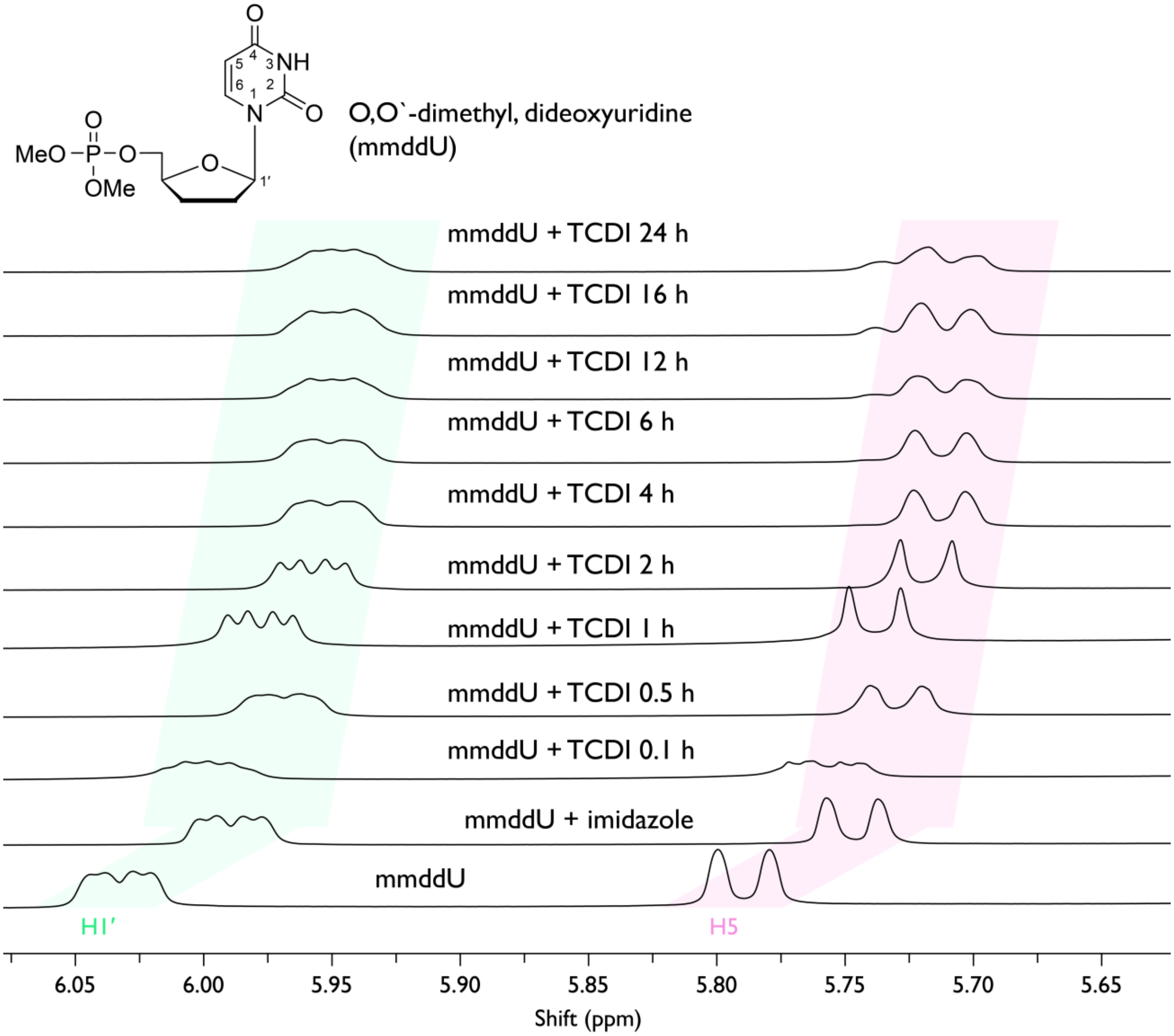


**Figure S35.** [1]H NMR spectra of mmddU (25 mM) incubated with 200 mM TCDI in 10 % (v/v) $D_2O$ and 90 % (v/v) $H_2O$ over the course of the reaction. Progressive shifts in the highlighted proton resonances arise from the increase in pH caused by imidazole release due to IDI hydrolysis.

**Figure S36.**

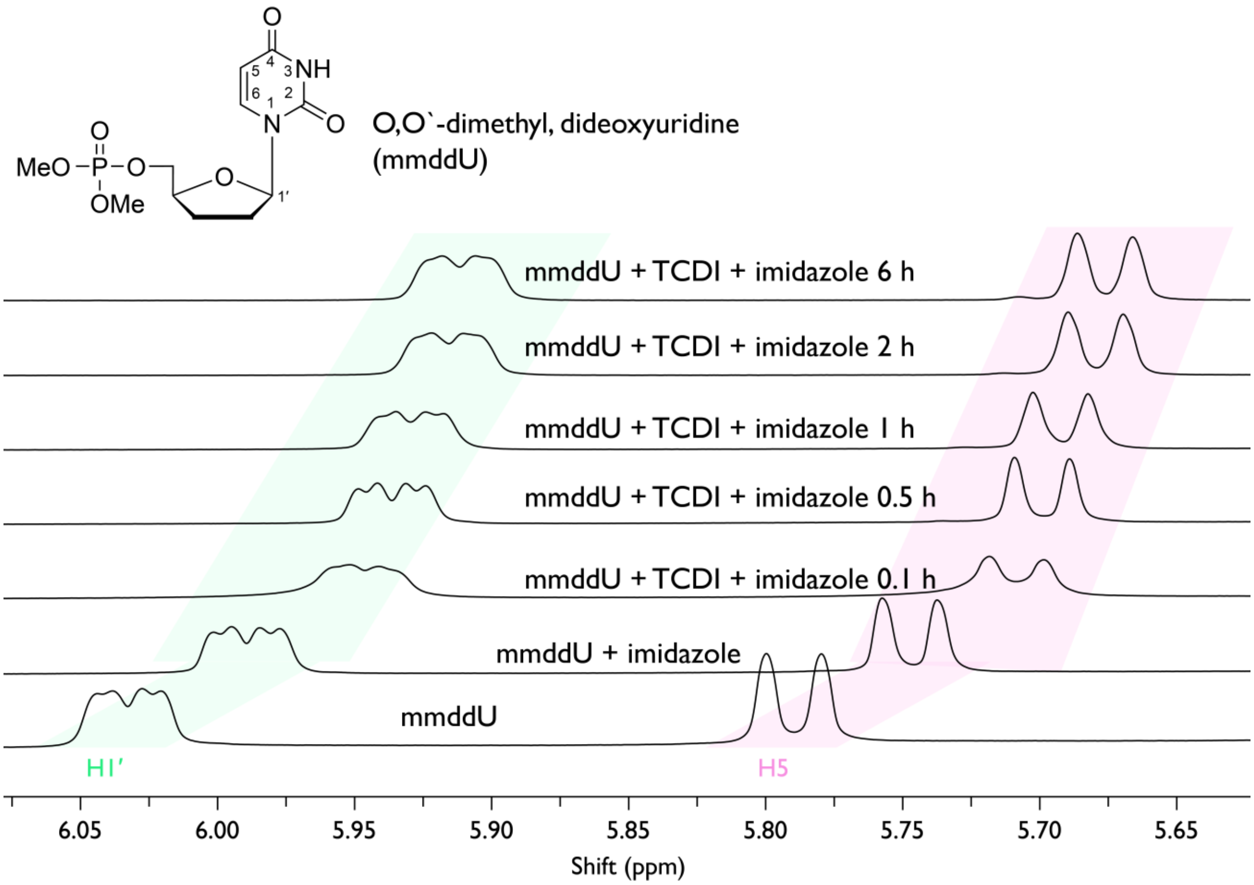


**Figure S36.** $^{1}H$ NMR spectra of mmddU (25 mM) incubated with 200 mM TCDI and 200 mM imidazole in 10 % (v/v) $D_2O$ and 90 % (v/v) $H_2O$ over the course of the reaction. Progressive shifts in the highlighted proton resonances arise from the increase in pH caused by imidazole release due to IDI hydrolysis.

**Figure S37.**

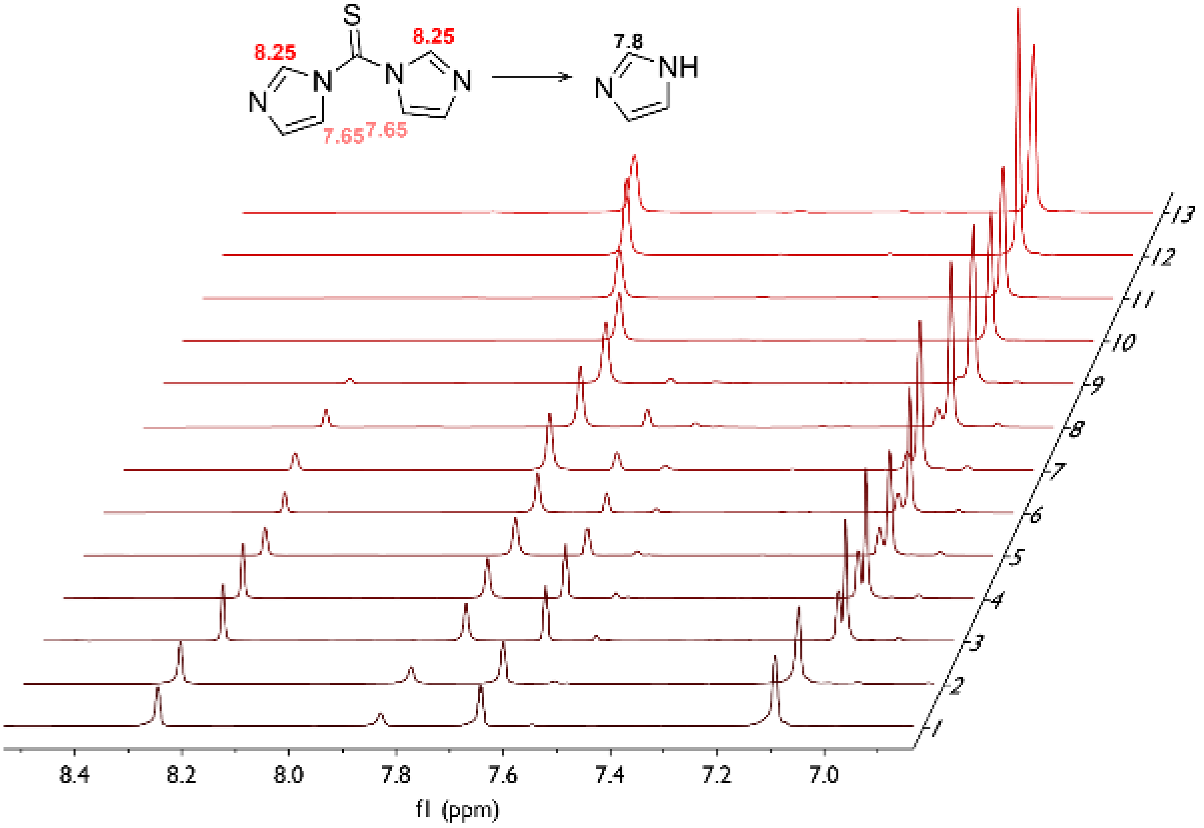


**Figure S37.** [1]H NMR spectra showing hydrolysis of 200 mM TCDI in 10 % (v/v) $D_2O$ and 90 % (v/v) $H_2O$ over time. Spectra labeled 1–13 correspond to reaction times of 0, 5, 10, 15, 20, 25, 30, 35, 120, 180, 240, 300, and 360 minutes.

**Figure S38.**

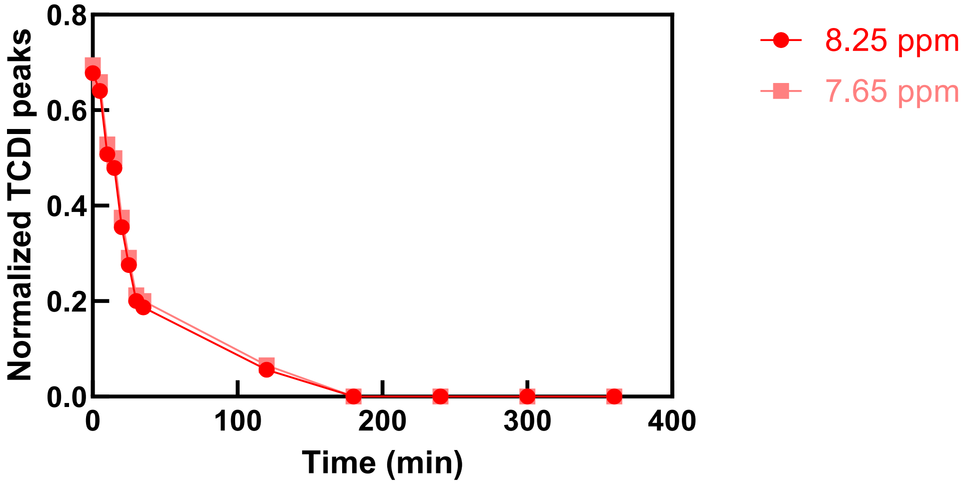


**Figure S38.** Data from [1]H NMR spectra showing hydrolysis of 200 mM TCDI in 10 % (v/v) $D_2O$ and 90 % (v/v) $H_2O$ over time.

**Figure S39.**

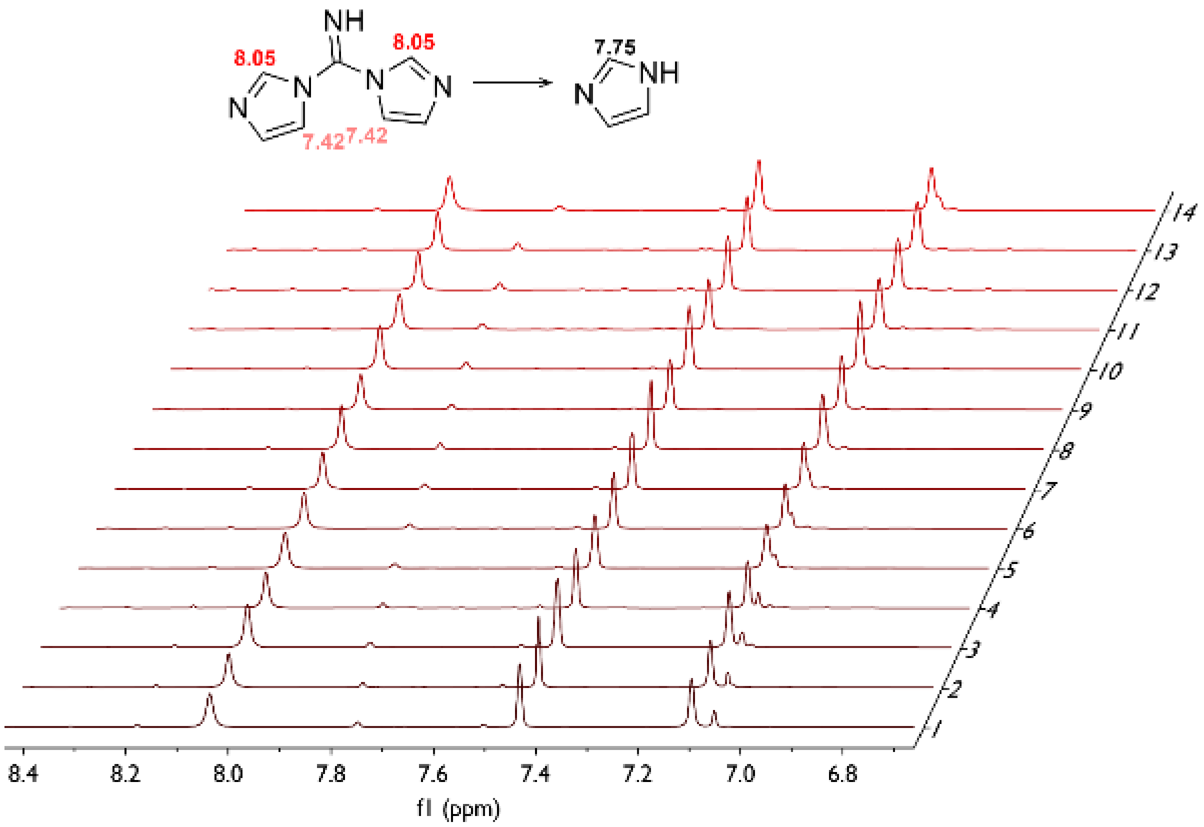


**Figure S39.** $^{1}H$ NMR spectra showing hydrolysis of 200 mM IDI in 10 % (v/v) $D_2O$ and 90 % (v/v) $H_2O$ over time. Spectra labeled 1–13 correspond to reaction times of 0, 5, 10, 15, 20, 25, 30, 35, 120, 180, 240, 300, and 360 minutes.

**Figure S40.**

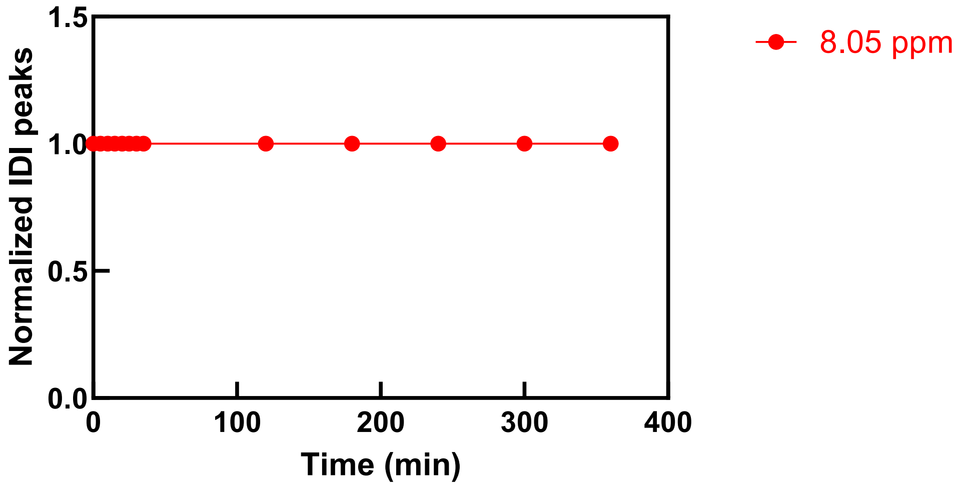


**Figure S40.** Data from [1]H NMR spectra showing hydrolysis of 200 mM IDI in 10 % (v/v) $D_2O$ and 90 % (v/v) $H_2O$ over time.

**Figure S41.**

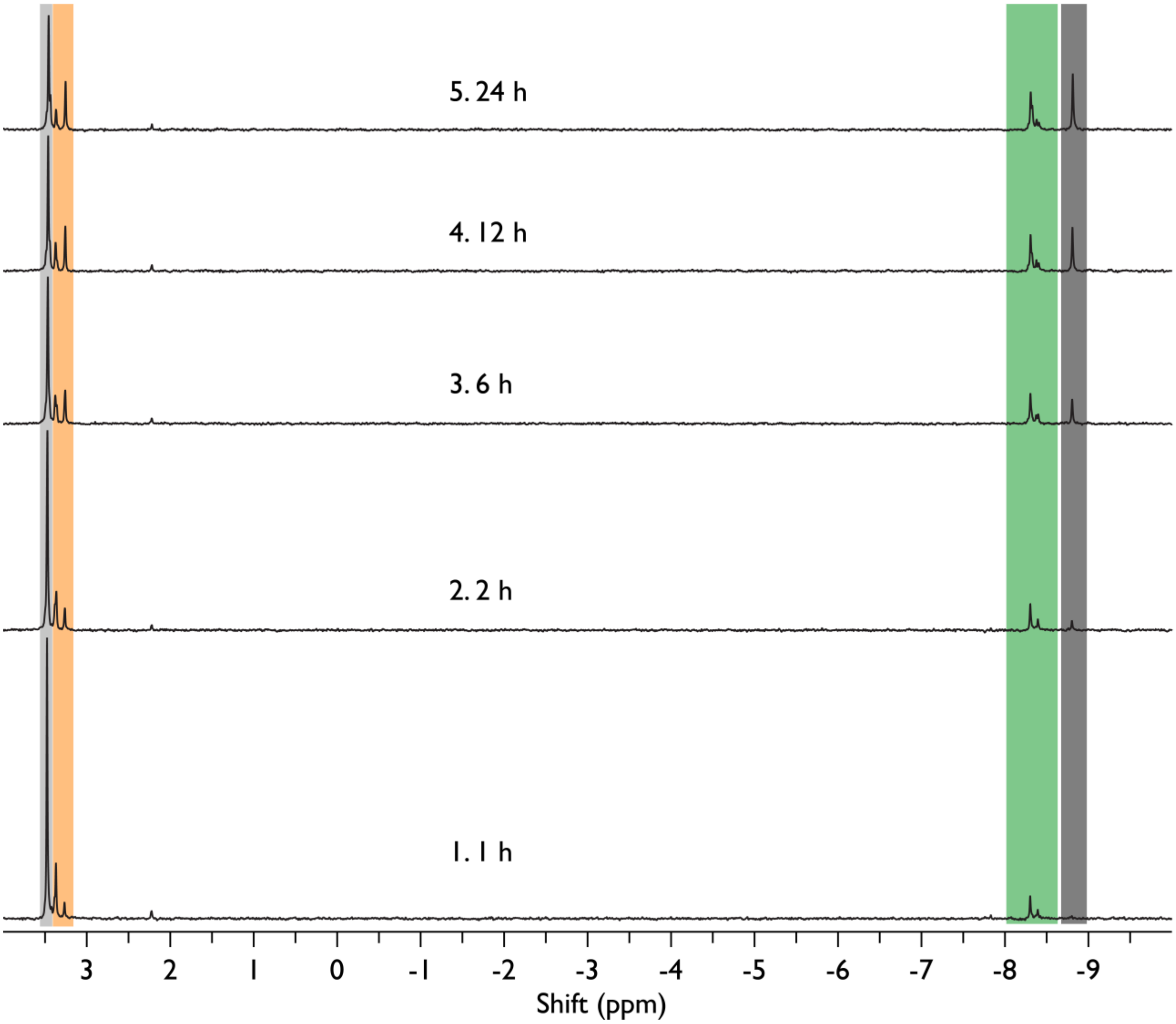


**Figure S41.** Time course of the reaction of UMP at pH 8.0 with IDI monitored by $^{31}P$ NMR. Shown are the consumption of unmodified UMP (blue), formation of the phosphorimidazolide intermediate (UMP-Im, green), and formation of the 2′/3′-acylated product and pyrophosphate (Acyl-UMP, orange + dark grey). Reactions contained 25 mM UMP, 200 mM IDI, and 200 mM HEPES buffer at pH 8.0 in 10% (v/v) $D_2O$. Spectra were collected at 1, 2, 6, 12, and 24 h at room temperature.

**Figure S42.**

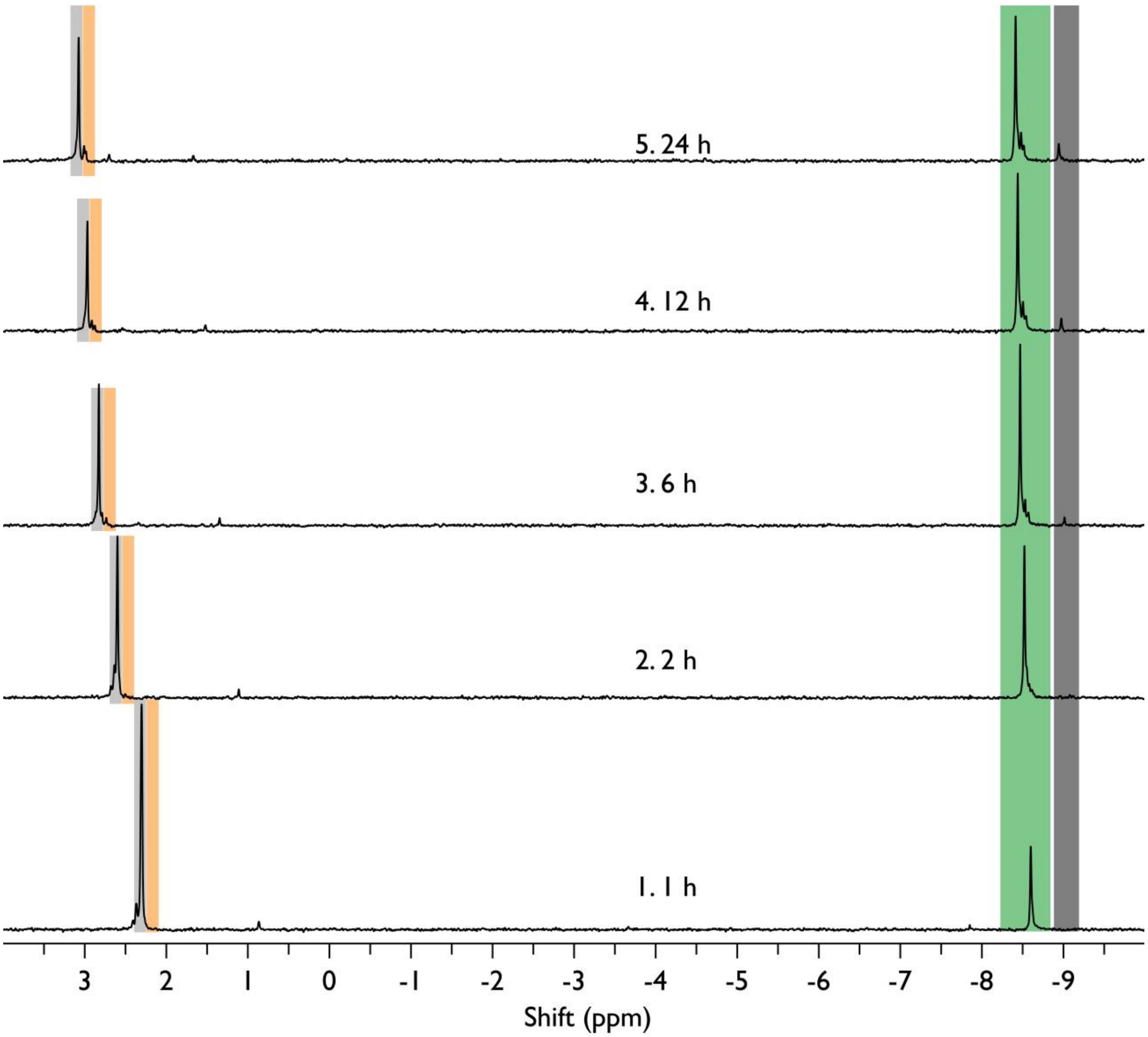


**Figure S42.** Time course of the reaction of UMP at pH 6.0 with IDI monitored by $^{31}$P NMR. Shown are the consumption of unmodified UMP (blue), formation of the phosphorimidazolide intermediate (UMP-Im, green), and formation of the 2′/3′-acylated product and pyrophosphate (Acyl-UMP, orange + dark grey). Reactions contained 25 mM UMP, 200 mM IDI, 200 mM HCl and 200 mM HEPES buffer at pH 8.0 in 10% (v/v) $D_2O$. Spectra were collected at 1, 2, 6, 12, and 24 h at room temperature.

**Figure S43.**

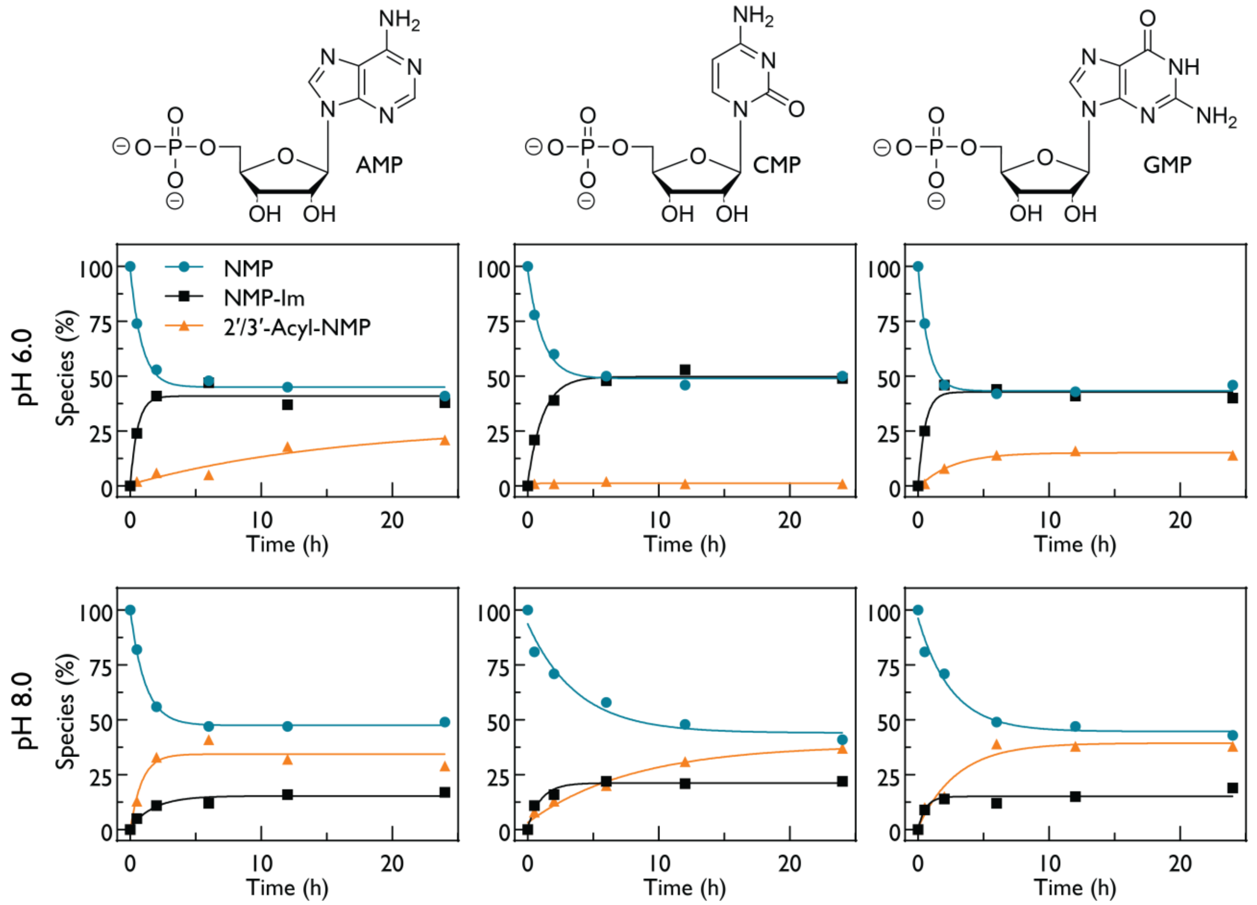


**Figure S43.** Time courses of reactions of nucleotide 5′-monophosphates with IDI, as monitored by $^{31}P$ NMR. Consumption of the unmodified nucleotide (NMP, blue), formation of the phosphorimidazolide intermediate (NMP-Im, black), and formation of the 2′/3′-acylated product (Acyl-NMP, orange). Nucleotides tested were adenosine 5′-monophosphate (AMP), cytidine 5′-monophosphate (CMP), and guanosine 5′-monophosphate (GMP) at 25 mM. Reactions contained 200 mM IDI, 200 mM HEPES buffer at pH 6.0 or pH 8.0 (as indicated), and 200 mM HCl for the pH 6.0 condition, in 10% (v/v) $D_2O$, incubated for 24 h at room temperature. Kinetic traces were fit to a single step model, and the corresponding rates for NMP, NMP-Im, and Acyl-NMP are reported in **Table S2**.

**Figure S44.**

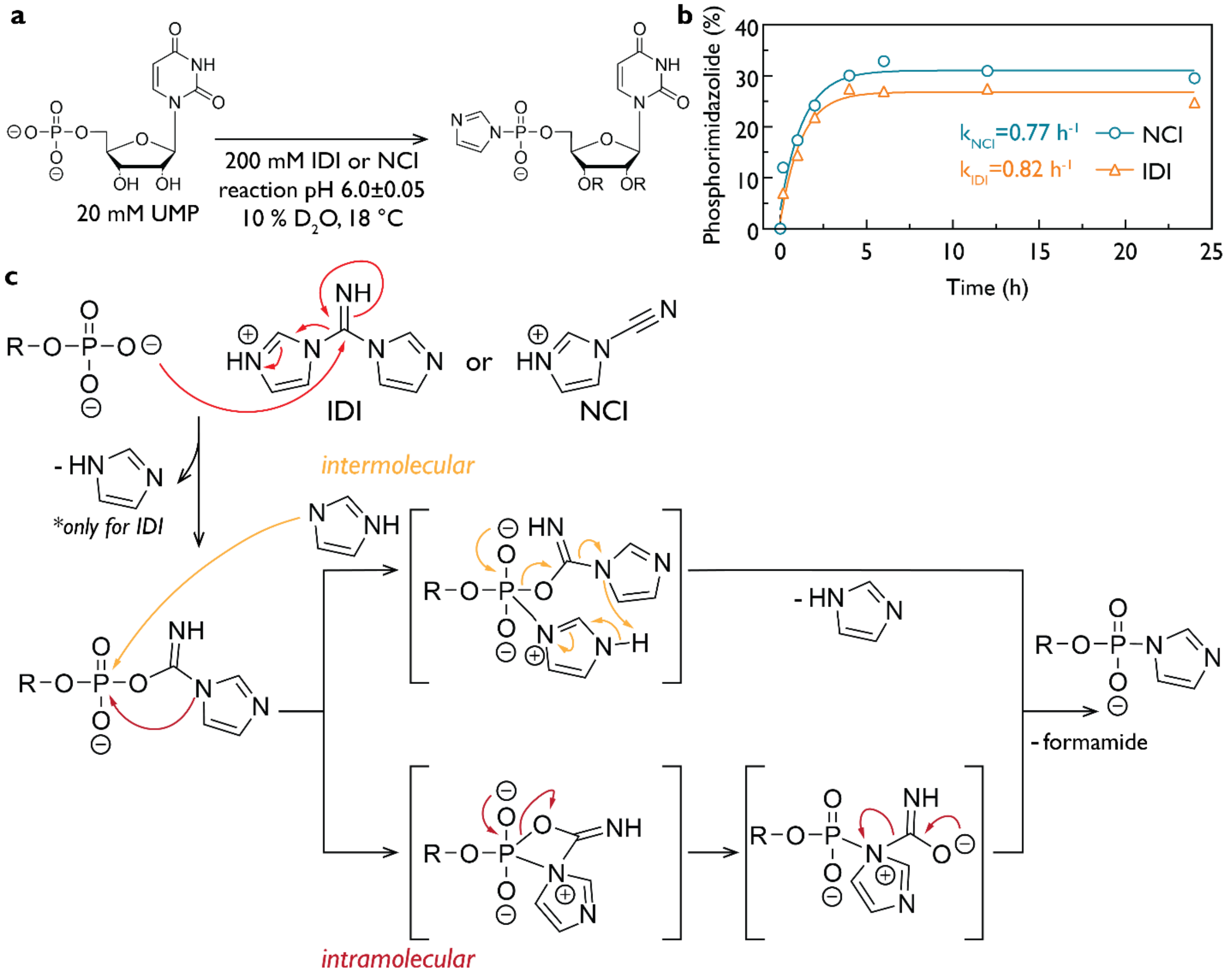


**Figure S44. (a)** Experimental setup for [1]H NMR kinetic analysis of phosphate activation of UMP (20 mM) by 200 mM IDI or 200 mM NCI at pH 6.0. The pH was maintained between 6.0 and 6.5 by periodic addition of concentrated HCl. Samples contained 10 % (v/v) $D_2O$ for NMR analysis. **(b)** $^{31}$P NMR time courses of UMP activation under the same conditions, showing formation of the phosphorimidazolide intermediate. Data were fit to a single-step kinetic model, yielding observed rates of 0.82 $h^{-1}$ for IDI and 0.77 $h^{-1}$ for NCI. **(c)** Proposed mechanistic pathways for IDI- and NCI-mediated activation of nucleotide 5′-monophosphates *via* intermolecular or intramolecular pathways.

**Figure S45.**

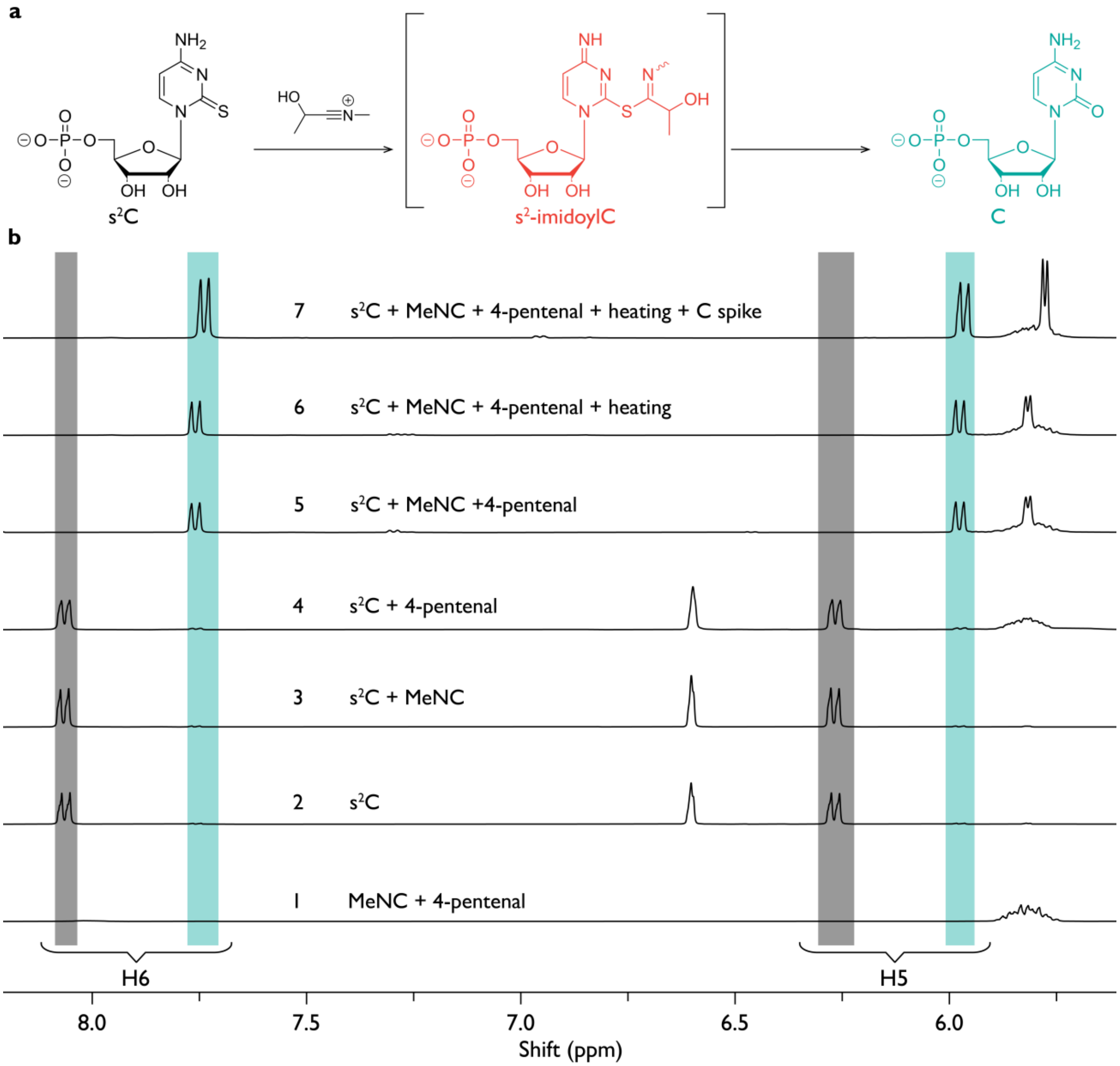


**Figure S45**. **(a)** Proposed reaction scheme for s²C exposed to isonitrile–aldehyde activation conditions, illustrating nucleophilic interception of the reactive intermediate by the C2-linked sulfur to form an S-imidoyl-s²C intermediate. **(b)** $^{1}$H NMR spectra of s²C (25 mM) in 200 mM HEPES pH 8.0 containing 10 % $D_2O$ under the following conditions: (1) 200 mM MeNC and 200 mM 4-pentenal; (2) s²C alone; (3) s²C with 200 mM MeNC; (4) s²C with 200 mM 4-pentenal; (5) s²C with 200 mM MeNC and 200 mM 4-pentenal; (6) sample from condition 5 heated at 95 °C for 15 min; and (7) sample from condition 6 spiked with 25 mM cytidine (C). Resonances corresponding to s²C H5 and H6 are shown in grey and C resonances in turquoise.

**Figure S46.**

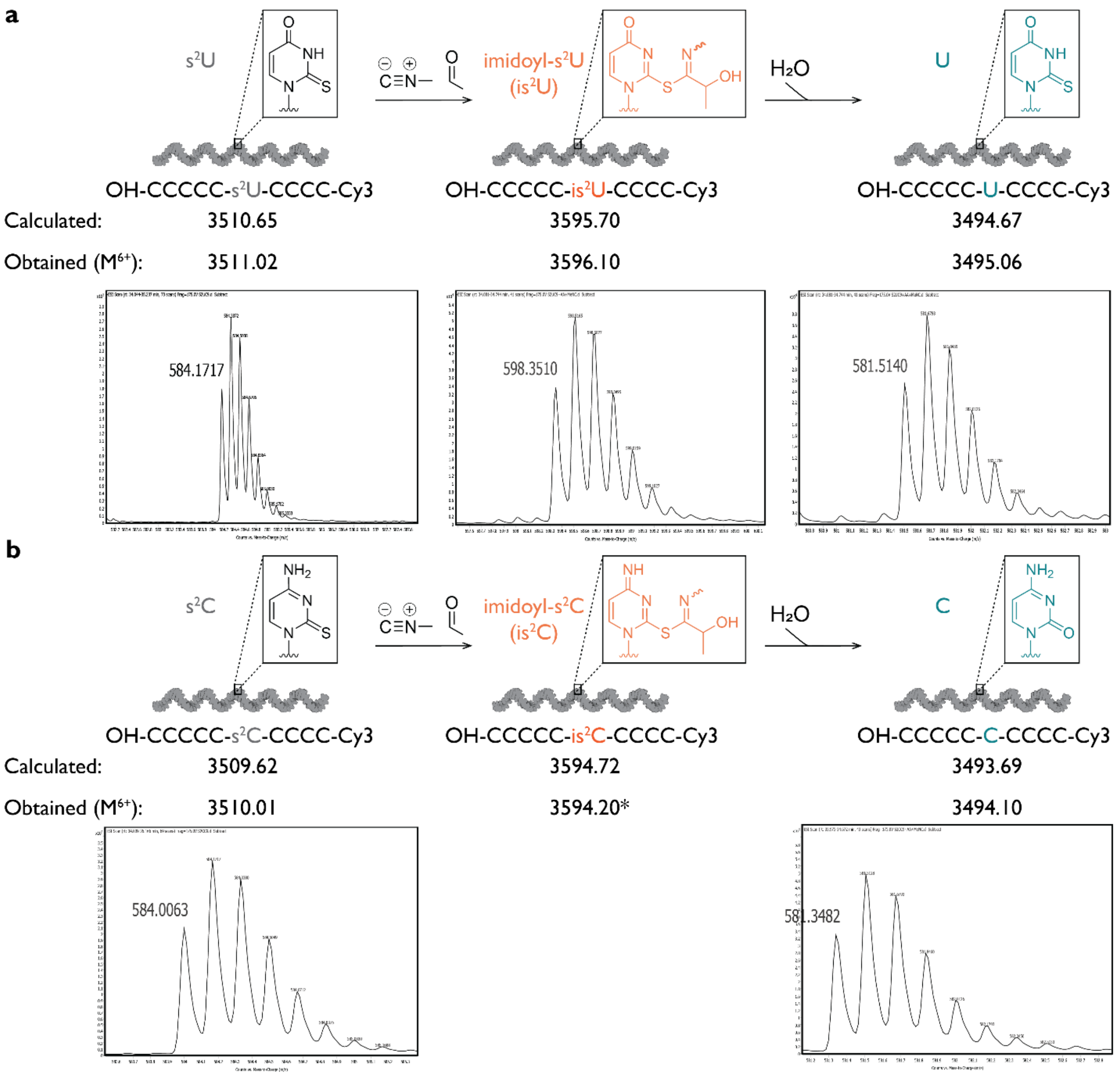


**Figure S46**. **(a)** HPLC-qTOF mass spectrometry analysis of a 10-mer 3′-Cy3 RNA oligonucleotide with sequence $C_5$-s²U-$C_4$ treated with 200 mM MeNC and 200 mM acetaldehyde for 24 h, followed by desalting and separation on a C18 column prior to detection. Signals corresponding to the s²U-containing RNA, an imidoyl-modified species, and a desulfurized U-containing RNA are indicated. **(b)** Corresponding HPLC-qTOF analysis of a 10-mer 3′-Cy3 RNA oligonucleotide with sequence $C_5$-s²C-$C_4$ under identical conditions. Signals corresponding to the s²C-containing RNA and a desulfurized C-containing RNA are observed. The $M^{6-}$ charge state is shown, with m/z values derived from multiple charge states. The imidoyl-s²C-modified species was within the detection limits but exhibited insufficient signal-to-noise for reliable assignment.

**Figure S47.**

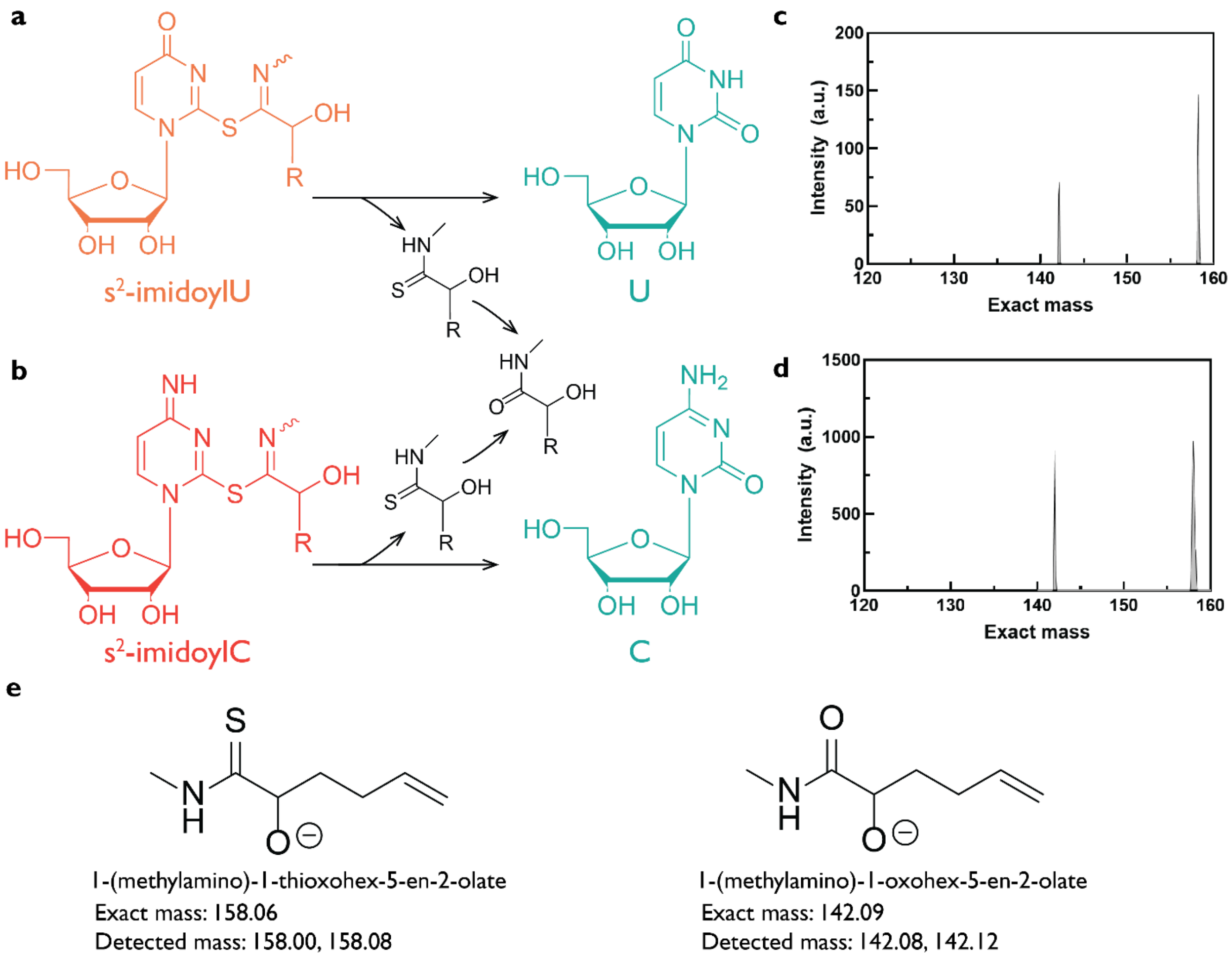


**Figure S47**. **(a)** Proposed hydrolytic transformations of an S-imidoyl intermediate generated under isonitrile–aldehyde activation conditions, leading to formation of thioamide and amide products. **(b)** Proposed analogous transformations for sulfur-modified cytidine derivatives. **(c)** Mass spectrometry of $s^2U$ reaction mixtures showing detection of a thioamide-containing byproduct. **(d)** Mass spectrometry of $s^2C$ reaction mixtures showing detection of the corresponding thioamide byproduct. **(e)** Structures of the proposed thioamide and amide products with calculated exact masses and experimentally detected exact masses.

**Figure S48.**

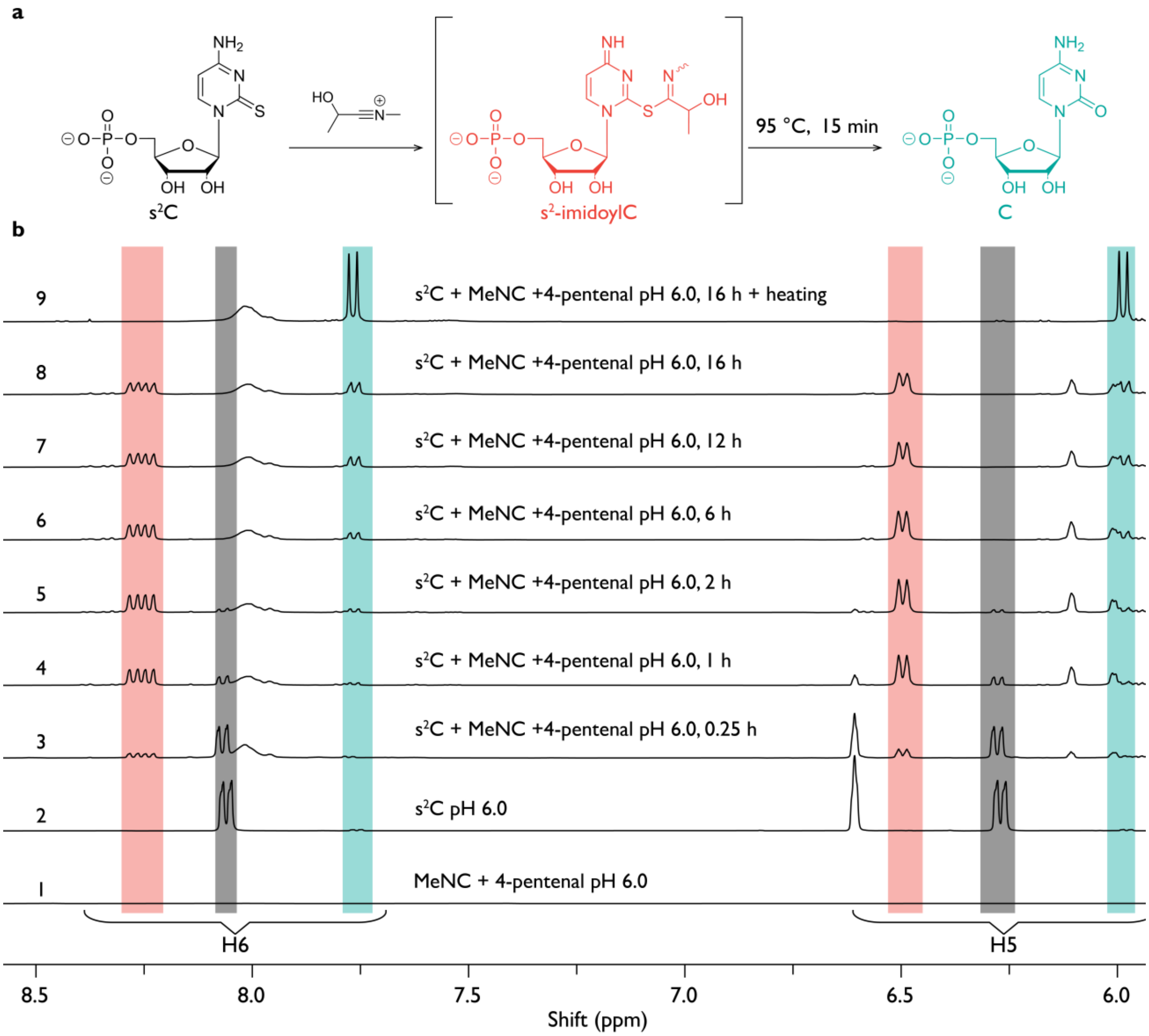


**Figure S48**. **(a)** Proposed reaction scheme for s²C under isonitrile–aldehyde activation conditions, illustrating formation of an S-imidoyl-s²C intermediate. **(b)** $^1$H NMR spectra of s²C (25 mM) in 200 mM HEPES **pH 6.0** containing 10 % $D_2O$ under the following conditions: (1) 200 mM MeNC and 200 mM 4-pentenal; (2) s²C alone; (3–9) s²C incubated with 200 mM MeNC and 200 mM 4-pentenal over time. Resonances corresponding to s²C H5 and H6 are shown in grey, cytidine resonances in turquoise, and intermediate resonances in orange.

**Figure S49.**

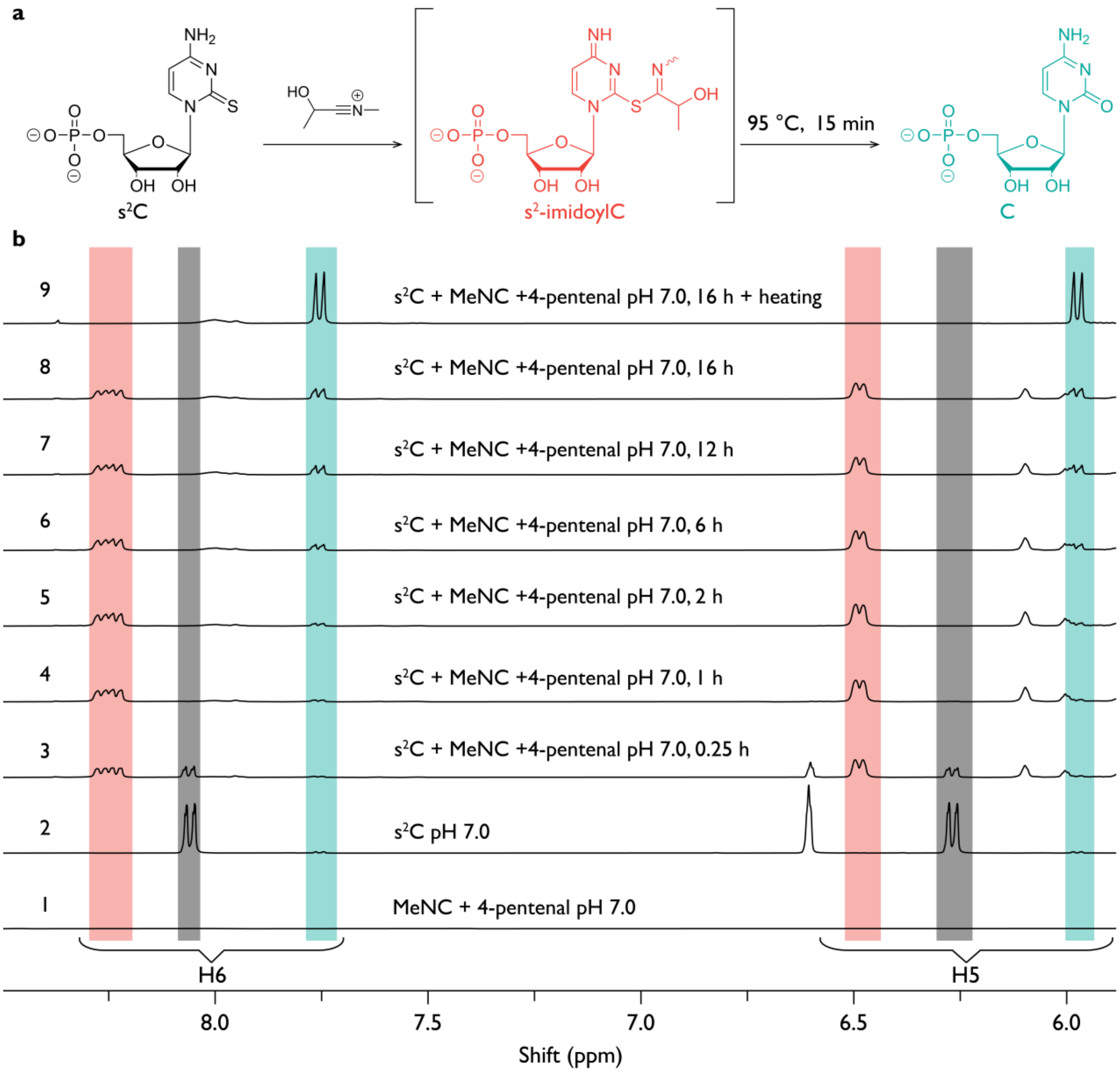


**Figure S49**. **(a)** Proposed reaction scheme for s²C under isonitrile–aldehyde activation conditions, illustrating formation of sulfur-modified intermediates. **(b)** $^{1}$H NMR spectra of s²C (25 mM) in 200 mM HEPES **pH 7.0** containing 10 % $D_2O$ under the following conditions: (1) 200 mM MeNC and 200 mM 4-pentenal; (2) s²C alone; (3–9) s²C incubated with 200 mM MeNC and 200 mM 4-pentenal over time. Resonances corresponding to s²C H5 and H6 are shown in grey, cytidine resonances in turquoise, and intermediate resonances in orange.

**Figure S50.**

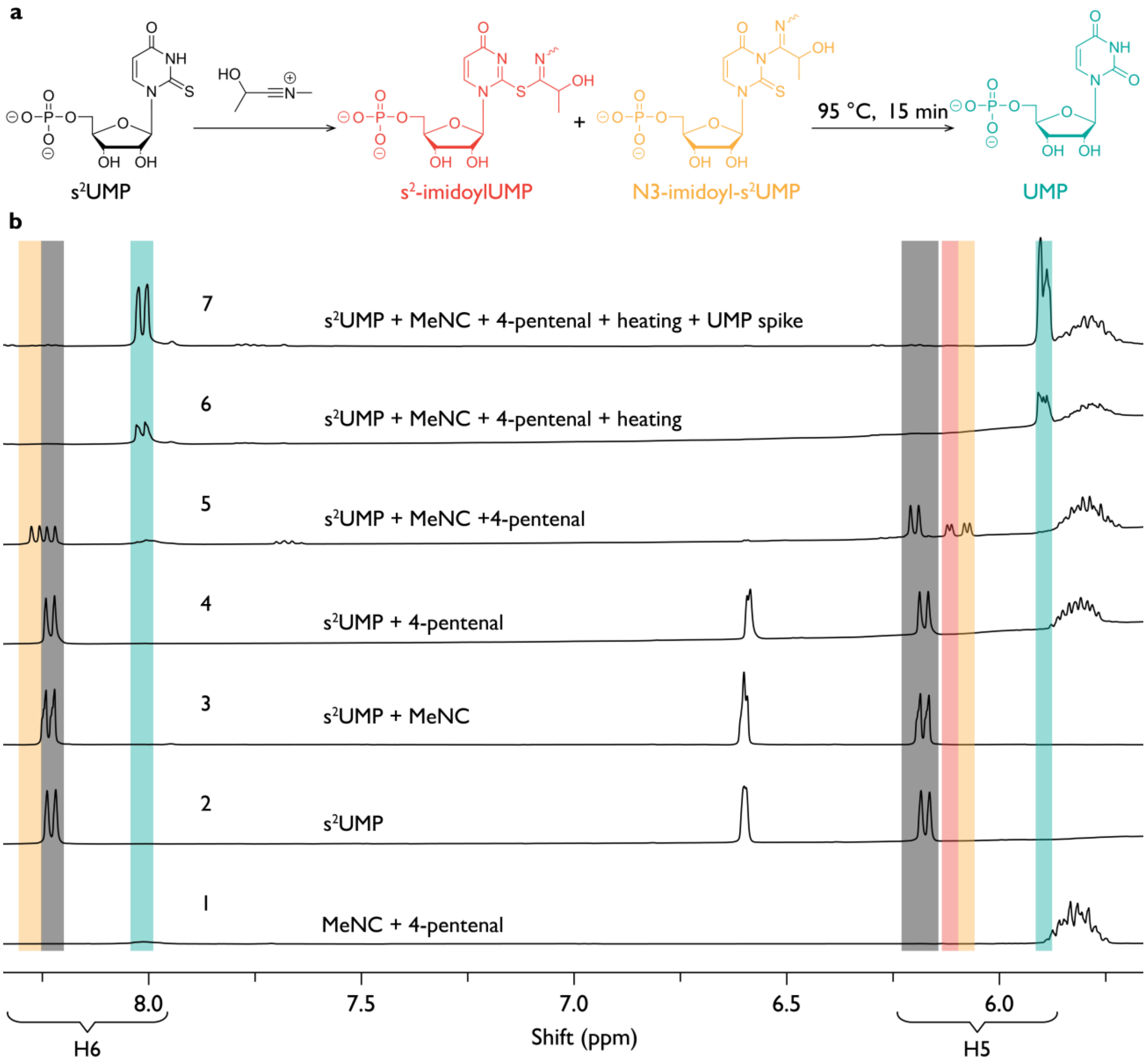


**Figure S50**. **(a)** Proposed reaction scheme for s²UMP exposed to isonitrile–aldehyde activation conditions, illustrating formation of sulfur- and nitrogen-modified intermediates. **(b)** $^{1}$H NMR spectra of s²UMP (25 mM) in 200 mM HEPES **pH 8.0** containing 10 % $D_2O$ under the following conditions: (1) 200 mM MeNC and 200 mM 4-pentenal; (2) s²UMP alone; (3) s²UMP with 200 mM MeNC; (4) s²UMP with 200 mM 4-pentenal; (5) s²UMP with 200 mM MeNC and 200 mM 4-pentenal; (6) sample from condition 5 heated at 95 °C for 15 min; and (7) sample from condition 6 spiked with 25 mM UMP. Resonances corresponding to s²UMP H5 and H6 are shown in grey, UMP in turquoise, and intermediate resonances in shades of orange.

**Figure S51.**

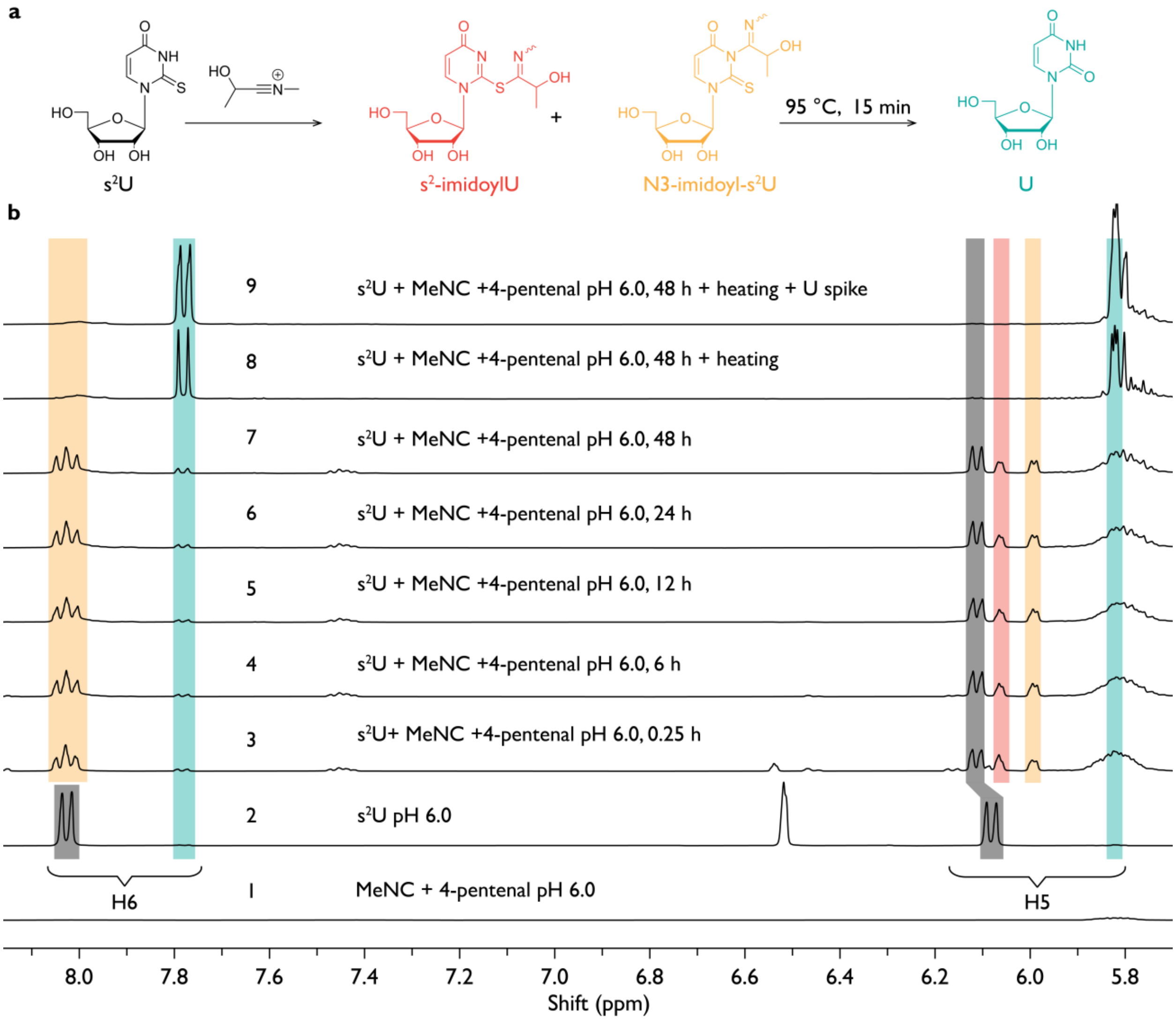


**Figure S51**. **(a)** Proposed reaction scheme for s²U exposed to isonitrile–aldehyde activation conditions, illustrating formation of imidoyl-modified intermediates. **(b)** $^1$H NMR spectra of s²U (25 mM) in 200 mM HEPES **pH 6.0** containing 10 % $D_2O$ under the following conditions: (1) 200 mM MeNC and 200 mM 4-pentenal; (2) s²U alone; (3–9) s²U incubated with 200 mM MeNC and 200 mM 4-pentenal over time; (9) sample spiked with 25 mM uridine. Resonances corresponding to s²U H5 and H6 are shown in grey, uridine resonances in turquoise, and intermediate resonances in orange.

**Figure S52.**

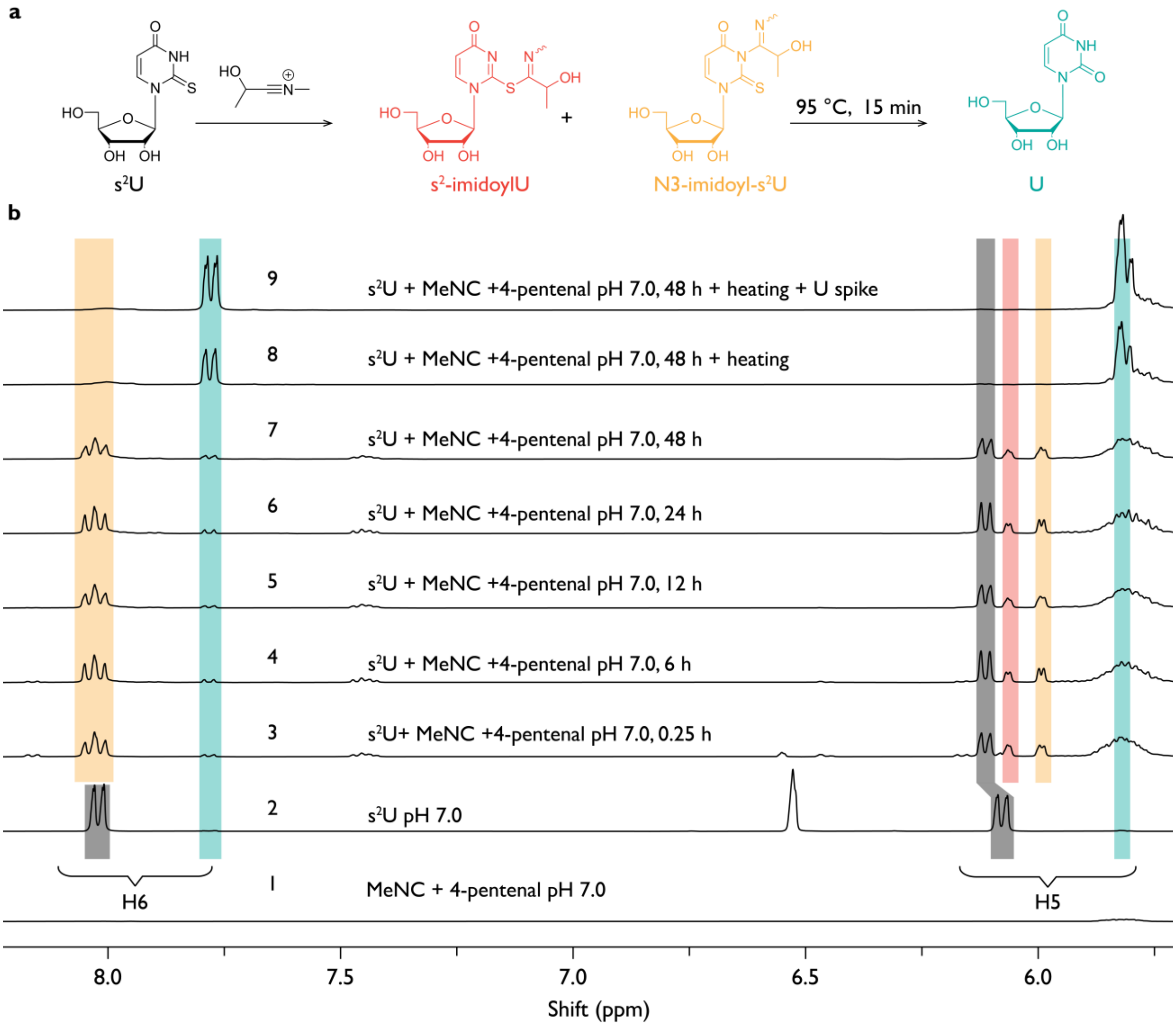


**Figure S52**. **(a)** Proposed reaction scheme for s²U under isonitrile–aldehyde activation conditions. **(b)** $^{1}$H NMR spectra of s²U (25 mM) in 200 mM HEPES **pH 7.0** containing 10 % $D_2O$ under the following conditions: (1) 200 mM MeNC and 200 mM 4-pentenal; (2) s²U alone; (3–9) s²U incubated with 200 mM MeNC and 200 mM 4-pentenal over time; (9) sample spiked with 25 mM uridine. Resonances corresponding to s²U H5 and H6 are shown in grey, uridine resonances in turquoise, and intermediate resonances in orange.

**Figure S53.**

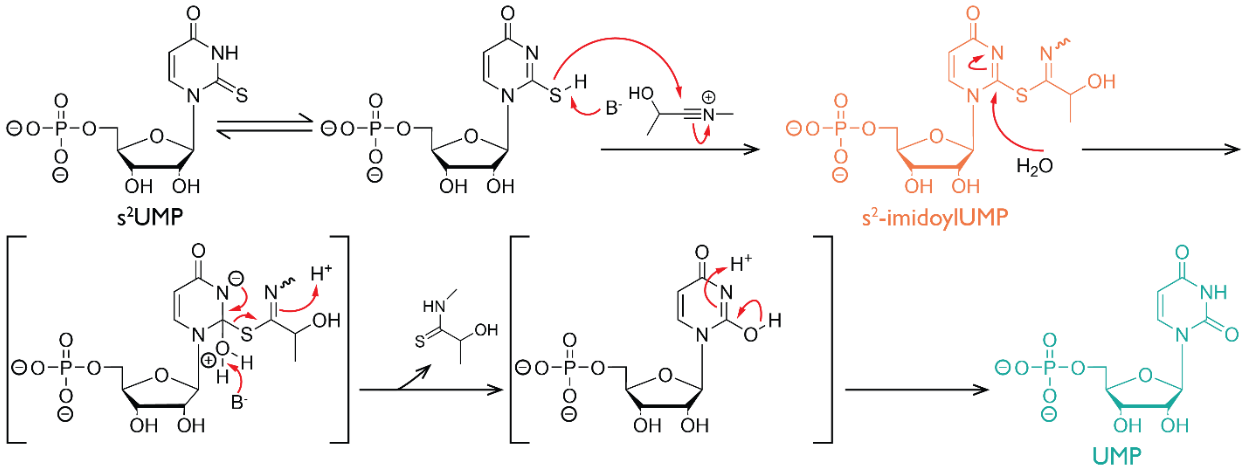


**Figure S53**. Proposed reaction pathway for s²UMP under isonitrile–aldehyde activation conditions, illustrating formation of sulfur-modified intermediates and their subsequent transformation upon heating.

**Figure S54.**

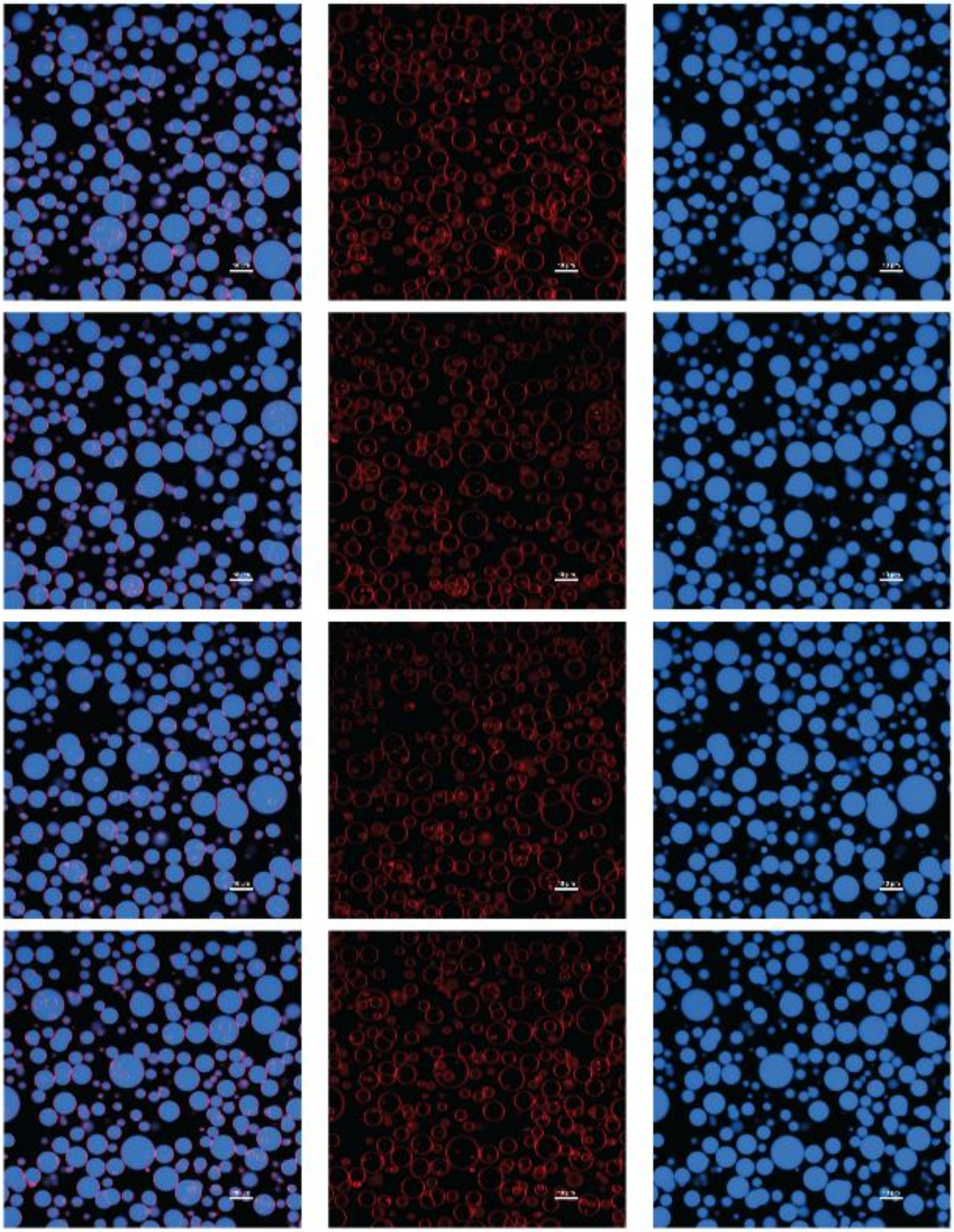

**Figure S54.** Confocal fluorescence images of oleic acid vesicles in the absence of phosphate activation agent(s). Three columns show the overlay of red (Rhodamine B, lipid bilayer) and blue (Cy5-labeled 12-nt RNA) channels, as well as the individual fluorescence channels. Scale bar, 20 μm.

**Figure S55.**

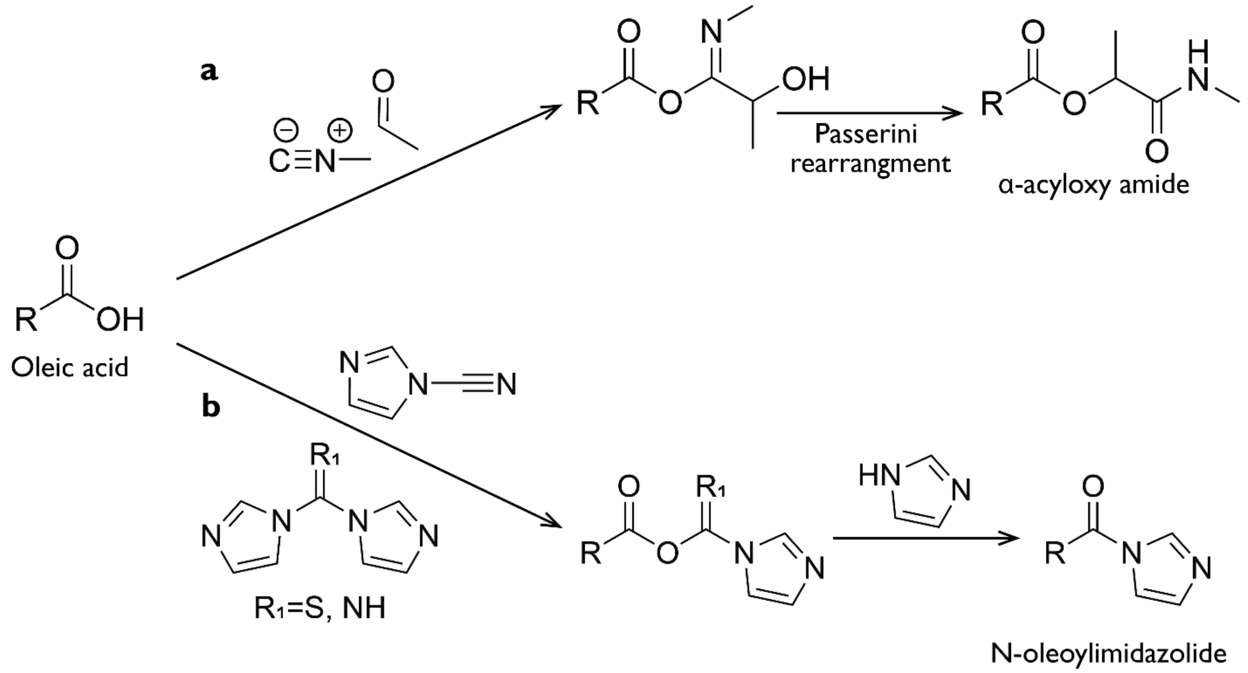


**Figure S55**. **(a)** Proposed products of the reaction of oleic acid vesicles with isonitrile–aldehyde activation chemistry[1]. Oleic acid is suggested to participate in a Passerini reaction with methyl isonitrile and acetaldehyde to form an α-acyloxyamide product that is poorly amphiphilic and therefore tends to phase separate from the oleic acid bilayer. Accumulation of this product is proposed to promote formation of oil-like domains associated with the vesicle membrane, resulting in oil-embedded vesicles observed under isonitrile–aldehyde activation conditions. **(b)** Proposed reaction of oleate with acylimidazoles[2], in which nucleophilic attack by oleate generates a reactive intermediate that can be intercepted by imidazole to form N-oleoylimidazolide.

**Figure S56.**

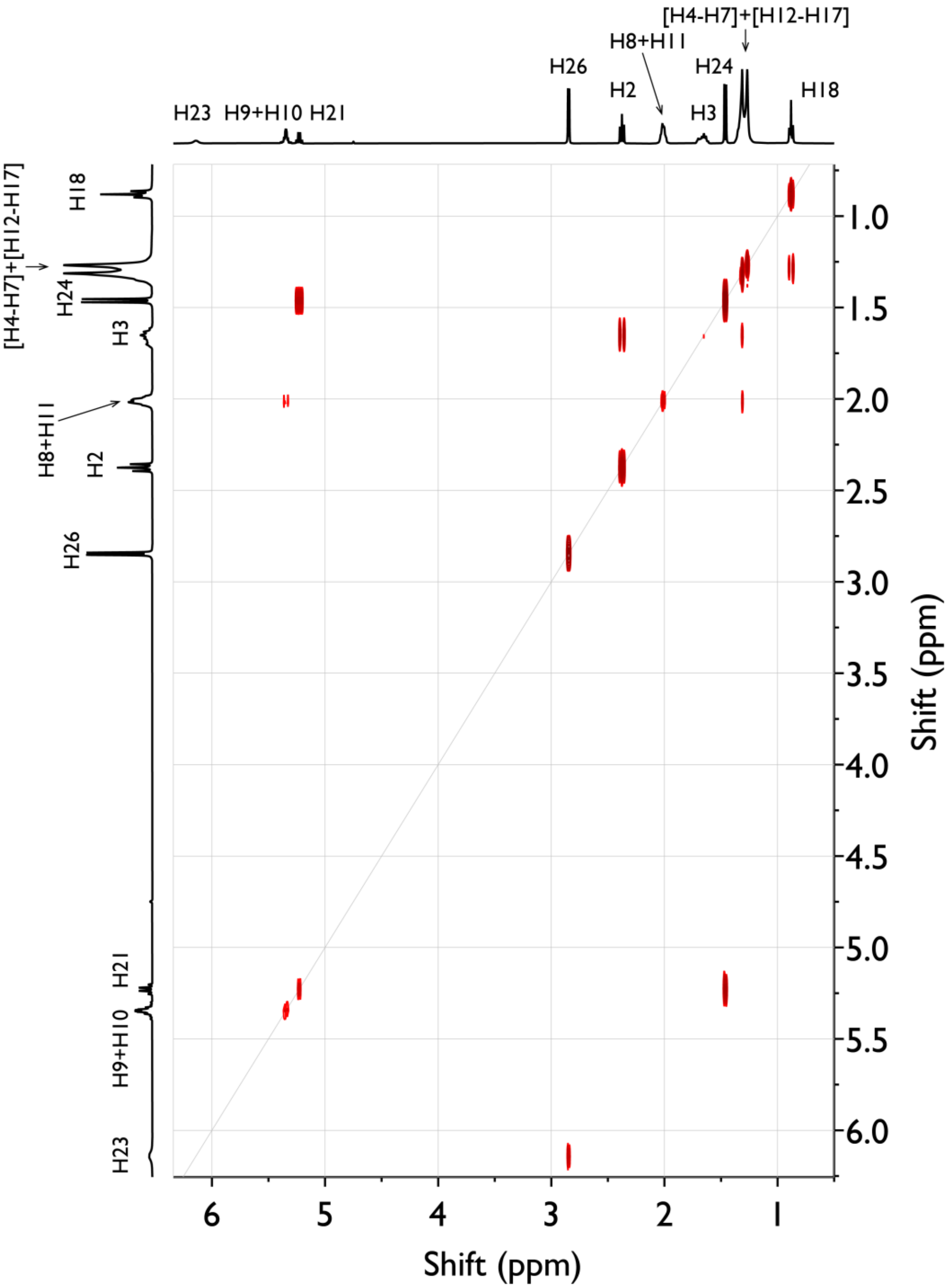


**Figure S56.** $^1H$–$^1H$ COSY NMR spectrum of 1-(methylamino)-1-oxopropan-2-yl oleate recorded in $CDCl_3$ at 400 MHz, showing through-bond proton–proton correlations of the α-oxopropanamide moiety, consistent with the assigned structure of the Passerini-derived oleic acid product.

**Figure S57.**

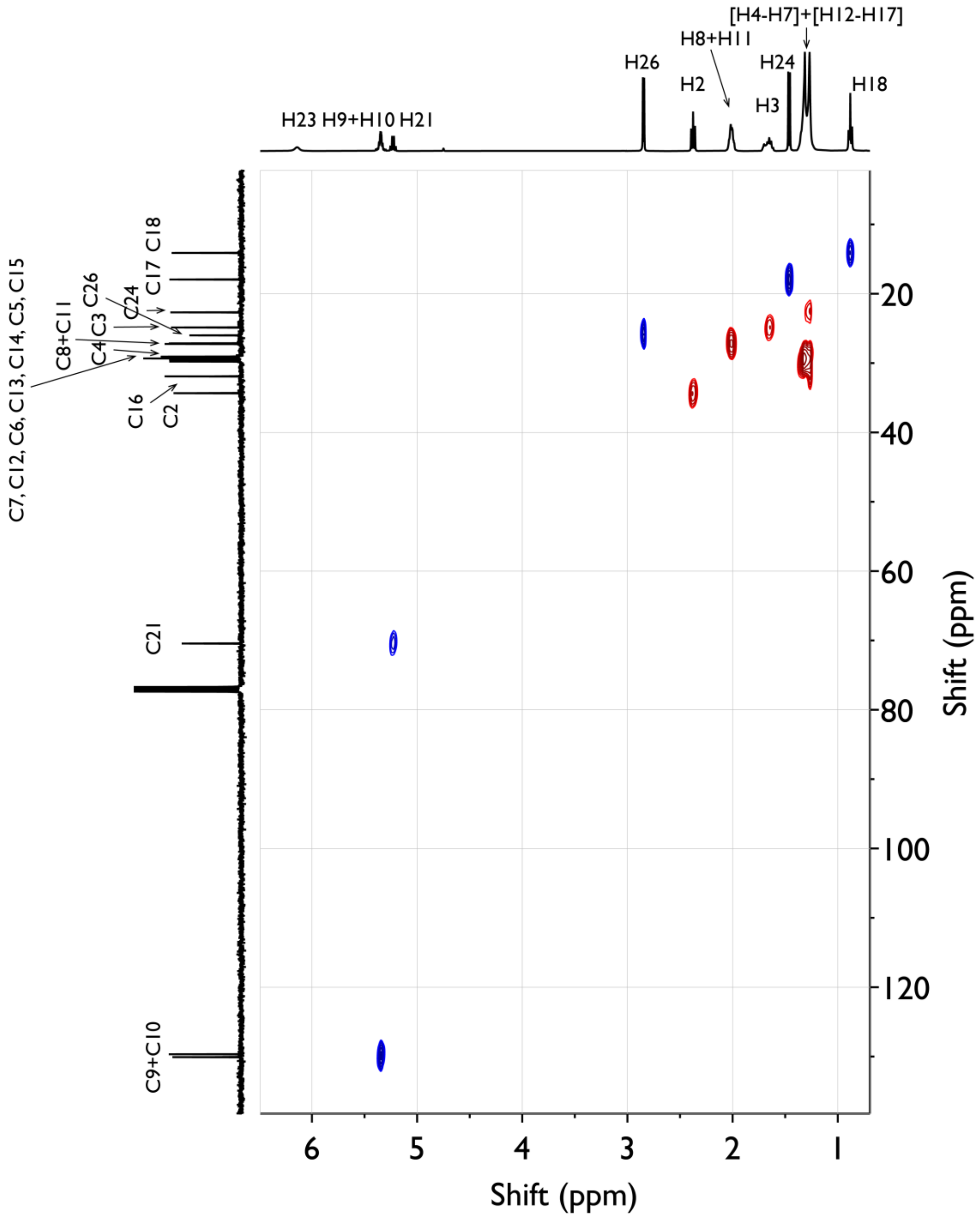


**Figure S57.** $^{13}C$–$^{1}H$ HSQC NMR spectrum of 1-(methylamino)-1-oxopropan-2-yl oleate recorded in $CDCl_3$ at 400 MHz, showing one-bond $^{13}C$–$^{1}H$ correlations that confirm the assignment of protonated carbon centers within the oleate chain and the α-oxopropanamide moiety.

**Figure S58.**

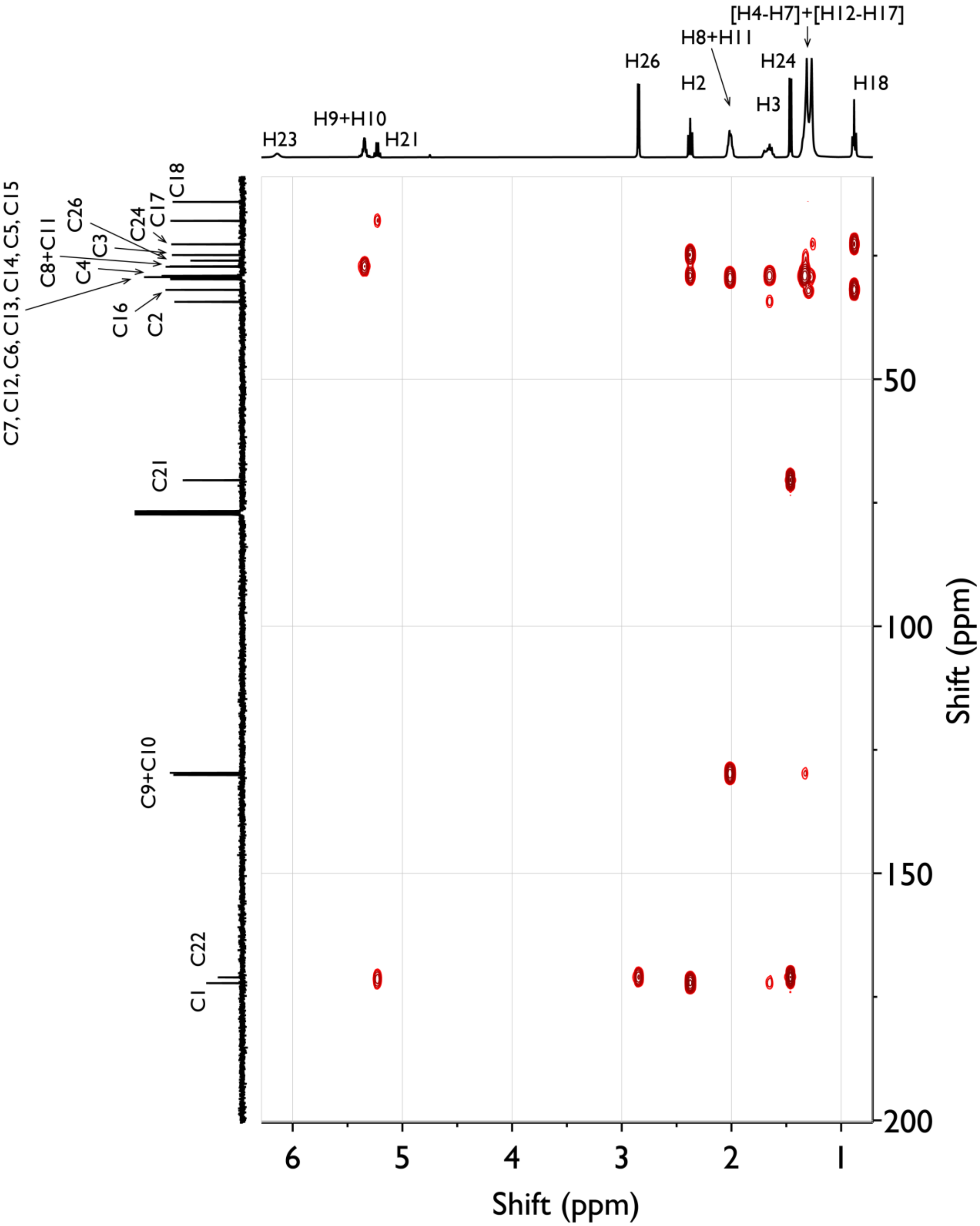


**Figure S58.** $^{1}H$–$^{13}C$ HMBC NMR spectrum of 1-(methylamino)-1-oxopropan-2-yl oleate recorded in $CDCl_3$ at 400 MHz, showing long-range two- and three-bond $^{1}H$–$^{13}C$ correlations.

**Figure S59.**

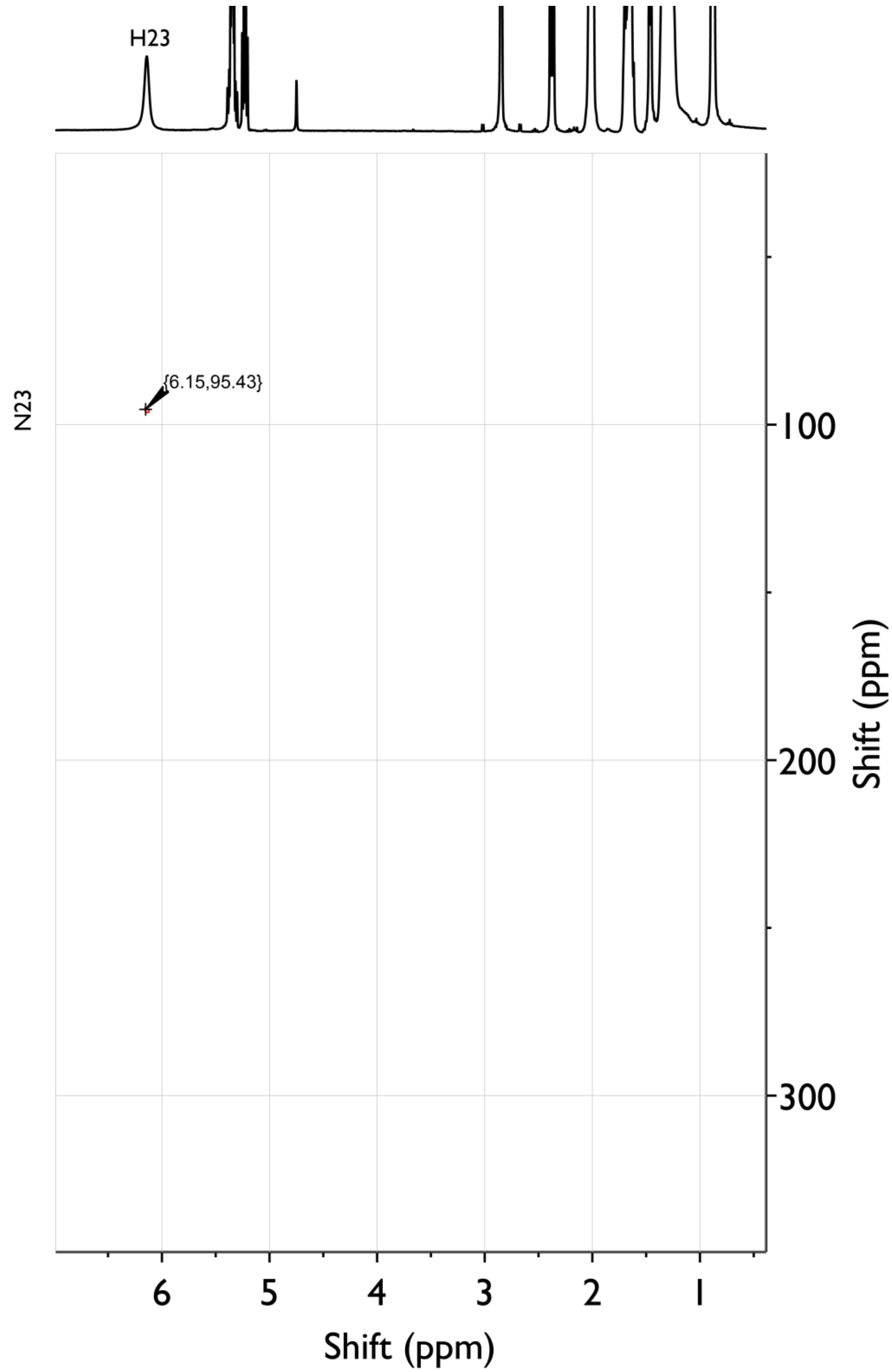


**Figure S59.** $^{15}N$–$^{1}H$ HSQC NMR spectrum of 1-(methylamino)-1-oxopropan-2-yl oleate recorded in $CDCl_3$ at 400 MHz, showing one-bond $^{15}N$–$^{1}H$ correlations that confirm the presence and connectivity of the amide nitrogen within the oxopropanamide moiety.

**Figure S60.**

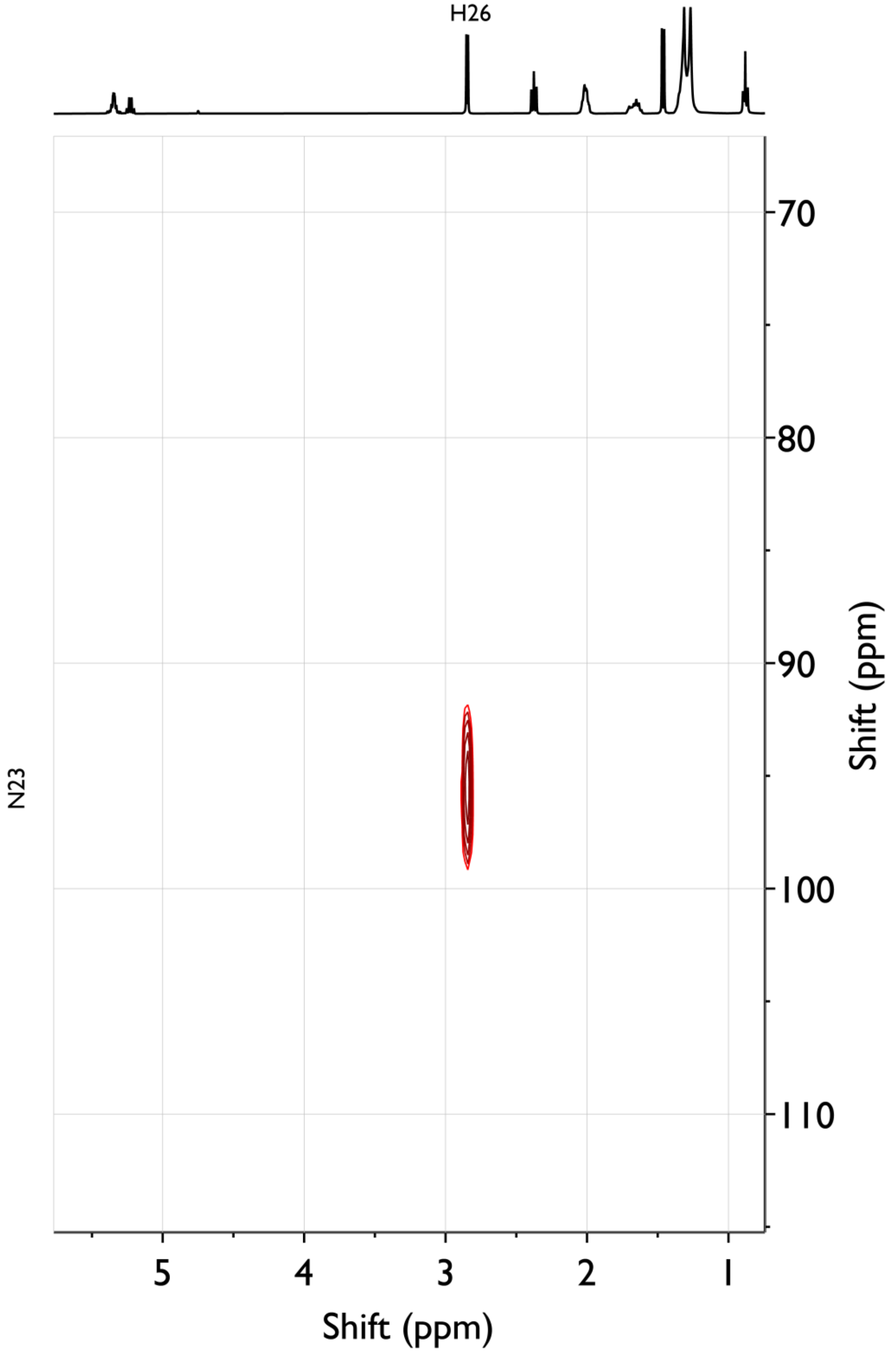


**Figure S60.** $^{15}N$–$^{1}H$ HMBC NMR spectrum of 1-(methylamino)-1-oxopropan-2-yl oleate recorded in $CDCl_3$ at 400 MHz, showing long-range $^{15}N$–$^{1}H$ correlations that link the amide nitrogen to adjacent proton.

**Figure S61.**

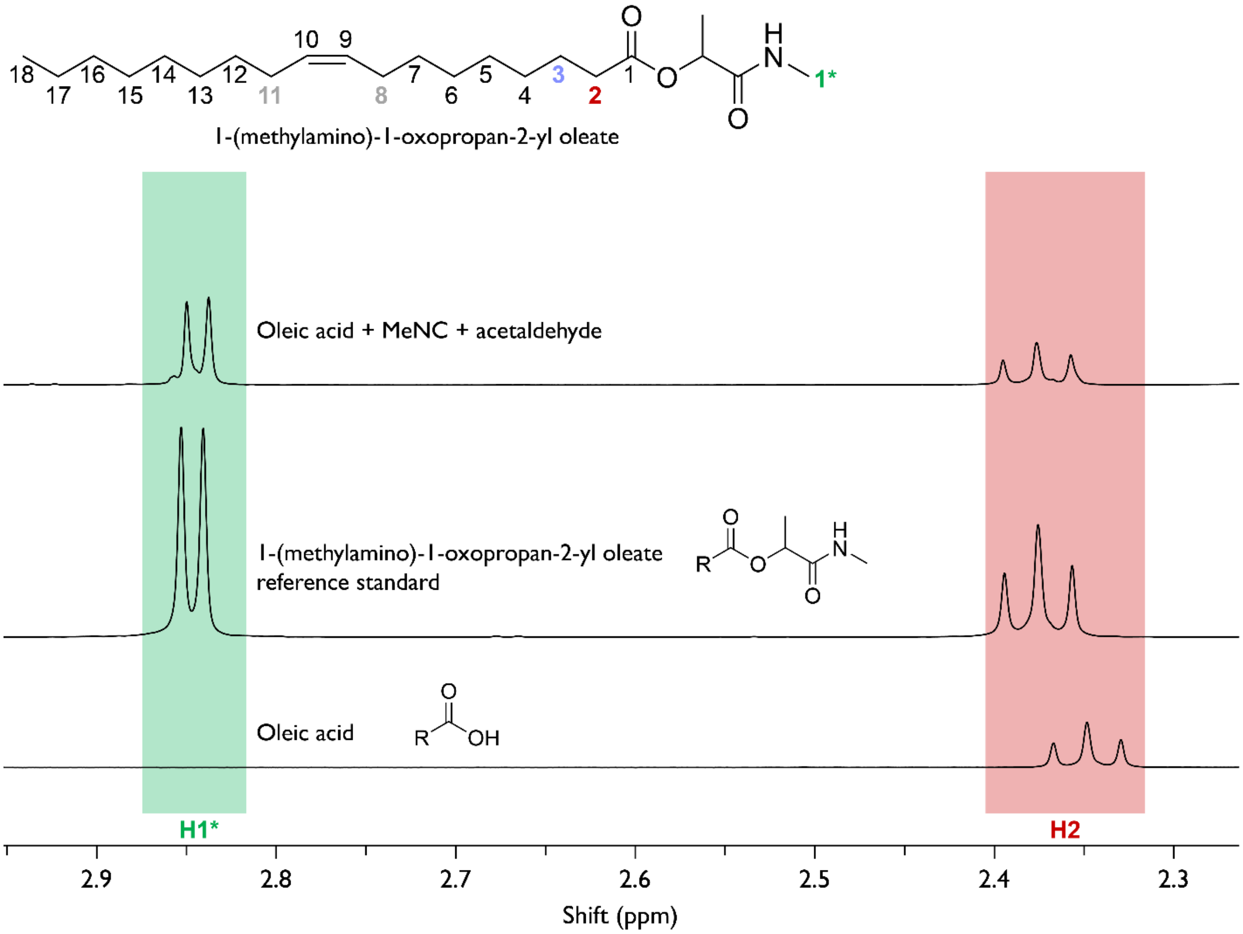


**Figure S61.** $^1H$ NMR spectrum of lipophilic products extracted from oleic acid vesicles incubated with 200 mM MeNC and 200 mM acetaldehyde at pH 8.0 for 48 h at 18 °C, followed by chloroform extraction, lyophilization, dissolution in $CDCl_3$, and analysis at 400 MHz. The spectrum is plotted alongside the reference standard for 1-(methylamino)-1-oxopropan-2-yl oleate, the expected Passerini reaction product. Diagnostic proton attached to C2 is highlighted in red, while proton from the adduct is highlighted in green. The dominant spectral features are consistent with formation of the α-acyloxyamide product (1-(methylamino)-1-oxopropan-2-yl oleate) derived from oleic acid under isonitrile–aldehyde activation conditions.

**Figure S62.**

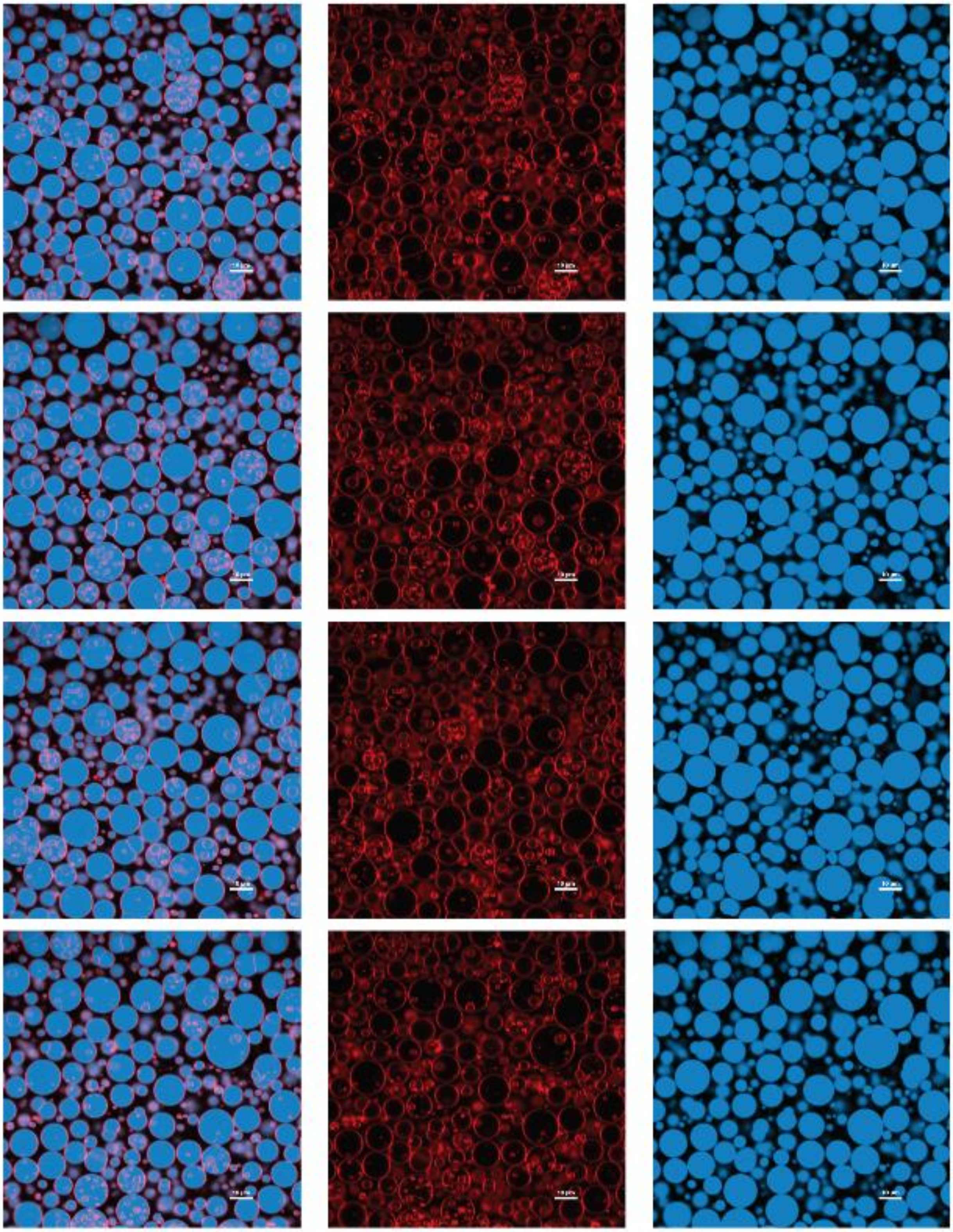

**Figure S62.** Example confocal images of oleic acid vesicles with 50 mM MeNC and 50 mM acetaldehyde after 5 minutes. Three columns represent overlay of red (Rhodamine B – lipid bilayer membrane signal) and blue (Cyanine 5 12 nt RNA) channels as well as their individual RGB channel signals. The scale bar represents 20 μm in length.

**Figure S63.**

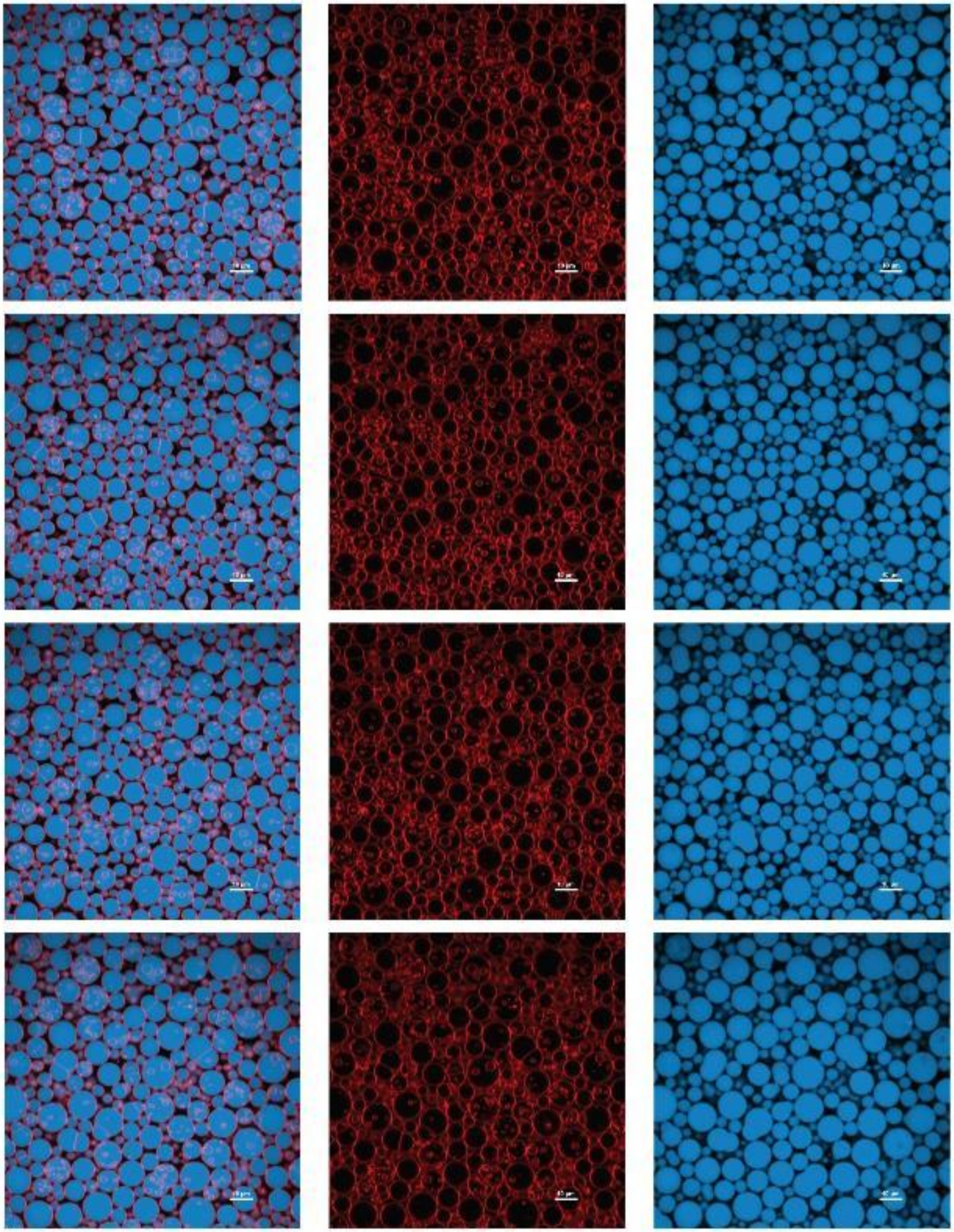

**Figure S63.** Example confocal images of oleic acid vesicles with 50 mM MeNC and 50 mM acetaldehyde after 24 hours. Three columns represent overlay of red (Rhodamine B – lipid bilayer membrane signal) and blue (Cyanine 5 12 nt RNA) channels as well as their individual RGB channel signals. The scale bar represents 20 μm in length.

**Figure S64.**

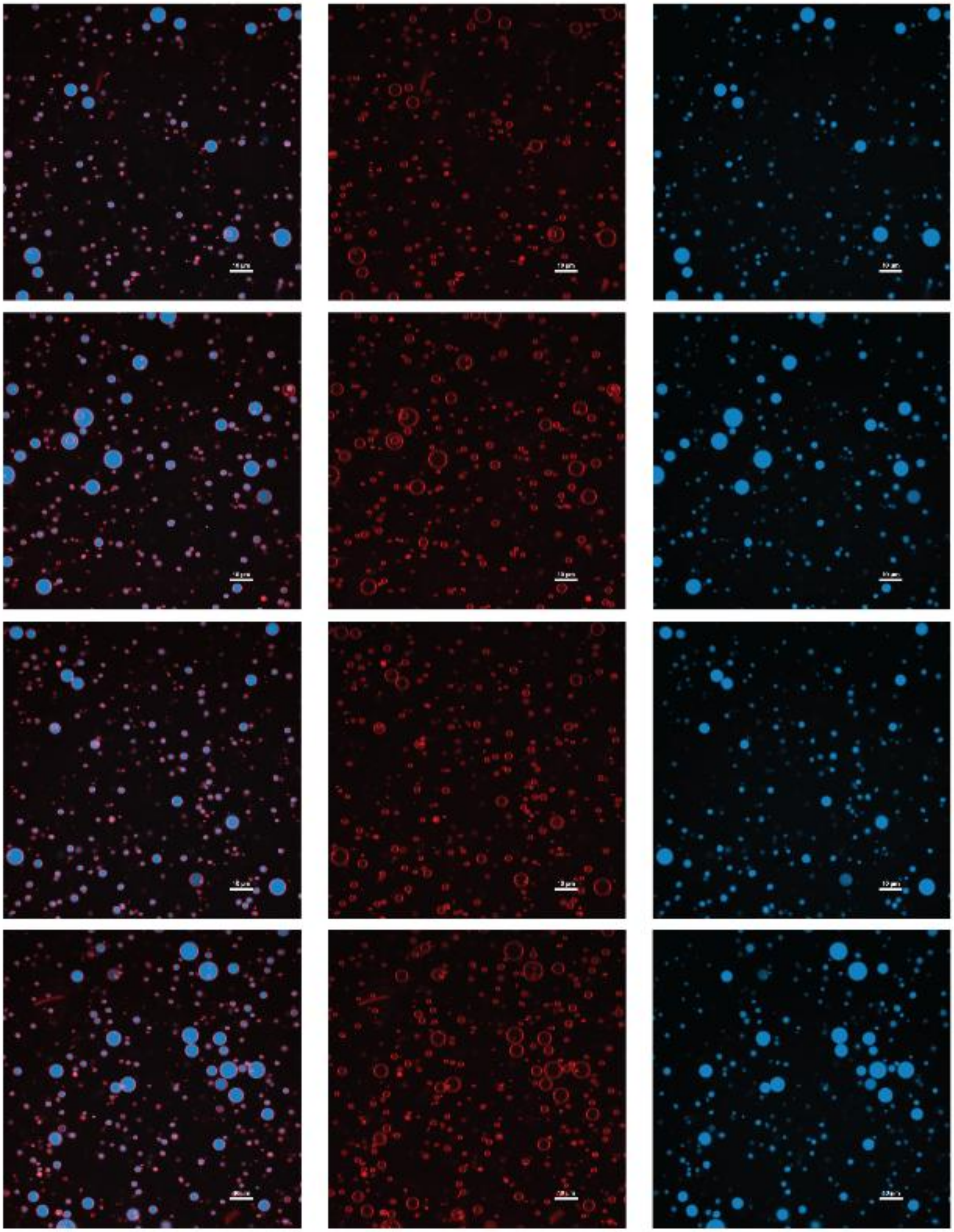

**Figure S64.** Example confocal images of oleic acid vesicles with 50 mM MeNC and 50 mM acetaldehyde after 72 hours. Three columns represent overlay of red (Rhodamine B – lipid bilayer membrane signal) and blue (Cyanine 5 12 nt RNA) channels as well as their individual RGB channel signals. The scale bar represents 20 μm in length.

**Figure S65.**

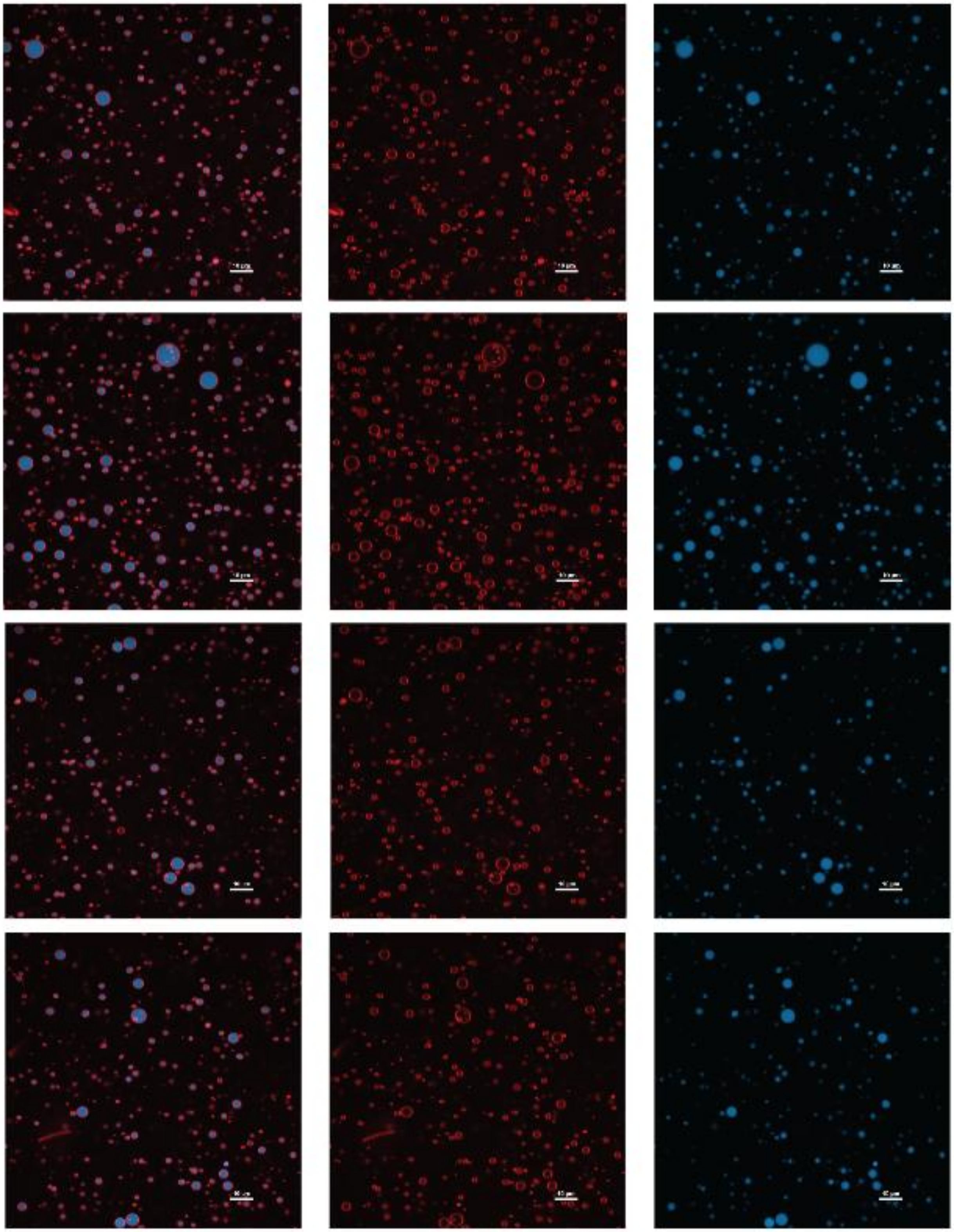

**Figure S65.** Example confocal images of oleic acid vesicles with 50 mM MeNC and 50 mM acetaldehyde after 168 hours. Three columns represent overlay of red (Rhodamine B – lipid bilayer membrane signal) and blue (Cyanine 5 12 nt RNA) channels as well as their individual RGB channel signals. The scale bar represents 20 μm in length.

**Figure S66.**

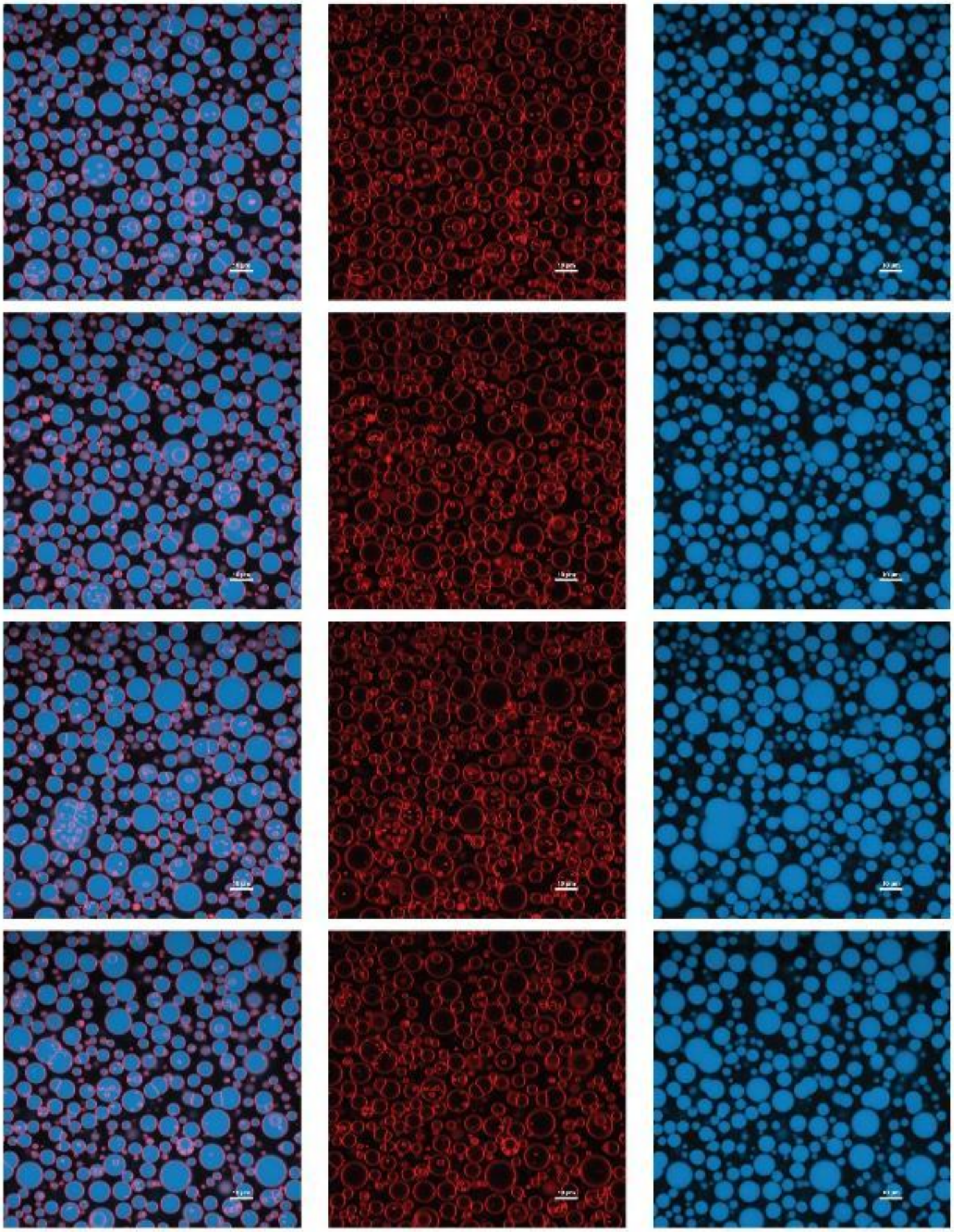

**Figure S66.** Example confocal images of oleic acid vesicles with 200 mM MeNC and 200 mM acetaldehyde after 5 minutes. Three columns represent overlay of red (Rhodamine B – lipid bilayer membrane signal) and blue (Cyanine 5 12 nt RNA) channels as well as their individual RGB channel signals. The scale bar represents 20 μm in length.

**Figure S67.**

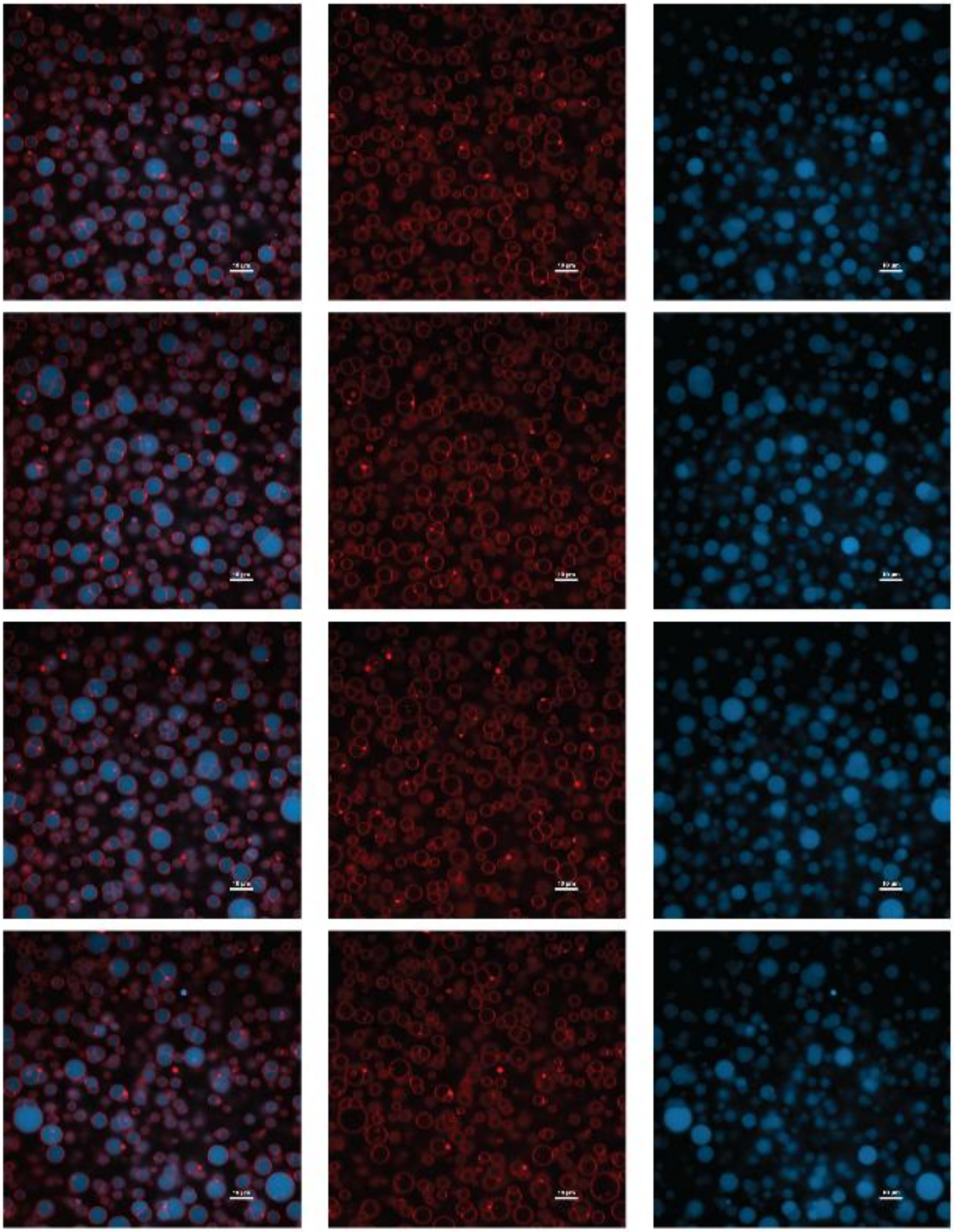

**Figure S67.** Example confocal images of oleic acid vesicles with 200 mM MeNC and 200 mM acetaldehyde after 24 hours. Three columns represent overlay of red (Rhodamine B – lipid bilayer membrane signal) and blue (Cyanine 5 12 nt RNA) channels as well as their individual RGB channel signals. The scale bar represents 20 μm in length.

**Figure S68.**

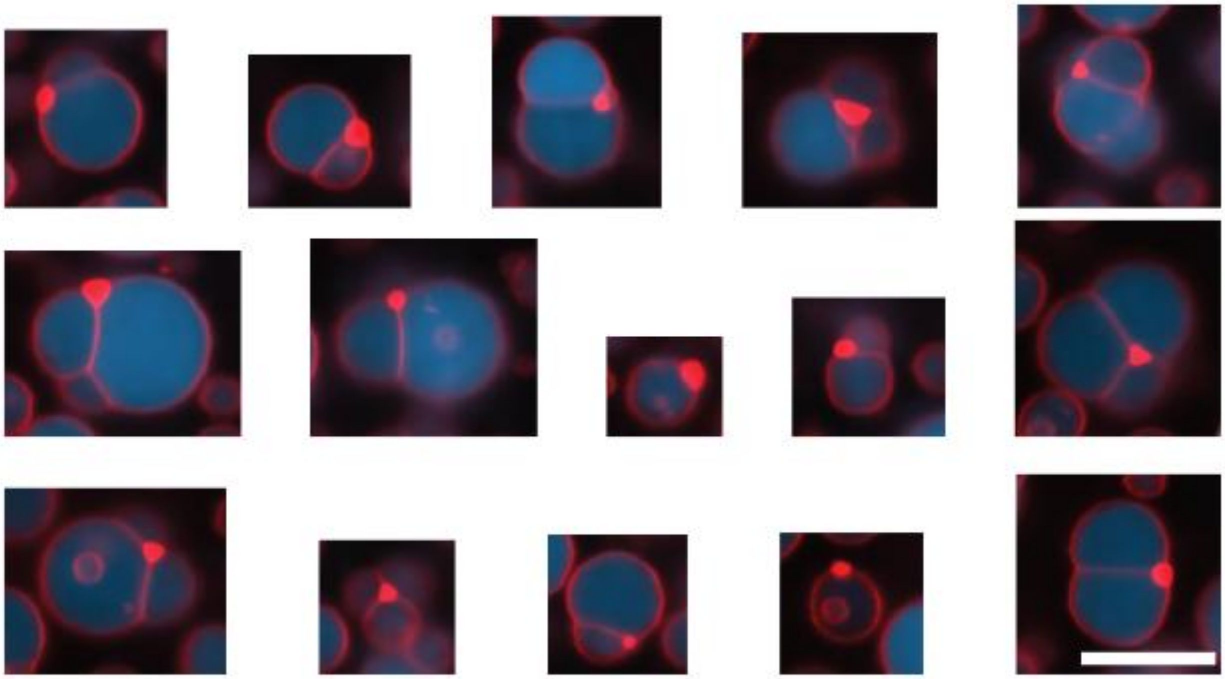

**Figure S68.** Zoomed-in confocal images of droplet-embedded vesicles (DBE) from 200 mM MeNC and 200 mM acetaldehyde after 24 hours images. DBE are made by conversion of oleic acid to its Passerini product of MeNC and acetaldehyde that is expected to be less soluble and form oil droplets (red dots on the images). Three columns represent overlay of red (Rhodamine B – lipid bilayer membrane signal) and blue (Cyanine 5 12 nt RNA) channels as well as their individual RGB channel signals. The scale bar represents 10 µm in length.

**Figure S69.**

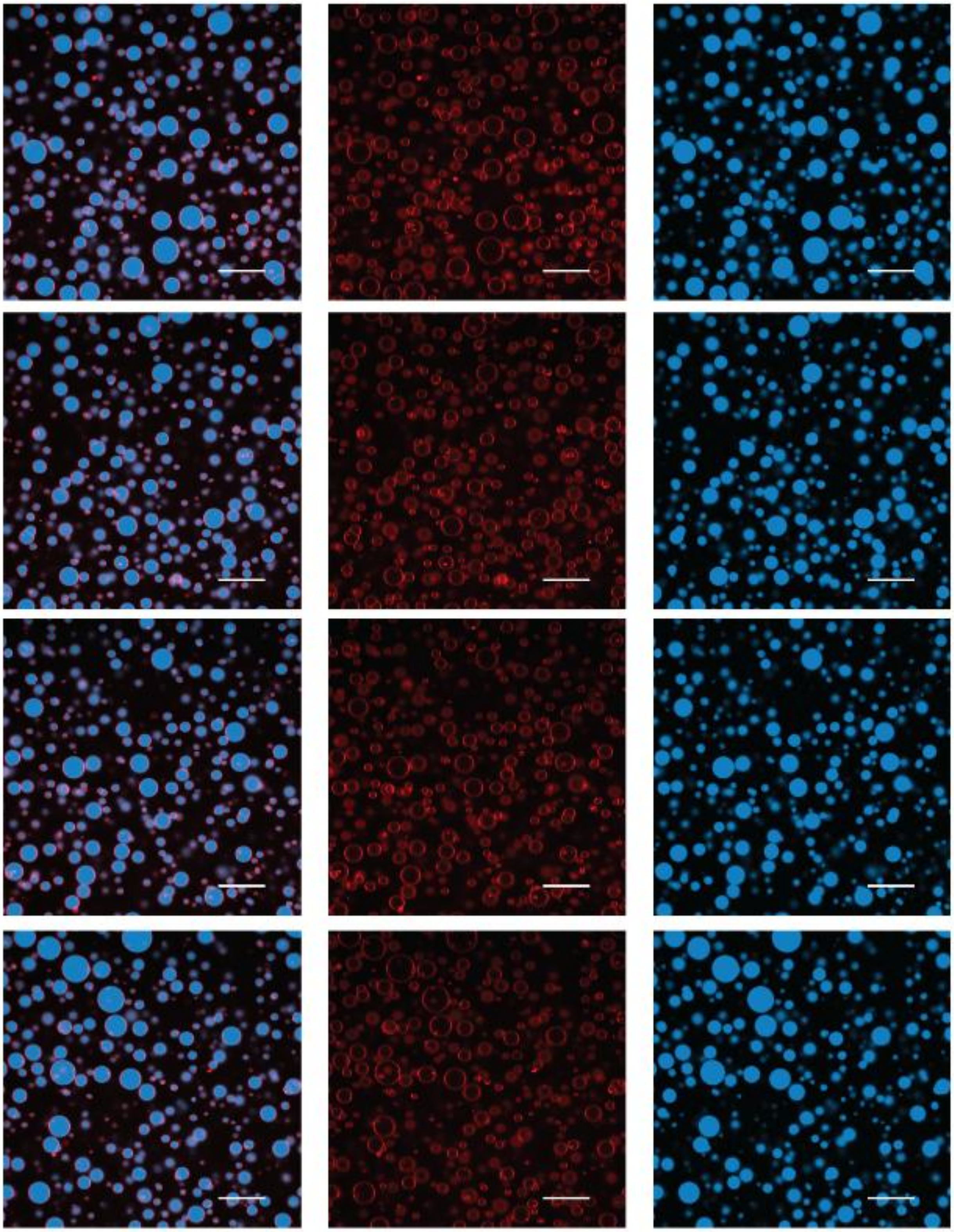

**Figure S69.** Example confocal images of oleic acid vesicles with 50 mM NCI after 5 minutes. Three columns represent overlay of red (Rhodamine B – lipid bilayer membrane signal) and blue (Cyanine 5 12 nt RNA) channels as well as their individual RGB channel signals. The scale bar represents 20 μm in length.

**Figure S70.**

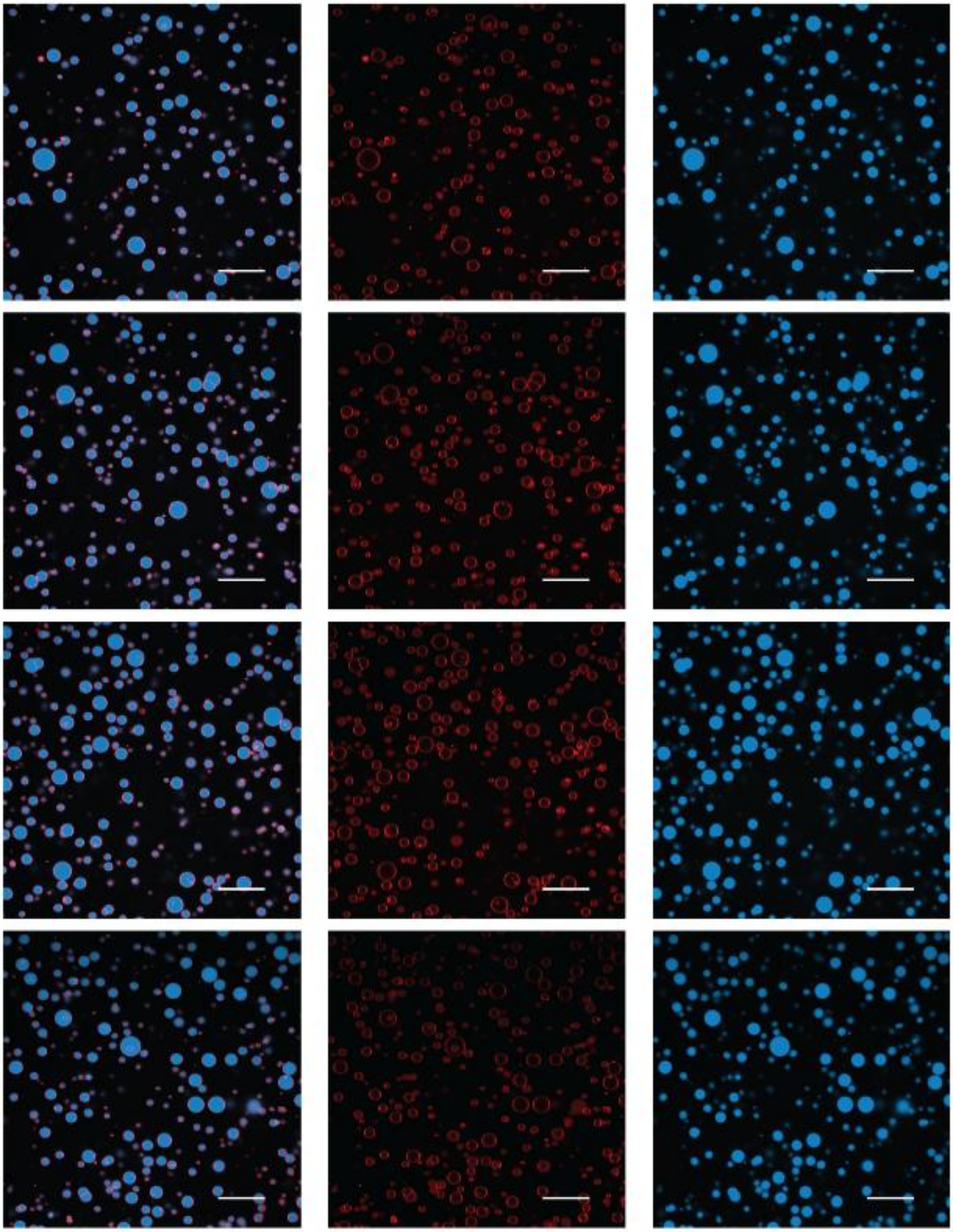

**Figure S70.** Example confocal images of oleic acid vesicles with 50 mM NCI after 2 hours. Three columns represent overlay of red (Rhodamine B – lipid bilayer membrane signal) and blue (Cyanine 5 12 nt RNA) channels as well as their individual RGB channel signals. The scale bar represents 20 μm in length.

**Figure S71.**

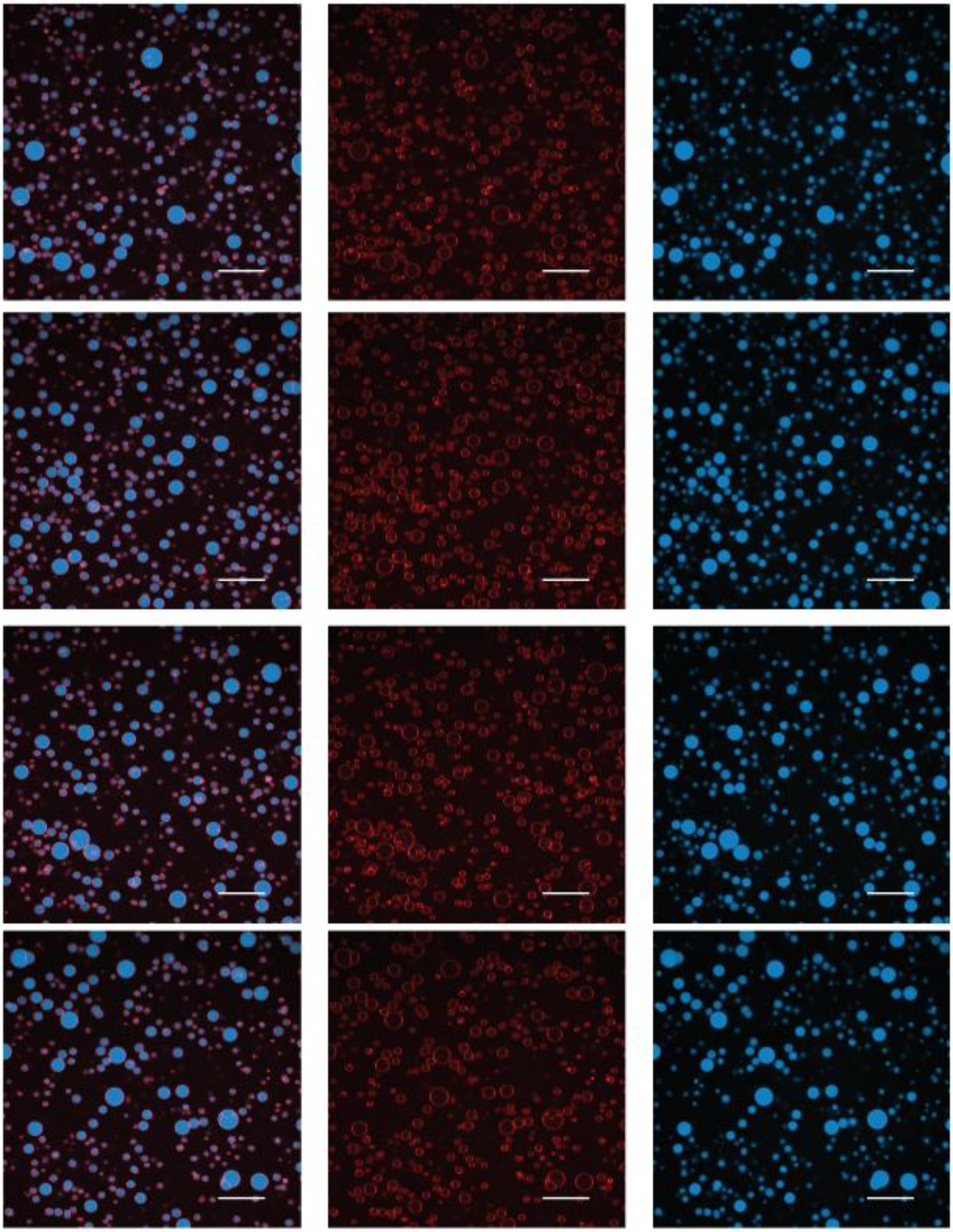

**Figure S71.** Example confocal images of oleic acid vesicles with 50 mM NCI after 24 hours. Three columns represent overlay of red (Rhodamine B – lipid bilayer membrane signal) and blue (Cyanine 5 12 nt RNA) channels as well as their individual RGB channel signals. The scale bar represents 20 µm in length.

**Figure S72.**

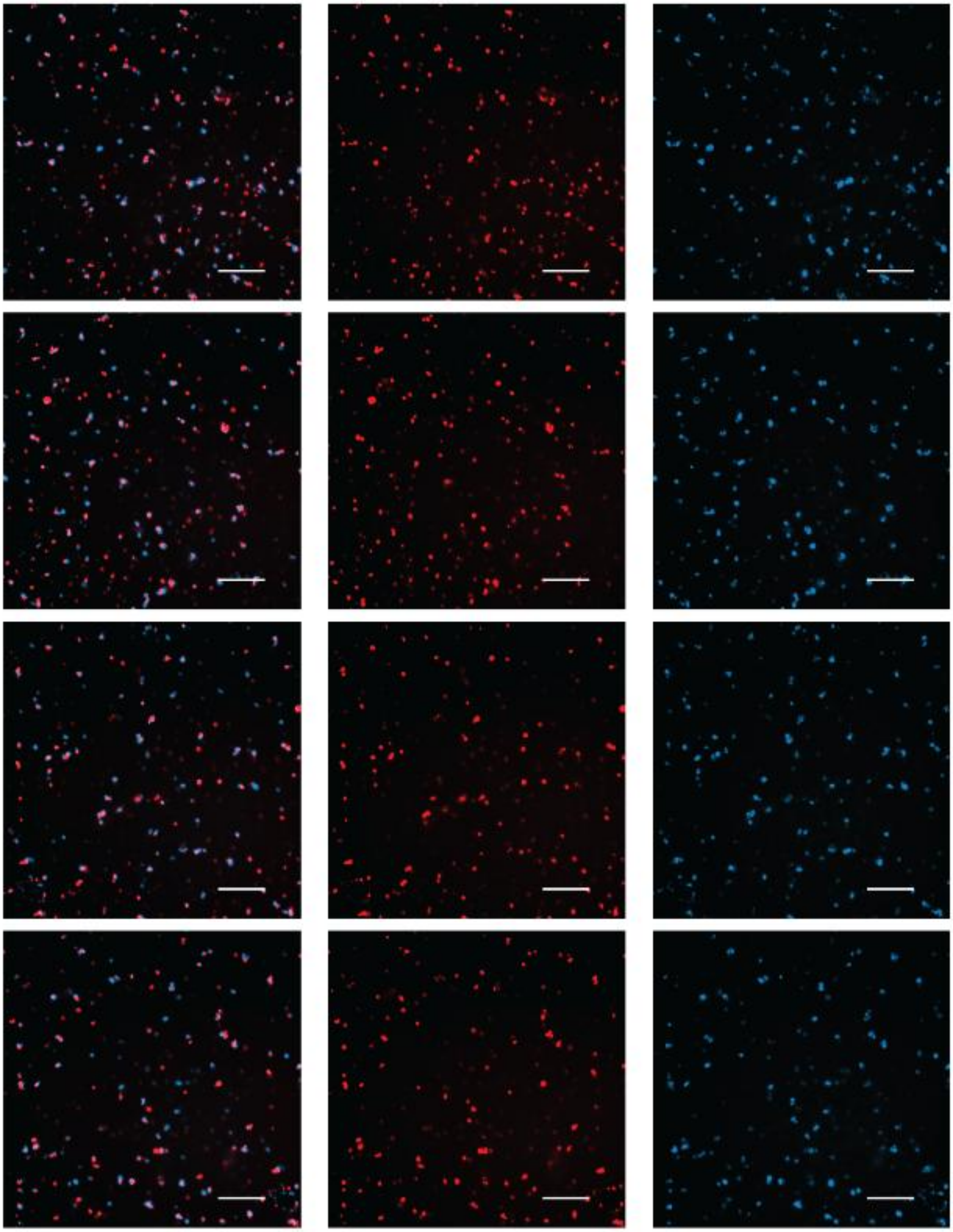

**Figure S72.** Example confocal images of oleic acid vesicles with 200 mM NCI after 0 hours. Three columns represent overlay of red (Rhodamine B – lipid bilayer membrane signal) and blue

(Cyanine 5 12 nt RNA) channels as well as their individual RGB channel signals. The scale bar represents 20 μm in length.

**Figure S73.**

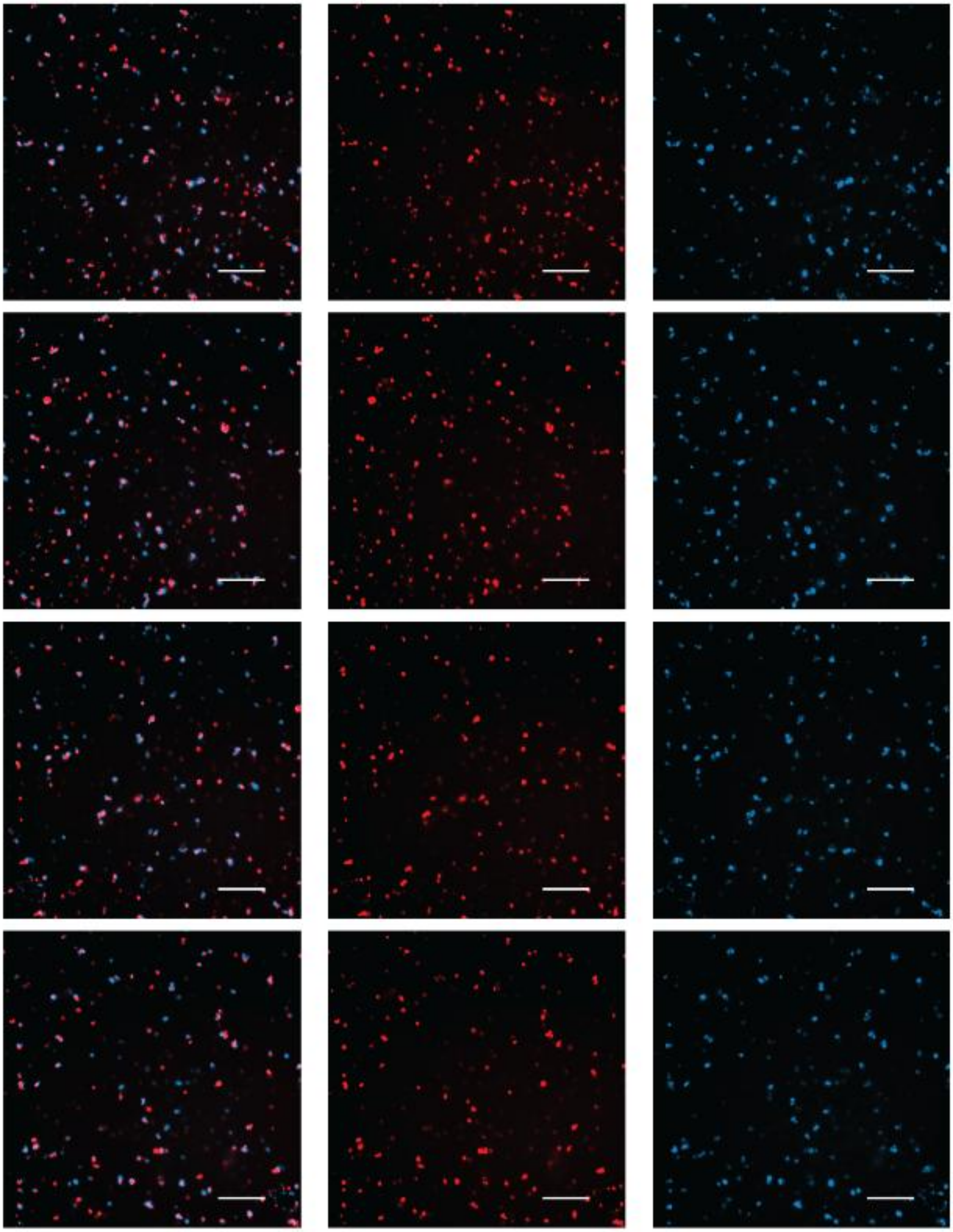

**Figure S73.** Example confocal images of oleic acid vesicles with 200 mM NCI after 2 hours. Three columns represent overlay of red (Rhodamine B – lipid bilayer membrane signal) and blue (Cyanine 5 12 nt RNA) channels as well as their individual RGB channel signals. The scale bar represents 20 μm in length.

**Figure S74.**

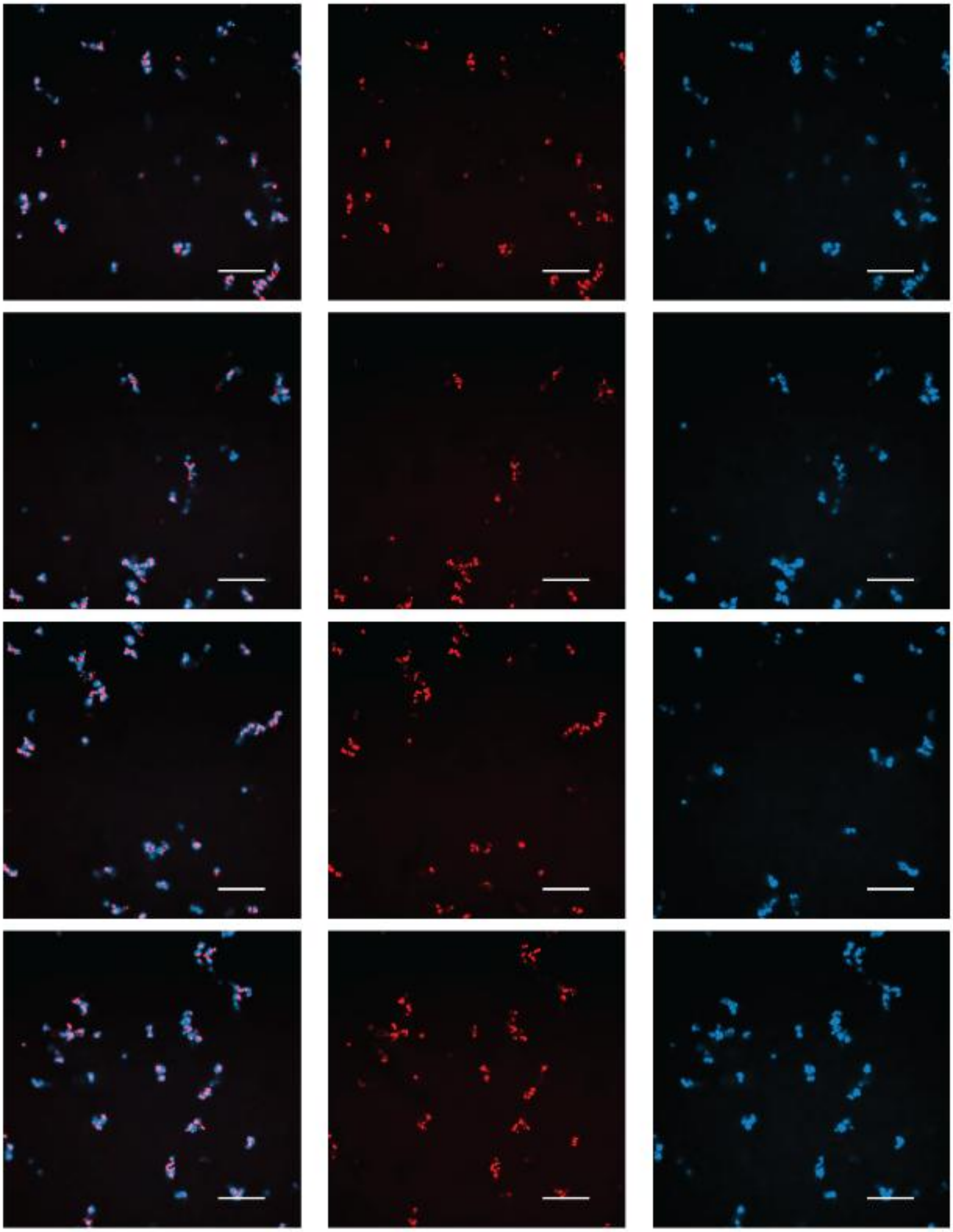

**Figure S74.** Example confocal images of oleic acid vesicles with 200 mM NCI after 24 hours. Three columns represent overlay of red (Rhodamine B – lipid bilayer membrane signal) and blue (Cyanine 5 12 nt RNA) channels as well as their individual RGB channel signals. The scale bar represents 20 μm in length.

**Figure S75.**

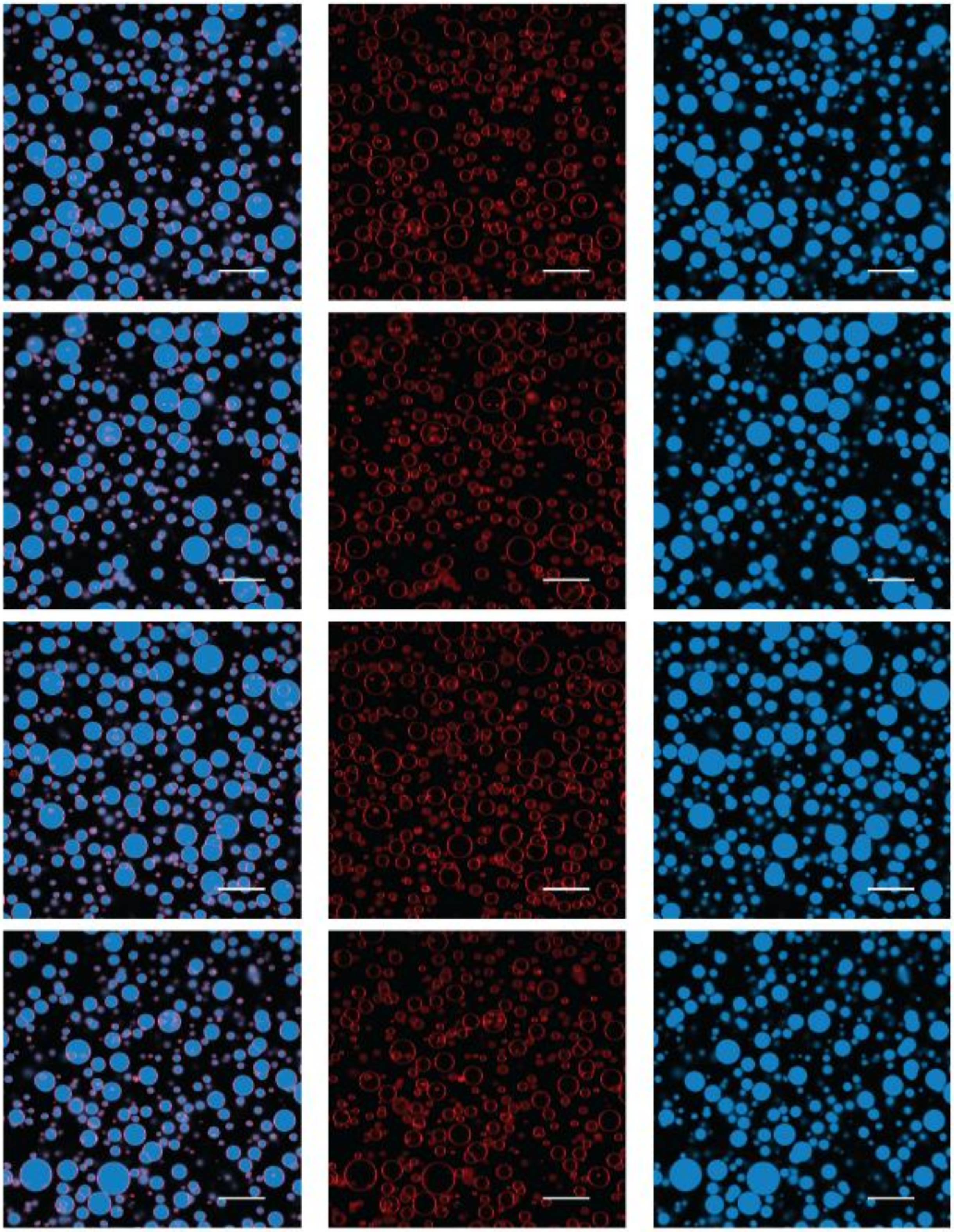

**Figure S75.** Example confocal images of oleic acid vesicles with 50 mM IDI after 0 hours. Three columns represent overlay of red (Rhodamine B – lipid bilayer membrane signal) and blue (Cyanine 5 12 nt RNA) channels as well as their individual RGB channel signals. The scale bar represents 20 μm in length.

**Figure S76.**

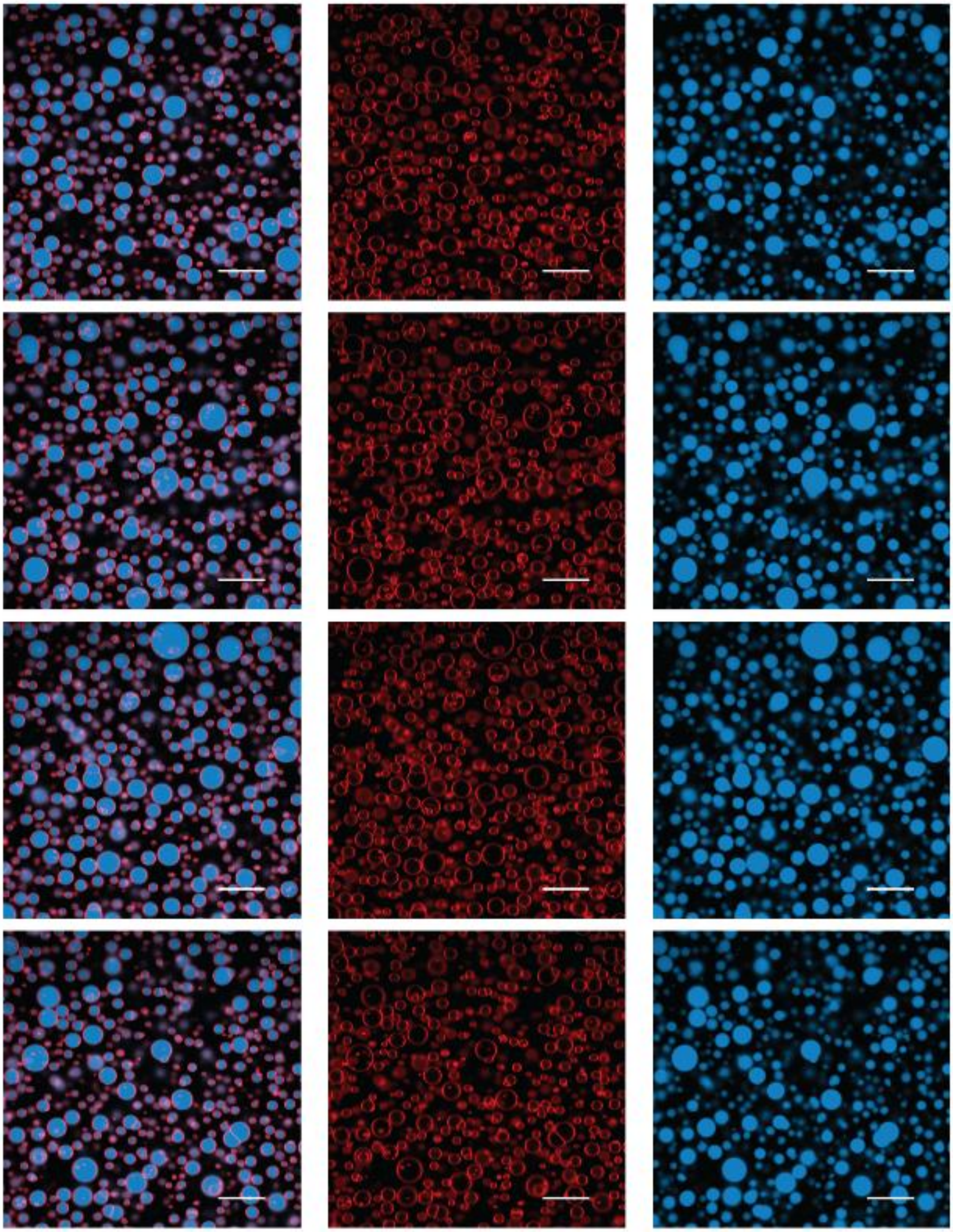

**Figure S76.** Example confocal images of oleic acid vesicles with 50 mM IDI after 2 hours. Three columns represent overlay of red (Rhodamine B – lipid bilayer membrane signal) and blue (Cyanine 5 12 nt RNA) channels as well as their individual RGB channel signals. The scale bar represents 20 μm in length.

**Figure S77.**

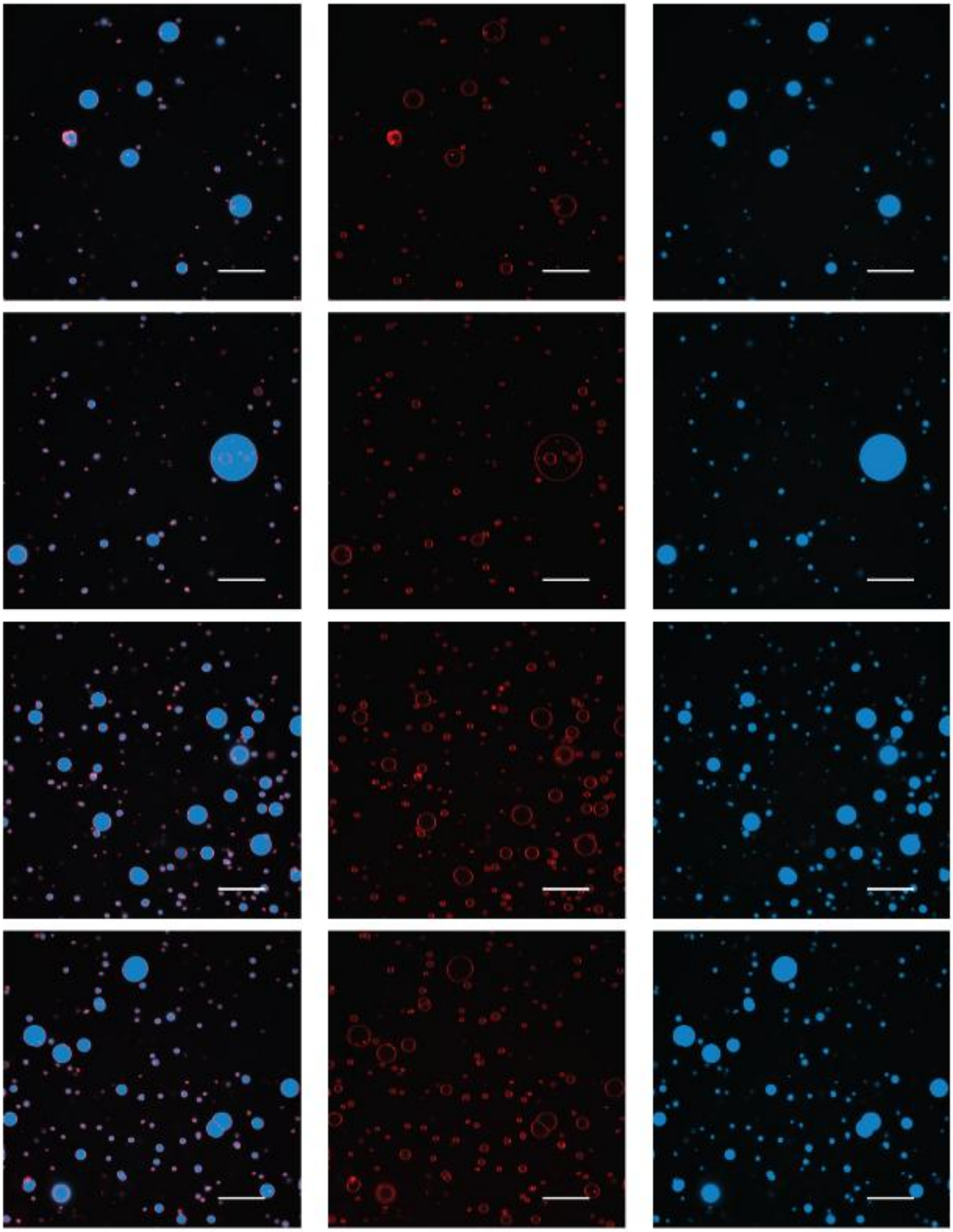

**Figure S77.** Example confocal images of oleic acid vesicles with 50 mM IDI after 24 hours. Three columns represent overlay of red (Rhodamine B – lipid bilayer membrane signal) and blue (Cyanine 5 12 nt RNA) channels as well as their individual RGB channel signals. The scale bar represents 20 μm in length.

**Figure S78.**

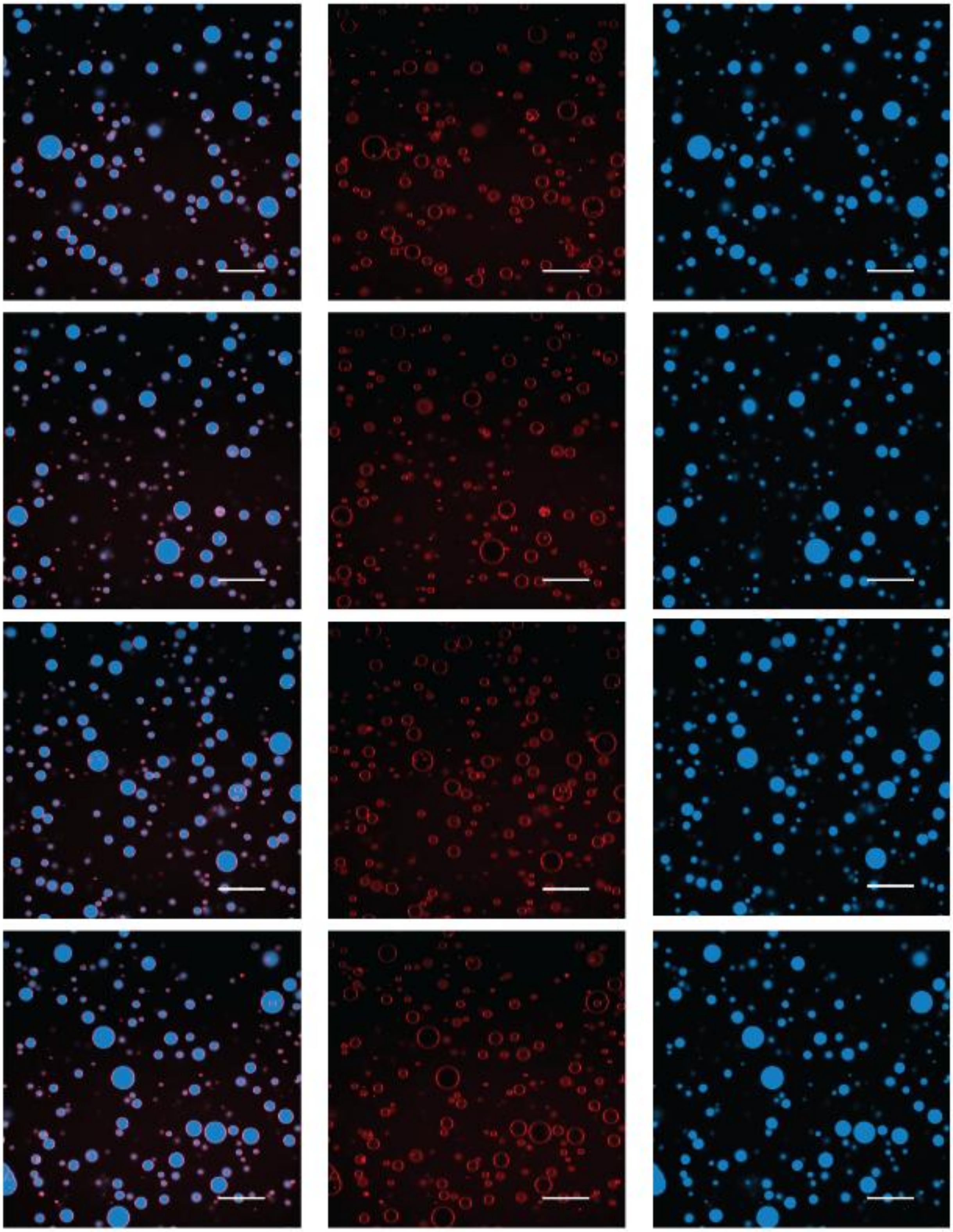

**Figure S78.** Example confocal images of oleic acid vesicles with 200 mM IDI after 0 hours. Three columns represent overlay of red (Rhodamine B – lipid bilayer membrane signal) and blue (Cyanine 5 12 nt RNA) channels as well as their individual RGB channel signals. The scale bar represents 20 μm in length.

**Figure S79.**

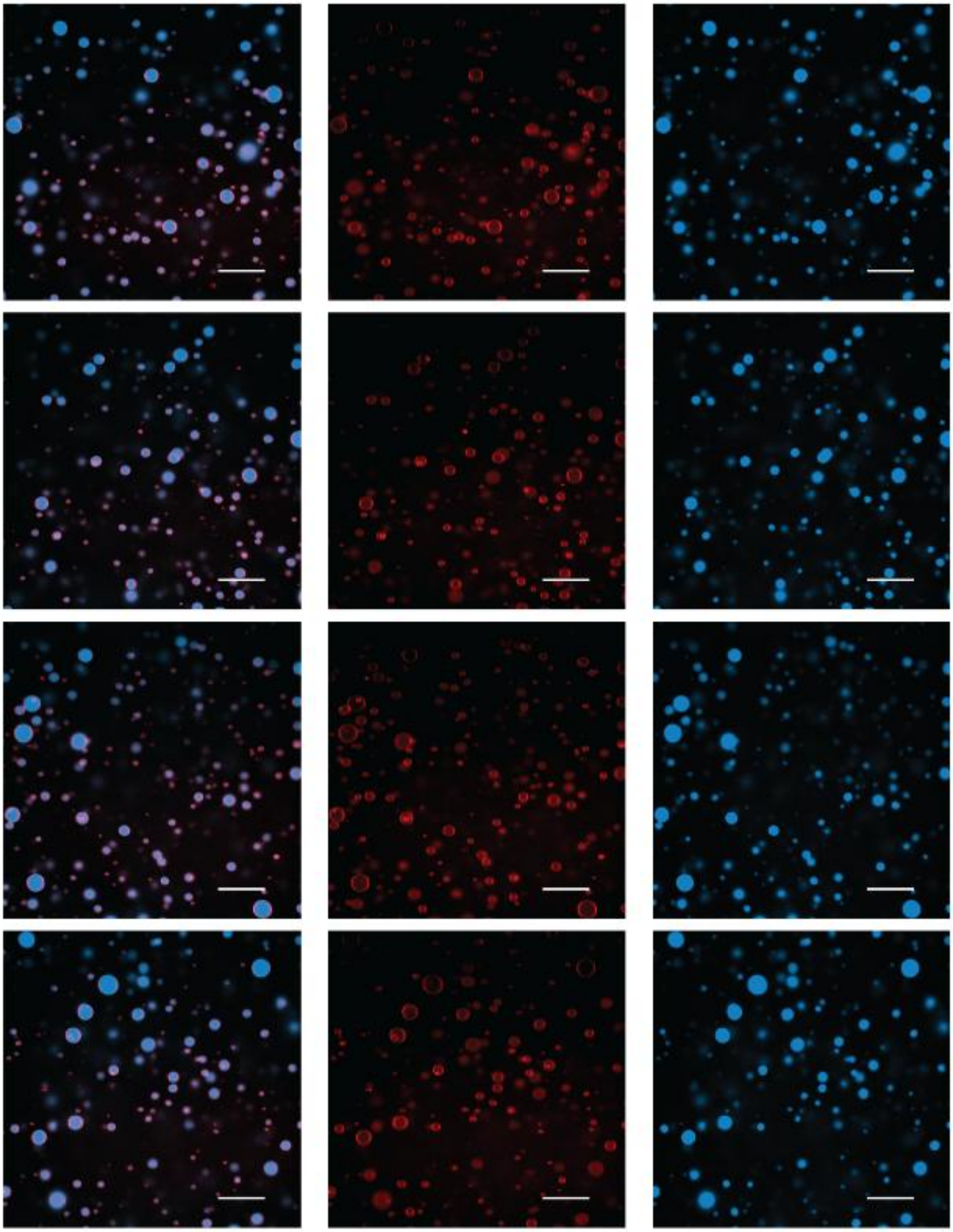

**Figure S79.** Example confocal images of oleic acid vesicles with 200 mM IDI after 2 hours. Three columns represent overlay of red (Rhodamine B – lipid bilayer membrane signal) and blue (Cyanine 5 12 nt RNA) channels as well as their individual RGB channel signals. The scale bar represents 20 μm in length.

**Figure S80.**

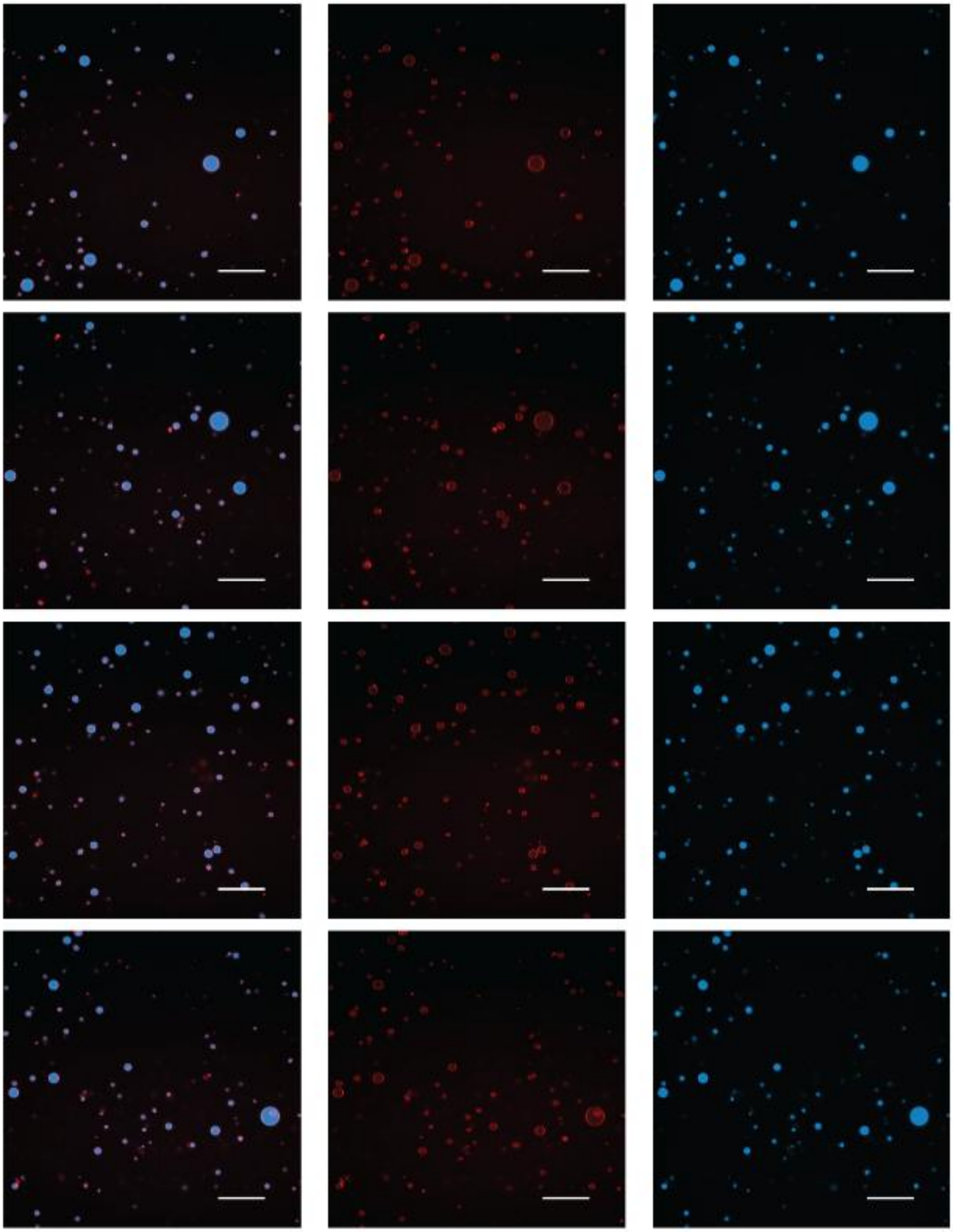

**Figure S80.** Example confocal images of oleic acid vesicles with 200 mM IDI after 24 hours. Three columns represent overlay of red (Rhodamine B – lipid bilayer membrane signal) and blue (Cyanine 5 12 nt RNA) channels as well as their individual RGB channel signals. The scale bar represents 20 μm in length.

**Figure S81.**

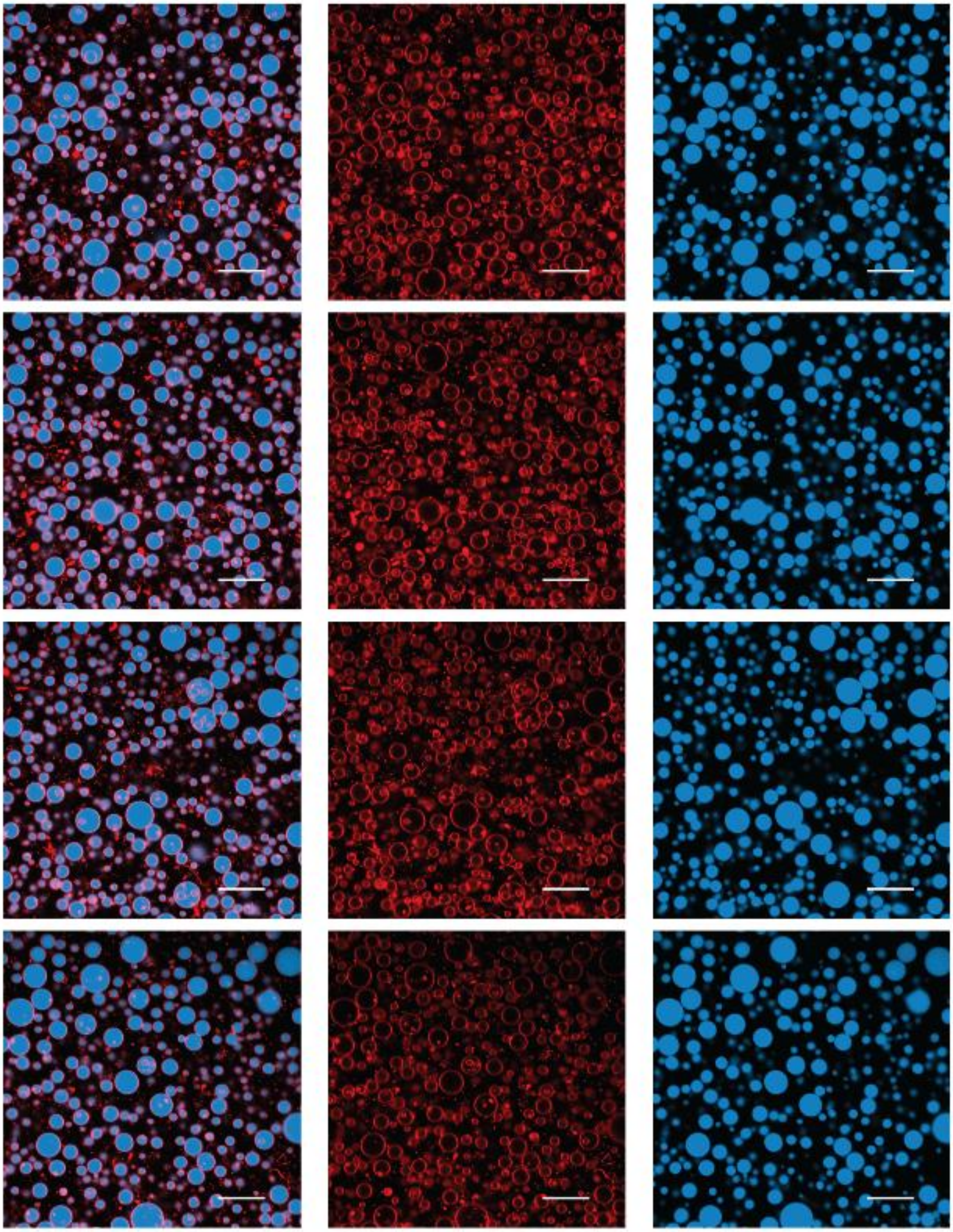

**Figure S81.** Example confocal images of oleic acid vesicles with 50 mM TCDI after 0 hours. Three columns represent overlay of red (Rhodamine B – lipid bilayer membrane signal) and blue (Cyanine 5 12 nt RNA) channels as well as their individual RGB channel signals. The scale bar represents 20 μm in length.

**Figure S82.**

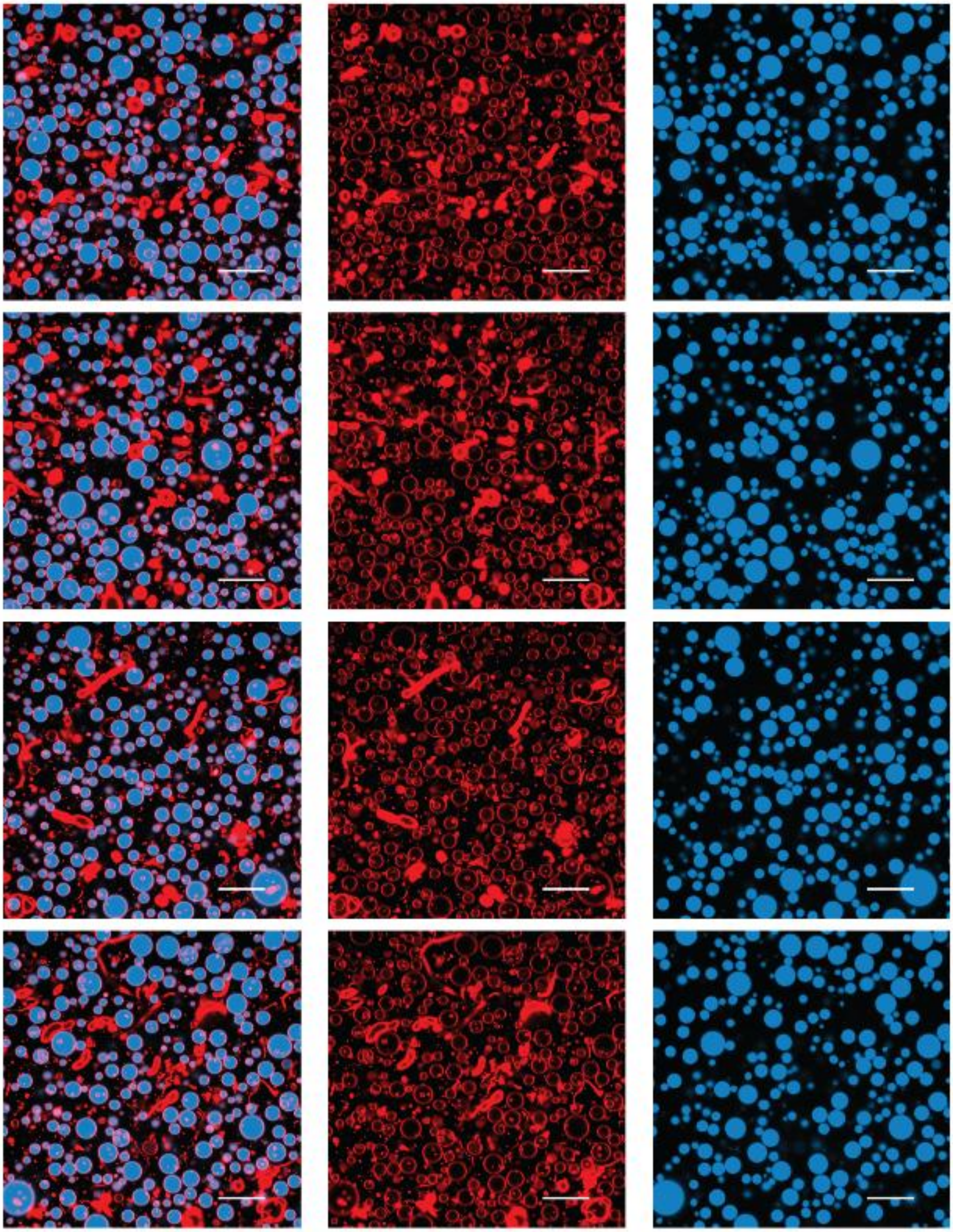

**Figure S82.** Example confocal images of oleic acid vesicles with 50 mM TCDI after 2 hours. Three columns represent overlay of red (Rhodamine B – lipid bilayer membrane signal) and blue (Cyanine 5 12 nt RNA) channels as well as their individual RGB channel signals. The scale bar represents 20 μm in length.

**Figure S83.**

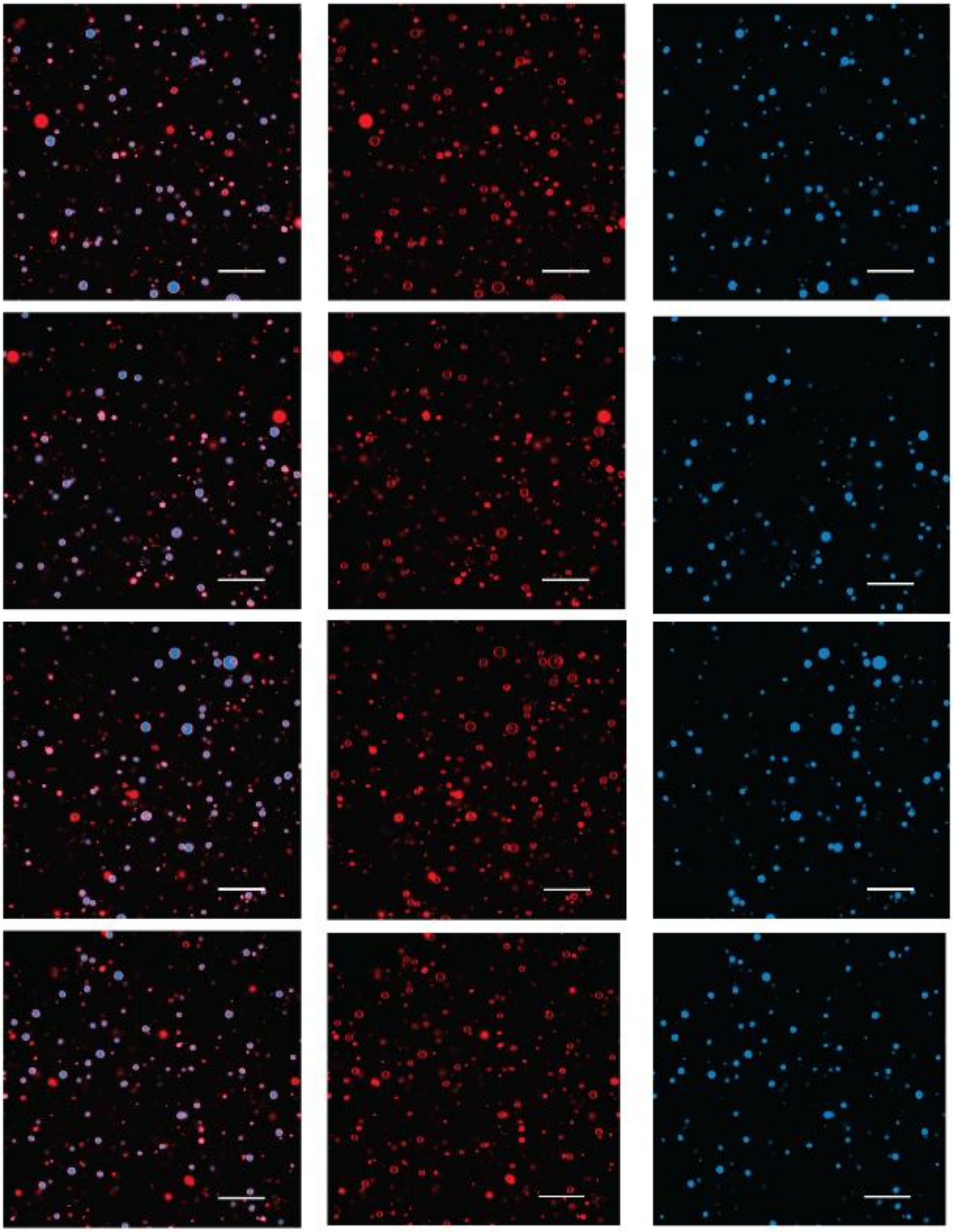

**Figure S83.** Example confocal images of oleic acid vesicles with 50 mM TCDI after 24 hours. Three columns represent overlay of red (Rhodamine B – lipid bilayer membrane signal) and blue (Cyanine 5 12 nt RNA) channels as well as their individual RGB channel signals. The scale bar represents 20 μm in length.

**Figure S84.**

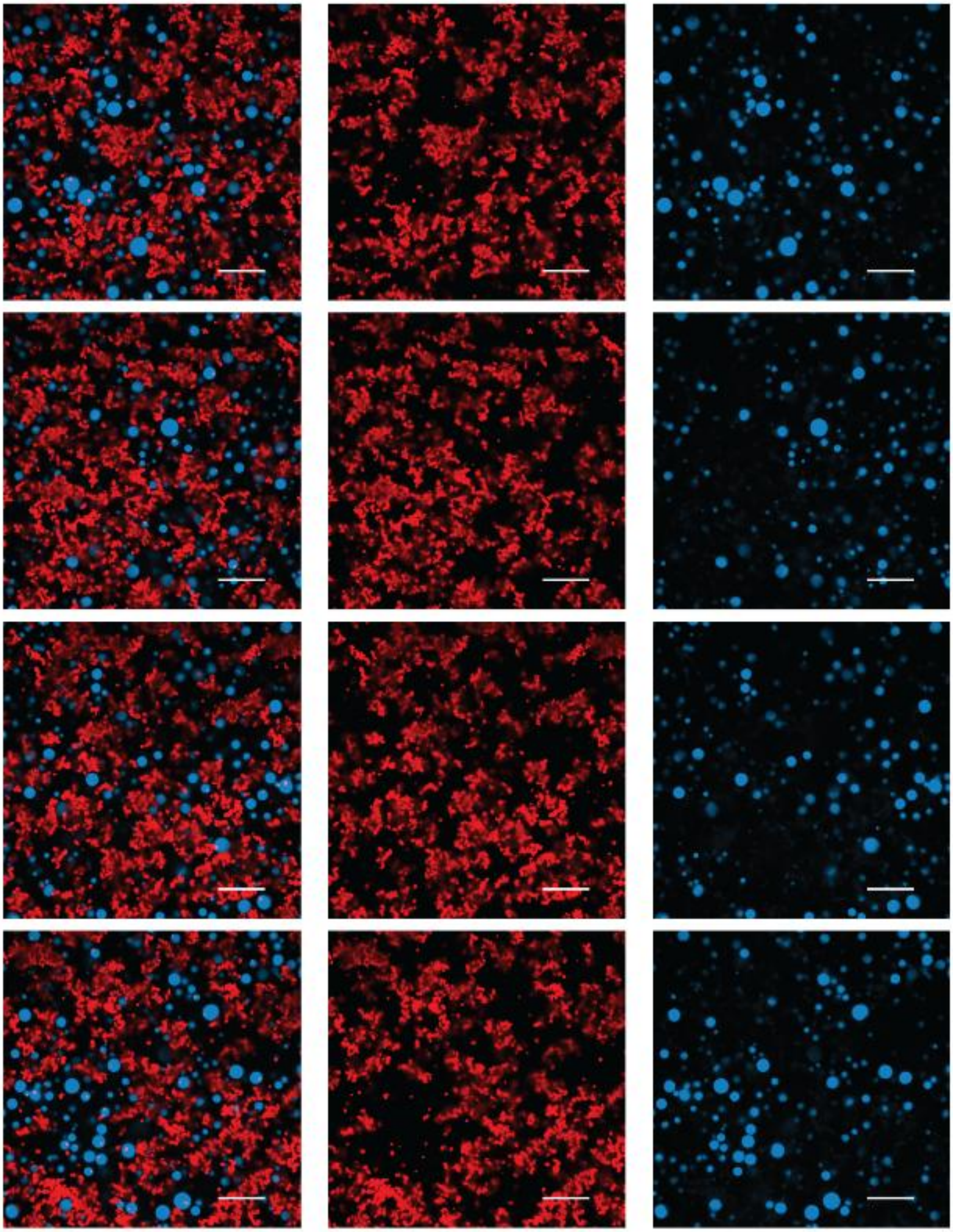

**Figure S84.** Example confocal images of oleic acid vesicles with 200 mM TCDI after 0 hours. Three columns represent overlay of red (Rhodamine B – lipid bilayer membrane signal) and blue (Cyanine 5 12 nt RNA) channels as well as their individual RGB channel signals. The scale bar represents 20 μm in length.

**Figure S85.**

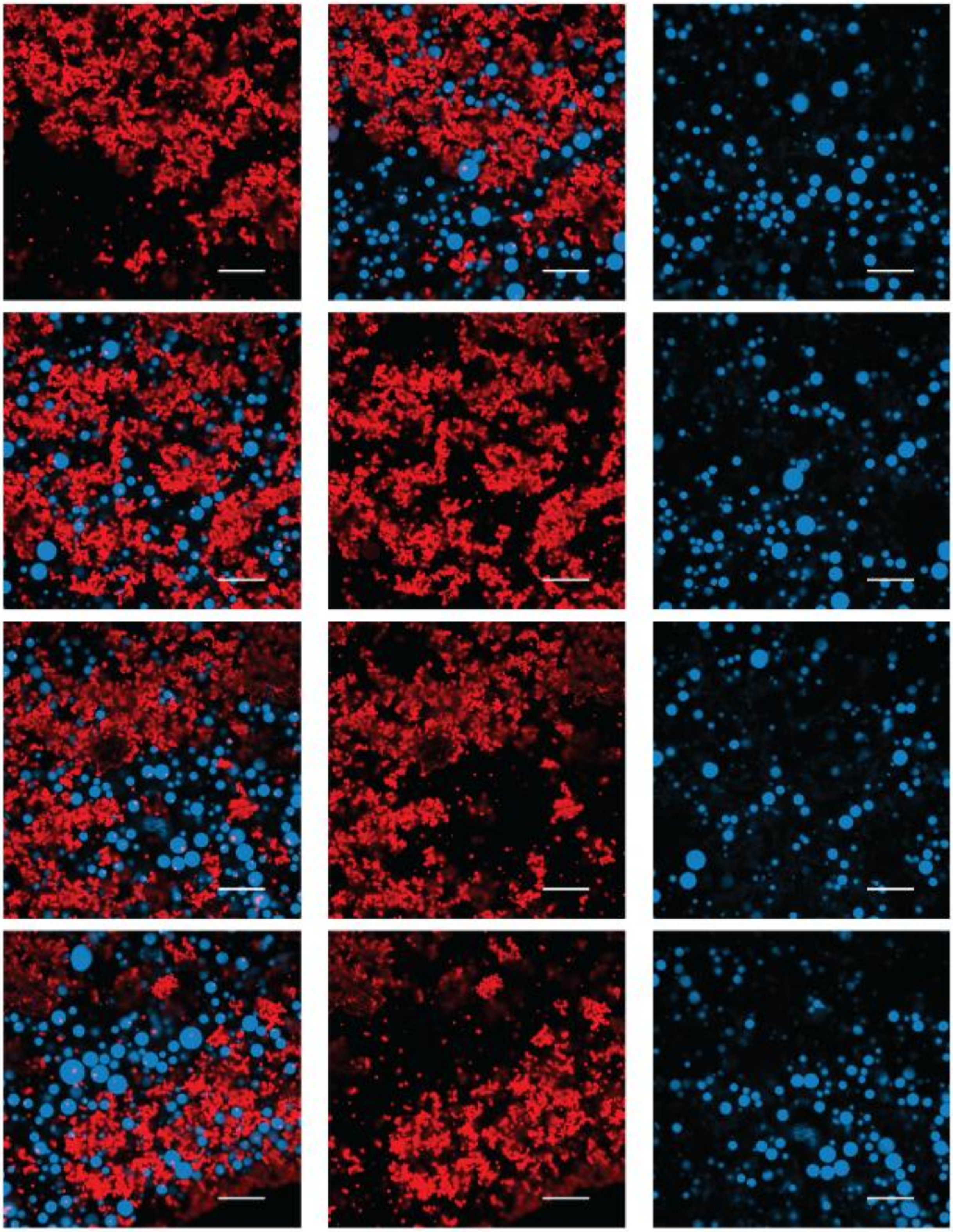

**Figure S85.** Example confocal images of oleic acid vesicles with 200 mM TCDI after 2 hours. Three columns represent overlay of red (Rhodamine B – lipid bilayer membrane signal) and blue (Cyanine 5 12 nt RNA) channels as well as their individual RGB channel signals. The scale bar represents 20 μm in length.

**Figure S86.**

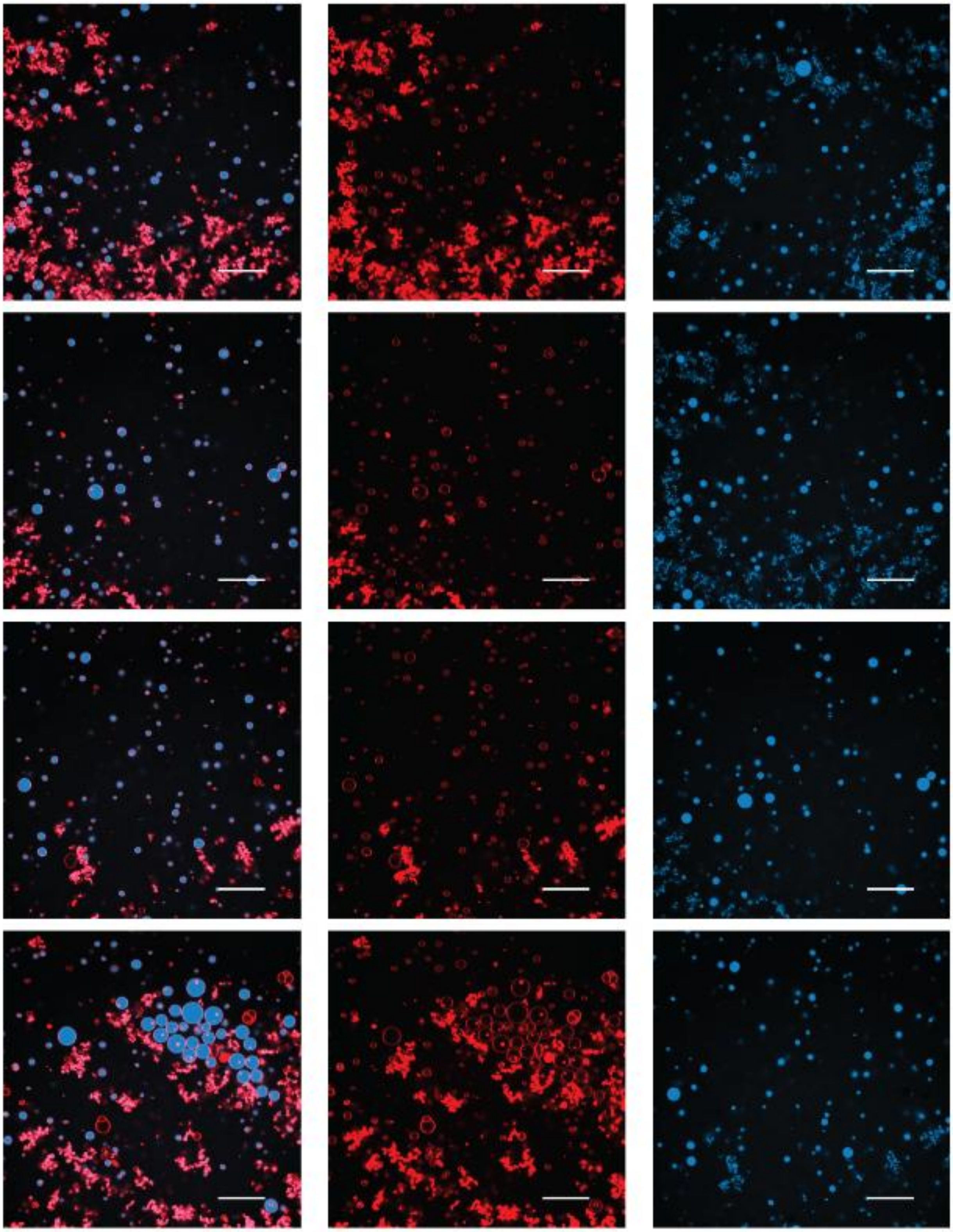

**Figure S86.** Example confocal images of oleic acid vesicles with 200 mM TCDI after 24 hours. Three columns represent overlay of red (Rhodamine B – lipid bilayer membrane signal) and blue (Cyanine 5 12 nt RNA) channels as well as their individual RGB channel signals. The scale bar represents 20 μm in length.

**Figure S87.**

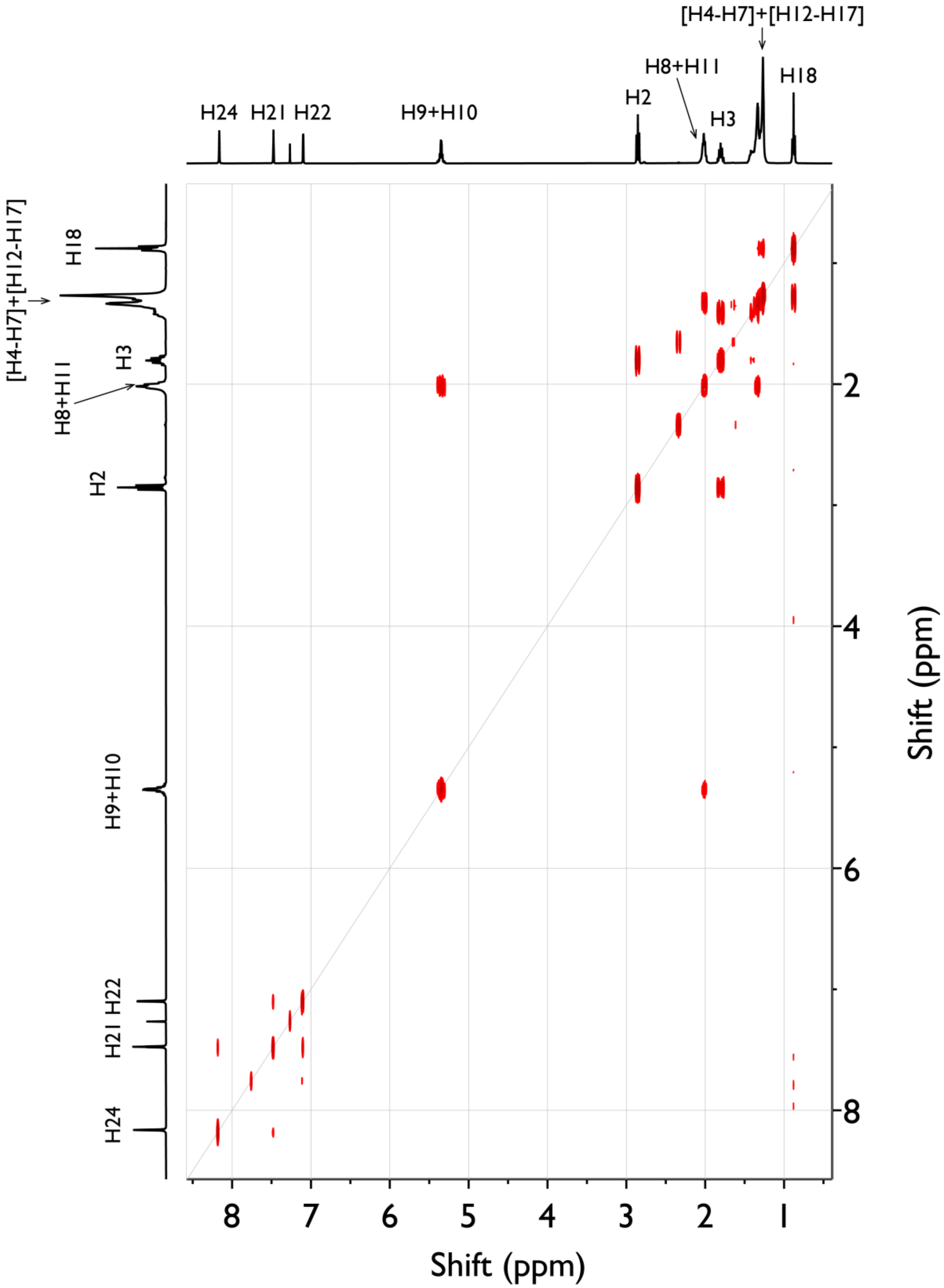

[H4-H7]+[H12-H17]
H8+H11
H18
H2
H3
H24
H21
H22
H9+H10
Shift (ppm)
Shift (ppm)

**Figure S87.** $^1H$–$^1H$ COSY NMR spectrum of *(Z)-1-(1H-imidazol-1-yl)octadec-9-en-1-one* recorded in $CDCl_3$ at 400 MHz, showing through-bond proton-proton correlations consistent with the assigned structure.

**Figure S88.**

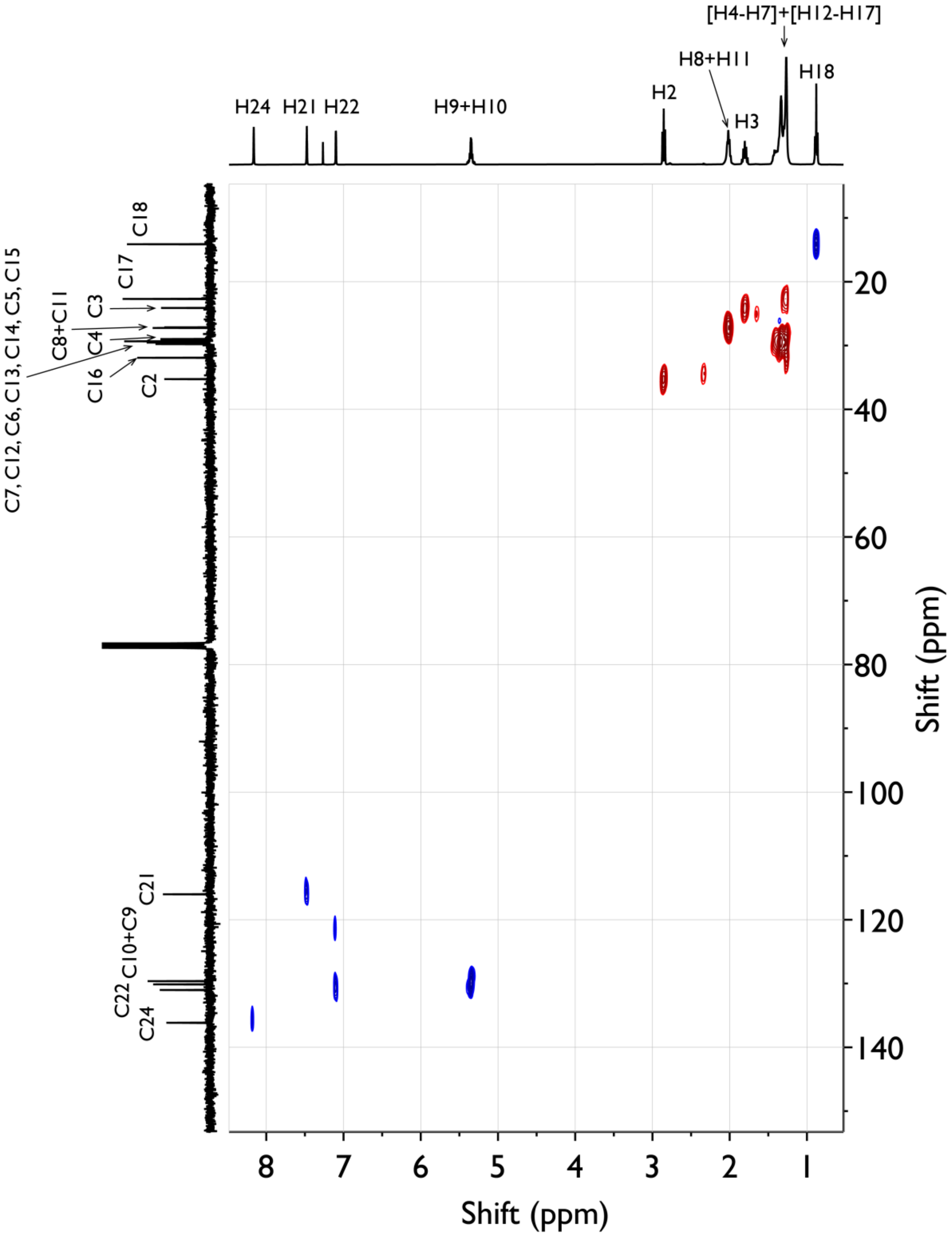


**Figure S88.** $^{13}C$–$^{1}H$ HSQC NMR spectrum of *(Z)-1-(1H-imidazol-1-yl)octadec-9-en-1-one* recorded in $CDCl_3$ at 400 MHz, showing one-bond $^{13}C$–$^{1}H$ correlations that confirm the assignment of protonated carbon centers within the alkenyl chain and the imidazolyl carbonyl moiety.

**Figure S89.**

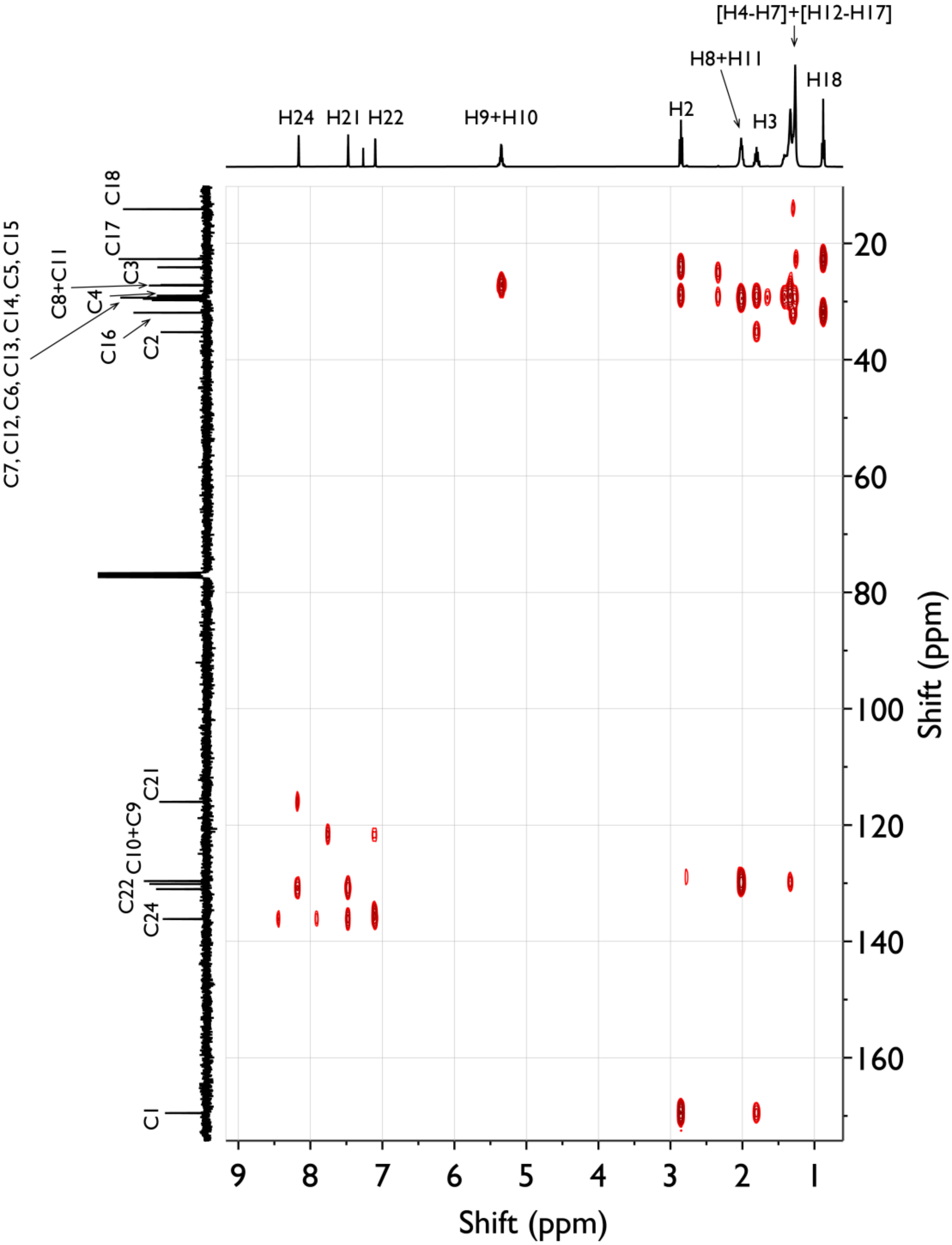


**Figure S89.** $^{1}H$–$^{13}C$ HMBC NMR spectrum of *(Z)-1-(1H-imidazol-1-yl)octadec-9-en-1-one* recorded in $CDCl_3$ at 400 MHz, showing long-range two- and three-bond $^{1}H$–$^{13}C$ correlations.

**Figure S90.**

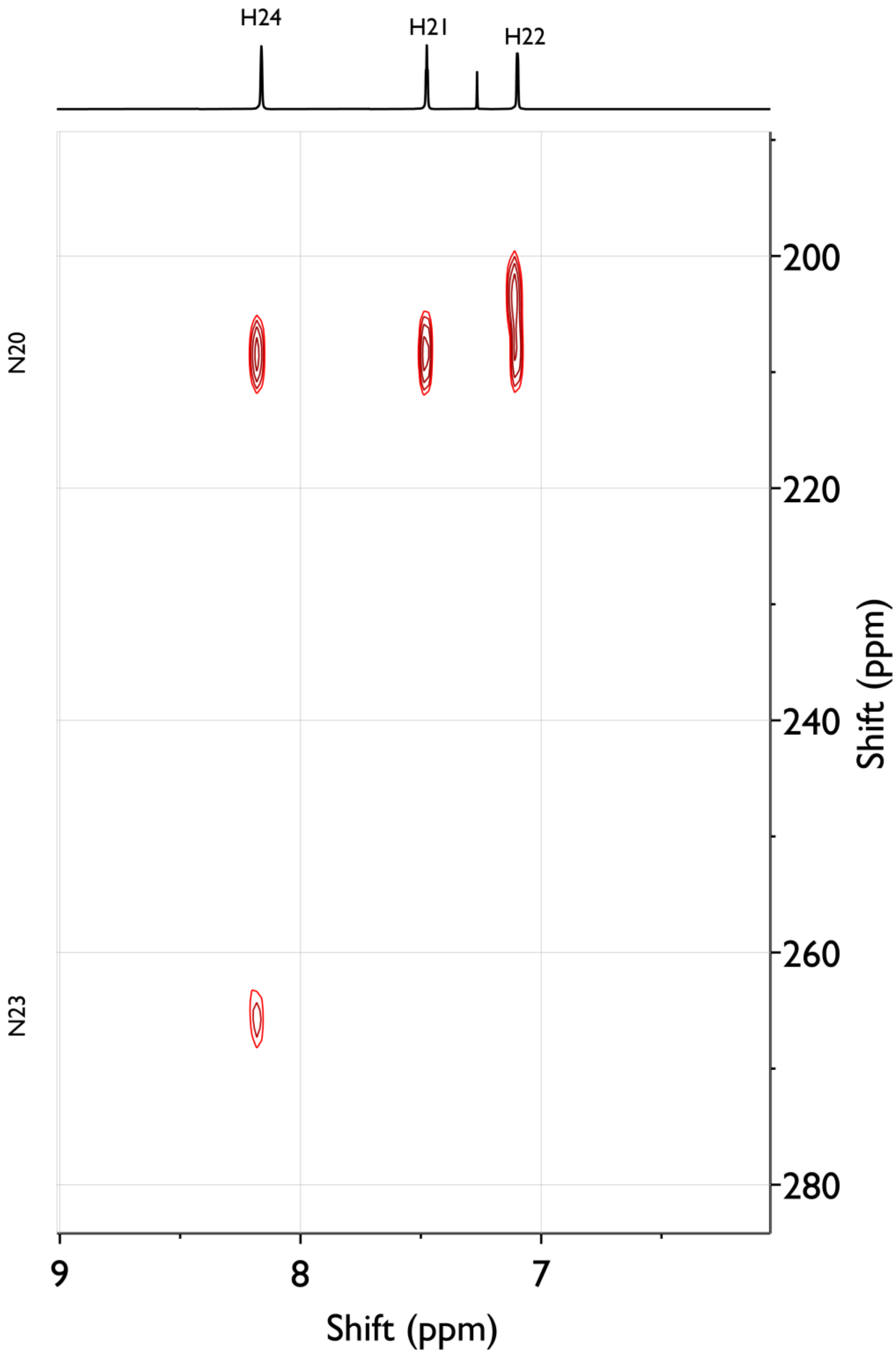


**Figure S90.** $^{15}N$–$^{1}H$ HMBC NMR spectrum of *a* recorded in $CDCl_3$ at 400 MHz, showing long-range $^{15}N$–$^{1}H$ correlations linking the imidazole nitrogen to adjacent proton environments. *$^{15}N$–$^{1}H$ HSQC NMR spectrum does not show any one-bond $^{15}N$–$^{1}H$ correlations as expected from the assigned structure.

**Figure S91.**

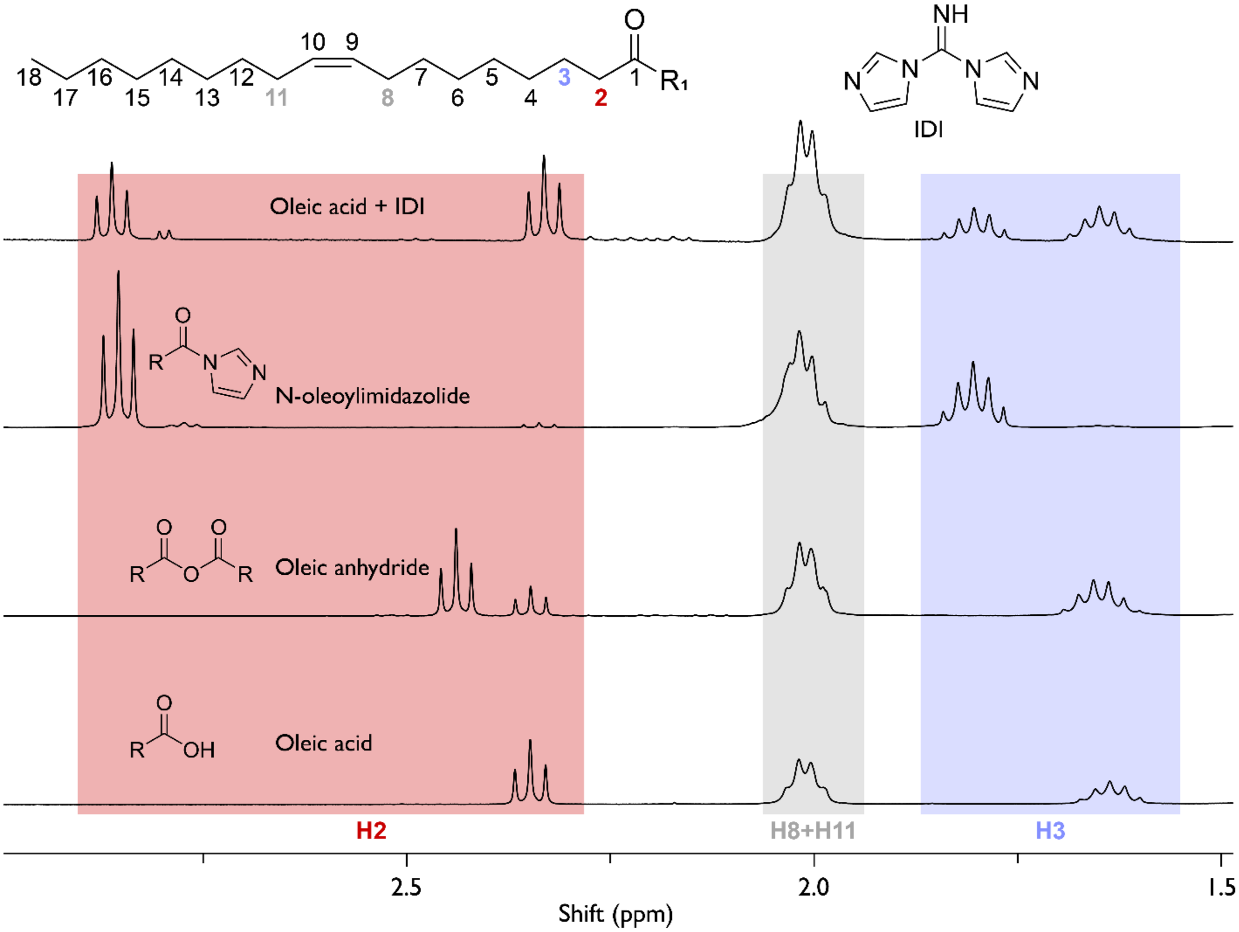


**Figure S91.** [1]H NMR spectra of lipophilic products extracted from oleic acid vesicles incubated with 200 mM IDI at pH 8.0 for 2 h at 18 °C, followed by chloroform extraction, lyophilization, dissolution in $CDCl_3$, and analysis at 400 MHz. The spectrum is plotted alongside relevant reference standards. Diagnostic protons attached to C2 and C3 are highlighted in red and blue, respectively, while protons attached to C8 and C11 are highlighted in grey. The dominant spectral features are consistent with formation of N-oleoylimidazolide as the major identified reaction product.

**Figure S92.**

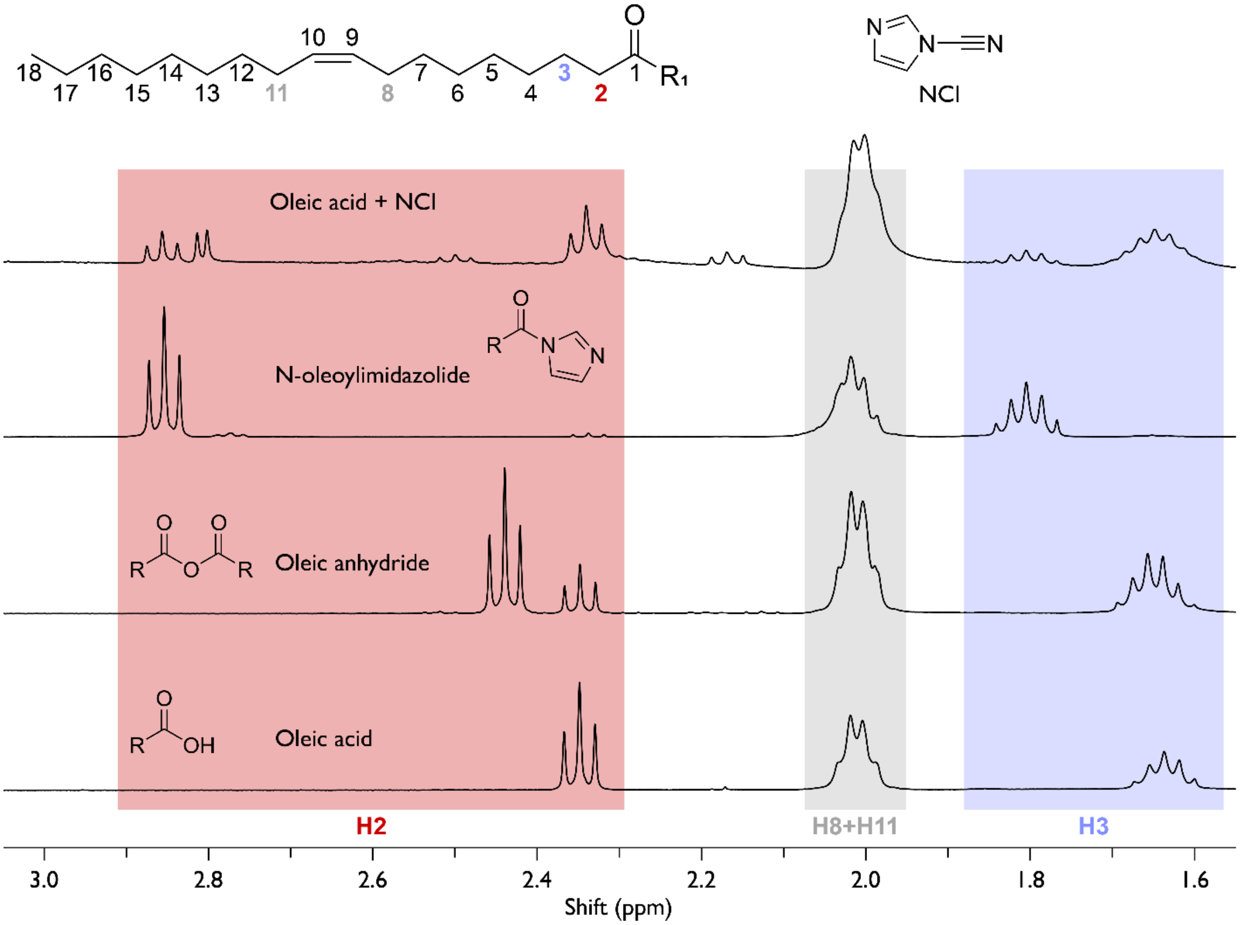


**Figure S92.** $^{1}H$ NMR spectra of lipophilic products extracted from oleic acid vesicles incubated with 200 mM NCI at pH 8.0 for 2 h at 18 °C, followed by chloroform extraction, lyophilization, dissolution in $CDCl_3$, and analysis at 400 MHz. The spectrum is plotted alongside relevant reference standards. Diagnostic protons attached to C2 and C3 are highlighted in red and blue, respectively, while protons attached to C8 and C11 are highlighted in grey. The dominant spectral features are consistent with formation of N-oleoylimidazolide as the major identified reaction product.

**Figure S93.**

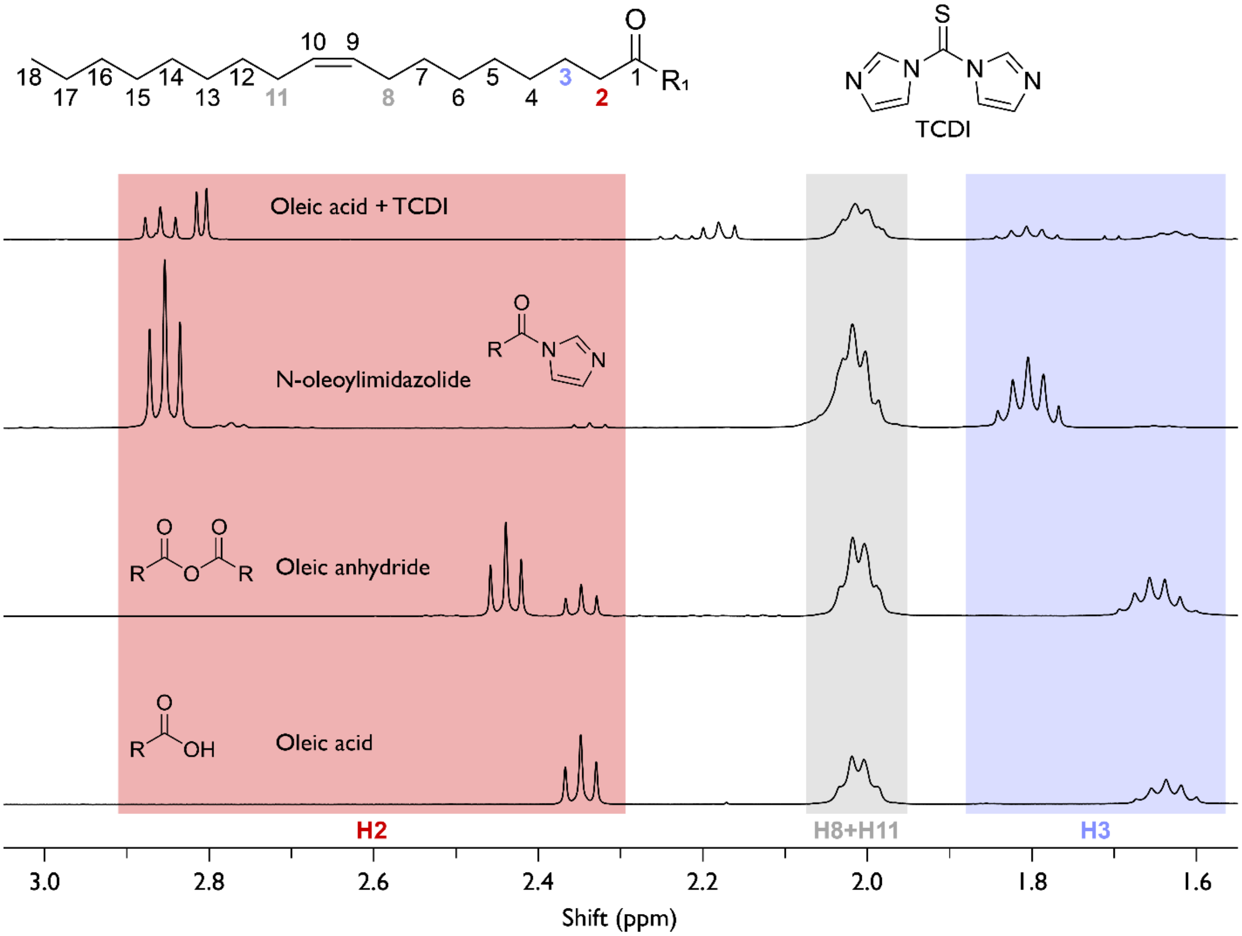


**Figure S93.** $^{1}H$ NMR spectra of lipophilic products extracted from oleic acid vesicles incubated with 200 mM IDI at pH 8.0 for 2 h at 18 °C, followed by chloroform extraction, lyophilization, dissolution in $CDCl_3$, and analysis at 400 MHz. The spectrum is plotted alongside relevant reference standards. Diagnostic protons attached to C2 and C3 are highlighted in red and blue, respectively, while protons attached to C8 and C11 are highlighted in grey. The dominant spectral features are consistent with formation of N-oleoylimidazolide as the major identified reaction product.

**Table S1.**

**Table S1.** Oligonucleotide sequences used in this study.

| **Name** | **Sequence(5′→3′)** |
|---|---|
| U10-Cy3 | UUUUUUUUUU-Cy3 |
| A10-Cy3 | AAAAAAAAAA-Cy3 |
| C10-Cy3 | CCCCCCCCCC-Cy3 |
| (GA)5-Cy3 | GAGAGAGAGA-Cy3 |
| dU10-Cy3 | dUdUdUdUdUdUdUdUdUdU-Cy3 |
| dA10-Cy3 | dAdAdAdAdAdAdAdAdAdA-Cy3 |
| dC10-Cy3 | dCdCdCdCdCdCdCdCdCdC-Cy3 |
| (dGdA)5-Cy3 | dGdAdGdAdGdAdGdAdGdA-Cy3 |
| ATTO550-dT10-ddC | ATTO550-dTdTdTdTdTdTdTdTdTdTddC |
| ATTO550-dA10-ddC | ATTO550-dAdAdAdAdAdAdAdAdAdAddC |
| ATTO550-dC10-ddC | ATTO550-dCdCdCdCdCdCdCdCdCdCddC |
| ATTO550-(dGdA)5-ddC | ATTO550-dGdAdGdAdGdAdGdAdGdAddC |
| RNA(or DNA)-Cy5 (for vesicle imaging) | Cy5-AUAUUGCUACUGGUC |

**Table S2.**

**Table S2.** Observed rates for reactions of nucleotide 5′-monophosphates under IDI activation, determined from $^{31}$P NMR kinetic analysis. Rates ($h^{-1}$) are reported for consumption of the unmodified nucleotide (NMP), formation of the phosphorimidazolide intermediate (NMP-Im), and formation of the 2′/3′-acylated product (Acyl-NMP). Measurements were performed for AMP, CMP, and GMP at pH 6.0 and pH 8.0. Rates were obtained by fitting time-course data to a single-step kinetic model.

| Nucleotide | pH | NMP ($h^{-1}$) | NMP-Im ($h^{-1}$) | Acyl-NMP ($h^{-1}$) |
|---|---|---|---|---|
| **AMP** | 6.0 | 0.6097 | 0.3843 | 10.98 |
| **AMP** | 8.0 | 0.8858 | 0.5566 | 1.150 |
| **CMP** | 6.0 | 0.8876 | 0.8704 | 3.129 |
| **CMP** | 8.0 | 0.2669 | 0.9751 | 0.1250 |
| **GMP** | 6.0 | 1.300 | 1.881 | 0.3900 |
| **GMP** | 8.0 | 0.3957 | 1.728 | 0.3370 |
